# Spatial profiling of longitudinal glioblastoma reveals consistent changes in cellular architecture, post-treatment

**DOI:** 10.1101/2025.01.31.635832

**Authors:** Shoaib Ajaib, Steven Pollock, Gemma Hemmings, Arief Gusnanto, Aruna Chakrabarty, Azzam Ismail, Erica Wilson, Bethany Hunter, Andrew Filby, David McDonald, Asa A. Brockman, Rebecca A. Ihrie, Lucy F. Stead

## Abstract

Glioblastoma (GBM), the most aggressive adult brain cancer, comprises a complex tumour microenvironment (TME) with diverse cellular interactions driving progression and pathobiology. How these spatial patterns and interactions evolve with treatment remains unclear. Here, we apply imaging mass cytometry to analyse protein-level changes in paired pre- and post-treatment GBM samples from five patients. We find a significant post-treatment increase in normal brain cells alongside a reduction in vascular cells. Moreover, despite minimal overall change in cellular diversity, interactions among astrocytes, oligodendrocytes, and vascular cells increase post-treatment, suggesting reorganisation of the TME. The GBM TME cells form spatially organized layers driven by hypoxia pre-treatment, but this influence diminishes post-treatment, giving way to less organised layers with organisation driven by reactive astrocytes and lymphocytes. These findings provide insight into treatment-induced shifts in GBM’s cellular landscape, highlighting aspects of the evolving TME that appear to facilitate recurrence and are, therefore, potential therapeutic targets.

**Key points:**

- Spatial organisation in primary GBM consist of layers driven by the presence of hypoxia
- The layers in recurrent GBM appear are driven more by the presence of reactive astrocytes
- Increased cellular cross-talk in recurrent GBM presents novel therapeutic targets

## Introduction

Isocitrate dehydrogenase (IDH)-wildtype glioblastoma (GBM) is the most common and aggressive form of adult diffuse glioma, with a median survival of ∼15 months ^1^. Standard treatment consists of surgical resection followed by radiation and chemotherapy with temozolomide^2^. However, tumour recurrence is inevitable due to: a) the infiltrative nature of primary GBM, which precludes complete surgical removal; and b) significant intra- and inter-tumour heterogeneity, which enables residual cells to resist chemoradiation and continue proliferating ^3,4^. Characterising how unresected GBM cells respond to treatment can highlight potential mechanisms of treatment resistance that could be additionally targeted with combined therapies.

It is known that IDH-wildtype GBM cells exhibit plasticity across four neoplastic cell states along a proneural to mesenchymal axis^5–7:^ neural progenitor-like (NPC), oligodendrocyte progenitor-like (OPC), astrocyte-like (AC), and mesenchymal-like (MES). However, these neoplastic cells do not function in isolation. In their updated hallmarks of cancer, Hanahan and Weinberg remarked that any understanding of tumours “must encompass the contributions of the tumour microenvironment (TME)’’ ^8^. In GBM the TME comprises a diverse array of tumour cells and also complex network of immune cells, stromal cells, and vascular elements, that play a critical role in GBM progression and treatment resistance, acting as a dynamic ecosystem that influences tumour behaviour and therapeutic response ^9,10^.

To truly understand GBM tumour response to treatment, therefore, requires characterisation at single cell level in ways that incorporate information about interactions with the TME. This is now possible through the use of spatial molecular profiling technologies ^11^. Such approaches have recently been applied to GBM tumours, revealing niches containing specific neoplastic cells and distinct immune-associated programs ^12–14^. These niches have also been shown to organize into structured layers, beyond what is visible via conventional microscopy and histopathology, and are associated with cellular states such as hypoxia ^12^.

These findings describe consistent organizational patterns across GBM tumours suggesting that neoplastic phenotypes are driven by environmental interactions. However, one crucial aspect remains unexplored: how the spatial patterns and interactions within the TME are impacted by treatment to enable some neoplastic cells to survive. To begin to address this, we analysed multiplex imaging mass cytometry (IMC) data ^15^, from five paired pre- and post-treatment IDH-wildtype GBM patient samples, focusing on protein-level changes that reveal alterations in cellular prevalence and states.

## Results

### Identifying and labelling cell types in GBM

To assess the spatial evolution of GBM tumours through treatment, we collected tumour samples from five patients who had undergone surgical resections of both primary and recurrent IDH-wildtype GBM. Each primary tumour developed *de novo*, and all patients received radiation, chemotherapy with temozolomide and had a local recurrence. For patient information see Supplementary Table 1. Three spatially distinct 1mm^2^ regions of interest (ROIs) were selected for each tumour sample, following immunohistochemical staining for key markers of proliferation (Ki67), hypoxia (HIF1A) and immune cells (CD45), to capture intra-tumour heterogeneity and avoid the bias of examining only a single small region (Figure 1A and Supplementary Table 2).

**Figure 1.**
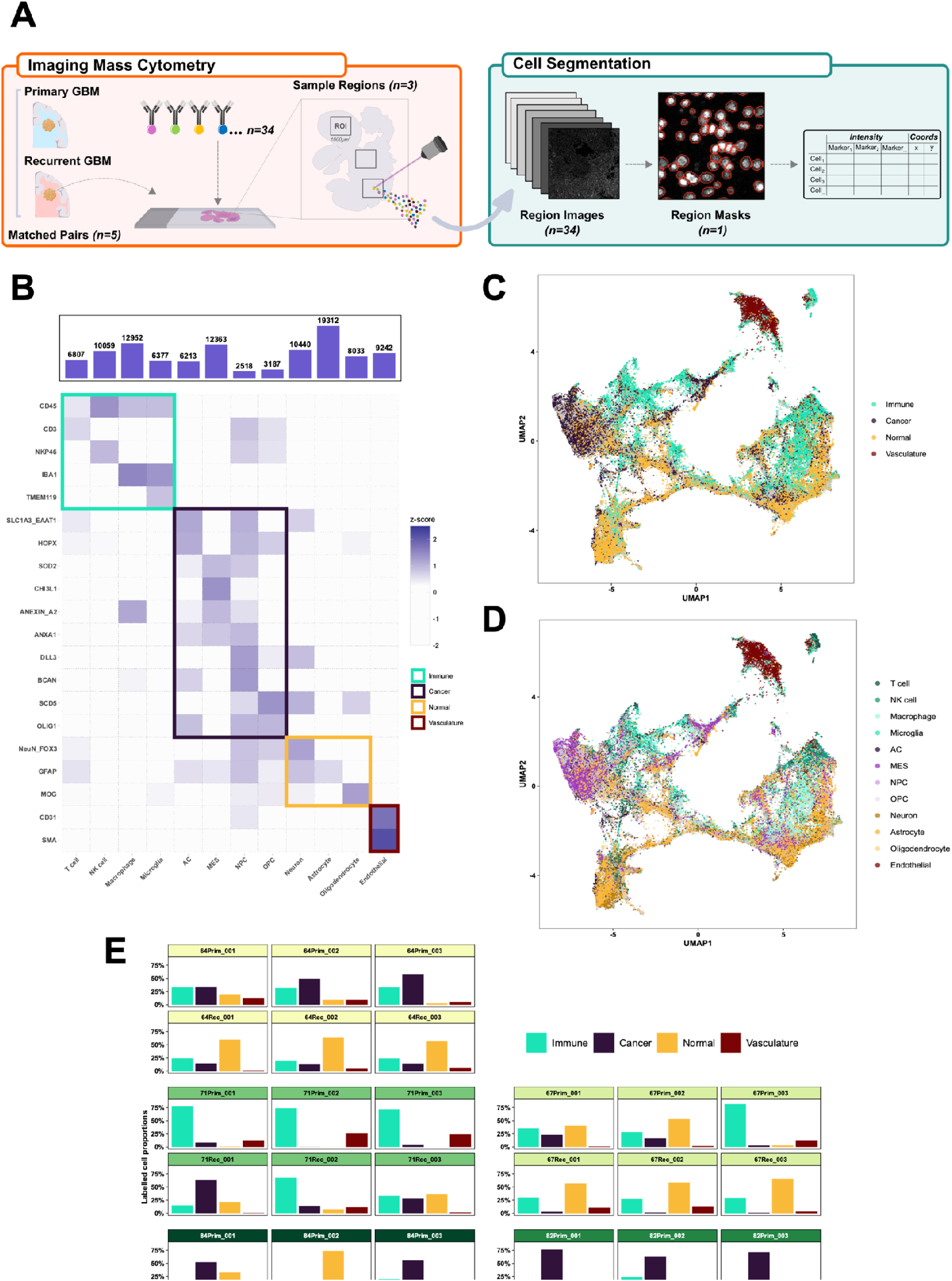
Cell segmentation and phenotyping overview. **A)** Schematic detailing the imaging mass cytometry (IMC) process for one patient sample, including the downstream analysis steps comprising of object segmentation and marker abundance quantification. **B)** Heatmap of protein marker abundances (rows) for each of the labelled cell types (columns). The tile colours denote the scaled (z-score) marker intensities, and the tile highlight colours represent the four main cell categories. **C-D)** UMAP of all (patients and surgeries) cell objects identified following segmentation, batch correction and phenotyping: cells are coloured by cell category (C) and cell type (D). **E)** Proportion of labelled cell categories (columns) across each region of interest (ROI). The facets are grouped by patient/surgery and each of the facet header colours denote an individual patient.

We designed a panel of 34 protein markers to identify GBM-specific cell types (neoplastic, immune, and normal brain cells) along with markers of cell states such as proliferation and hypoxia (see Supplementary Table 3). Using a deep learning-based image segmentation approach, we assigned cell type labels to each segmented object and also subsequently grouped cells into four categories: immune, cancer, normal brain, and vasculature (Figure 1B). Approximately 107,000 cells were labelled across all samples (Figure 1C-D) after applying batch effect correction to account for variability between individual patients and to ensure that expression profiles were comparable (Supplementary Figures 1-2).

A comparison of cell categories across each ROI (Figure 1E) showed surprisingly consistent within-sample distributions, confirming that there is intra-tumour TME heterogeneity but that this is not as significant as inter-tumour TME heterogeneity. ANOVA analysis (see Supplementary Table 4) confirmed that the effect of patient & surgery was significant for all cell types (p < 0.001), indicating considerable inter-tumour heterogeneity. In contrast, intra-tumour heterogeneity, represented by differences across ROIs, was not significant for any cell category suggesting that intra-tumour TME variability is less pronounced compared to inter-tumour heterogeneity. Therefore, we combined the three ROIs per sample, prior to subsequent downstream analyses, to increase the number of cells per sample whilst minimizing sampling bias from specific regions.

### Alterations in cellular prevalence through treatment in GBM

We first assessed how the prevalence of each cell category changed through treatment, between primary and recurrent samples (Figure 2A and Supplementary Table 5). Whilst a reduction in the percentage of both immune and neoplastic cells was observed, from primary to recurrence, only the decrease in vasculature cells was significant (Wilcoxon p = 0.09, 0.06 and 9.88E-03 respectively). The only significant increase was in the proportion of normal brain cells (Wilcoxon p = 4.52E-04).

**Figure 2.**
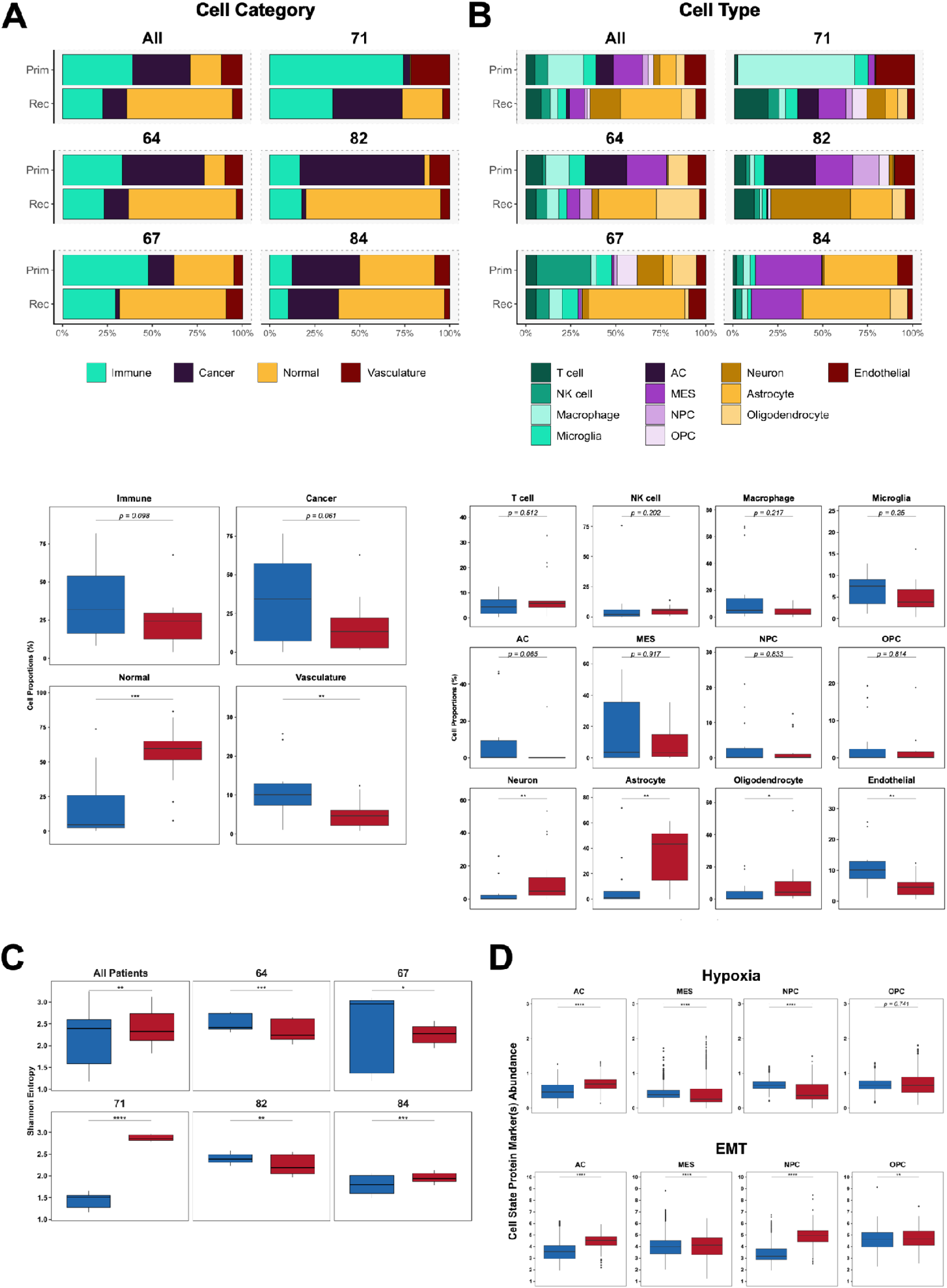
Changes in GBM cell categories and types through treatment. **A)** Top: stacked bar charts showing the labelled cell category prevalences across all patients (top left) and also separately for each individual patient. Bottom: boxplots showing the distribution of each cell category proportion across all patients grouped by primary and recurrent surgeries. **B)** Top: stacked bar charts showing the labelled cell type prevalences across all patients (top left) and also separately for each individual patient. Bottom: boxplots showing the distribution of each cell type proportion across all patients grouped by primary and recurrent surgeries. **C)** Boxplots showing the distribution of Shannon’s entropy values grouped by surgery and split across all patients (top left) and also for each individual patient. **D)** Boxplots, grouped by surgery showing the distribution of protein marker abundance for markers which define hypoxia (top) and the epithelial-to-mesenchymal-transition (bottom). The black horizontal boxplot lines represent the median and the upper and lower box bounds denote the 25^th^ and 75^th^ quantiles, respectively. Astrocyte-like (AC); Mesenchymal-like (MES); neural progenitor-like (NPC); oligodendrocyte progenitor-like (OPC); Epithelial to mesenchymal transition (EMT). Significant p values are represented as: *p < 0.05; **p < 0.01; ***p < 0.001; ****p < 0.0001.

Drilling down into how specific cell types change through treatment showed that no individual immune cell type exhibited significant changes (Figure 2B and Supplementary Table 6). Similarly, no individual cancer cell types altered in a consistent direction, although the astrocyte-like (AC) cancer cell type showed the largest and most consistent decrease (Wilcoxon p = 0.065). All normal brain cell types showed significant increases during treatment, with astrocytes exhibiting a particularly notable increase from primary to recurrence (Wilcoxon p = 3.02E-03), that was consistent across each patient. Of note, astrocytes appear to be the most prevalent normal brain cell type overall, consistent with reports of their high prevalence in both normal brain and the GBM TME ^16,17^.

These changes agree with those from our larger cohort studies where we performed deconvolution from bulk RNAseq, validating our approach ^18,19^.

The limited number of consistent, significant changes in the prevalence of cell category or type over time highlights the variability in immune and neoplastic cell categories, post-treatment, across patients (Figure 2A-B). We, therefore decided to systematically evaluate how cell diversity changes through treatment, both overall and at an individual patient level, to determine if any consistent patterns emerge.

### Alterations in cellular diversity through treatment in GBM

To inspect cellular diversity in our samples, we quantified the Shannon’s entropy (H) for each one (Figure 2C and Supplementary Table 7). A high Shannon’s entropy value indicates a tumour with many different cell types of similar frequency, whereas low entropy suggests that the tumour is dominated by few(er) cell types. This metric thus serves as a good proxy for assessing intra-tumour cellular heterogeneity for each sample, for example pre- and post-treatment.

We found that, overall, Shannon’s entropy significantly decreased from primary to recurrence (Wilcoxon q = 3.93E-03, Figure 2C), suggesting that cell distributions become less diverse, likely owing to certain cell types becoming more dominant within the distribution at recurrence. Linking this back to the results in Figure 2A and B, this appears to be driven by the greater abundance of normal brain cells, and especially astrocytes, in the recurrent tumours. However, analysis of individual patients revealed variability in how cellular heterogeneity changed over time. Two patients (71 and 84) had significantly increased diversity through treatment (Wilcoxon q = 8.45E-17 and 7.10E-04, respectively, Figure 2C). In patient 84, this increase was primarily driven by the appearance of oligodendrocytes at recurrence, which weren’t present in the primary tumour (Figure 2B). Conversely, for patient 71, the increase in entropy was associated with a reduction of dominating macrophages in the primary and presence of a larger neoplastic and normal brain cell fraction at recurrence (Figure 2B).

Given a lack of consistent trends in how treatment affects cell type prevalence or dominance, we proceeded to investigate whether changes in cell state could indicate how treatment shapes cancer cell phenotypes.

### Alterations in neoplastic cellular states through treatment in GBM

The mesenchymal (MES) phenotype in GBM cancer cells is characterised by high proliferative and metastatic potential, often leading to a poorer prognosis compared to proneural subtypes ^20–23^. Moreover, elevated hypoxia and the expression of epithelial-to-mesenchymal transition (EMT) genes, typically involved in neural tube formation or wound healing, have been shown to be closely linked to the MES cell state ^24^.

In our IMC panel we included antibodies against proteins indicating hypoxia (HIF1A) and epithelial to mesenchymal transition (SNAI1 & TGFBeta) to assess the proportion of each of the four identified neoplastic cancer cell types that are in these cellular states, and how they changed through treatment. We found that significantly more AC cancer cells expressed hypoxia markers post-treatment (Wilcoxon p = 4.98E-115), whilst significantly fewer MES and NPC cells did (Wilcoxon p = 9.62E-125 and p = 5.49E-70, respectively) (Figure 2D and Supplementary Table 8).

All four neoplastic cell types had a significantly higher proportion of cells expressing markers of EMT post-treatment, with the largest effect sizes observed in AC and NPC cells (Wilcoxon p = 5.29E-161 and p = 2.32E-183, respectively).

The power of our approach is not just in inspecting paired longitudinal GBM samples at single cell resolution, but also describing how treatment alters the cellular landscape in terms of spatial context. Hence, we moved on to looking at *in situ* cellular interactions, i.e. cells directly adjacent to one another.

### Alterations in cellular interactions through treatment in GBM

We evaluated pairwise cell-cell adjacency, serving as an indicator of cell interaction partners, to assess whether any were significantly more likely (cells are ’interacting’) or less likely (cells are ‘avoiding’) compared to the null hypothesis of spatial randomness (Figure 3A and Supplementary Table 9-10).

**Figure 3.**
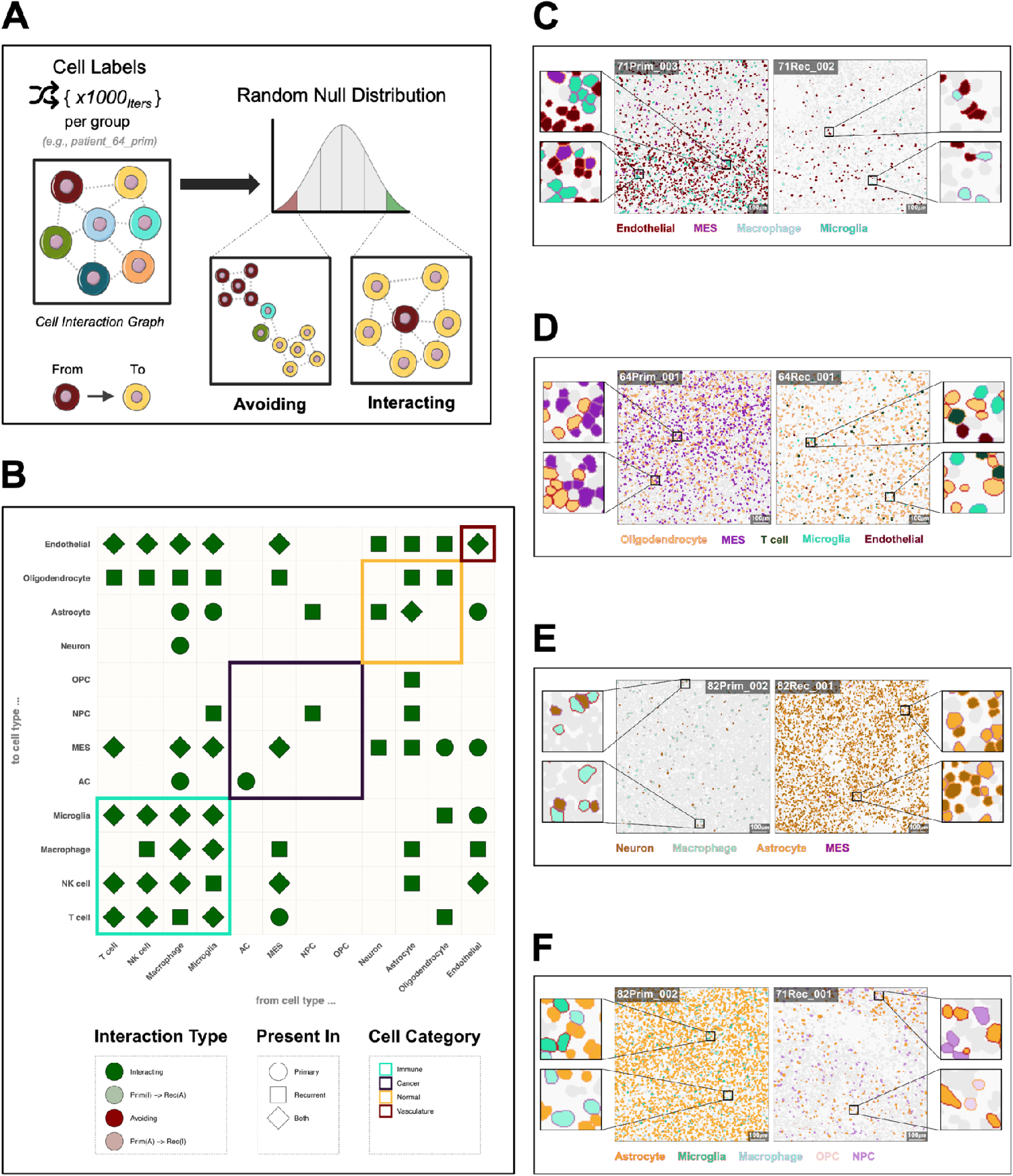
Identifying significant GBM cell-cell interactions through treatment. **A)** Schematic showing the process for testing if cell types interact more or less frequently than random. Each cell-cell interaction (edge) from an interaction graph is counted and averaged across a defined group (e.g., patient, surgery etc.). These average cell-cell interaction are then divided by the number of cells of type A that have at least one neighbour of type B. Finally, each observed cell-cell interaction count is compared against a null distribution that is generated by shuffling the cell-type labels 1000 times (1000 iterations) and counting the interactions between two specific cell types, giving the interaction counts under spatial randomness. Two cell types are “avoiding” when there are the significantly fewer interactions compared to random expectation for a given p values threshold. Conversely, when there are significantly more interactions between the two cell types they are “interacting”. **B)** Dotplot showing the significant (p <0.01) cell-cell interactions which are present across a minimum of three patients. Shape denotes whether a specific cell-cell interaction is significant across either primary, recurrent or both surgeries. Point colours denote the type of significant interaction, i.e., interacting/avoiding and also cases where the interaction type changes through surgery. The tile highlights denote the cell category of each cell type. **C-F)** Representative IMC images showing single-cell segmentation masks coloured by the corresponding cell type labels for primary (left) and recurrent (right) surgery regions of interest.

We performed this analysis on the primary and recurrent samples separately to see which significant findings were timepoint dependent. Many cell types predominantly interacted with themselves in the primary tumours (Figure 3B). This is in keeping with previous spatial analysis of GBM that used a ‘spot-based’ technology, that is not resolved at the single-cell level but rather aggregated over a small defined area (spot), which found that signal from the majority of spots seemed to emanate from a single cell type ^12^. Our expansion to recurrent samples shows that these ‘self’ interactions remained consistent through treatment (Figure 3B). Two additional, clear observations from our results are that there are no cells significantly avoiding one another, and there are many more recurrence-specific, significant cell-cell interactions than primary-specific ones (27 versus 10). Hence, despite finding an overall reduction in cell diversity at recurrence (Figure 2C), there are more interactions between differing cell types, suggesting that these are non-random and, thus, phenotypically important.

#### Neoplastic cells

Amongst the GBM cancer cell types, MES cells formed the highest number of significant interactions with other cell types. MES interactions with immune cells remained consistent between paired samples but interactions with normal brain cells were increased at recurrence.

#### Vasculature

Despite decreasing through treatment (Figure 2A), endothelial cells still formed significant interactions at both time points (Figure 3C). Unique to the primary tumours were significant interactions from the endothelial cells to the microglia (permutation test, p = 9.99E-04) and MES cancer cells (permutation test, p = 9.99E-04). MES cells interacting with myeloid lineage cells (e.g., macrophages and microglia) have been shown to lead to a highly proliferative state, increasing angiogenesis and contributing to a more invasive phenotype, which may explain these findings in the primary tumour ^25^. Moreover, these interactions have also been shown to induce chemoresistance in GBM, which has the potential to be addressed therapeutically ^26^.

The interactions from the endothelial cells to the macrophages were particular to recurrent tumours (permutation test, p = 9.99E-04). These findings could be visualised in the IMC data (Figure 3C) where a clear reduction in endothelial cells over time coincided with changes in the cells interacting with the remaining vasculature. It has previously been shown that bone derived macrophages populate a GBM tumour post-treatment, via the vascular system, which may explain this result and further indicate that therapies which hijack this infiltration could be effective for preventing or prolonging GBM recurrence ^27^. Interactions from all normal brain cells to endothelial cells were also specific to the recurrent tumour. The post-treatment increase in normal brain cell abundance within the resected tissue (Figure 2A) may reflect the brain’s wound healing response, with neuronal and glial cells re-populating the void left by surgery ^28,29^.

#### Normal Brain Cells

Significant interactions from and to oligodendrocytes almost universally occurred in the recurrent tumours, barring those from oligodendrocytes to MES cancer cells, which were primary-specific (Figure 3B). This could be observed in the IMC visualisations (Figure 3D). The prevalence of oligodendrocytes increases from primary to recurrent (Figure 2B) suggesting that this population did not simply expand *in situ* but rather infiltrated the recurrent TME. In GBM, oligodendrocyte lineage cells have commonly been reported to reside at tumour border niches including the invasion front/resection border where they co-localise with macrophages/microglia ^30^. Moreover, oligodendrocytes have been shown to support GBM tumorigenicity and migration by promoting angiogenesis in GBM ^31,32^. We also found evidence supporting the model of interactions, as microglia and endothelial cells were significantly interacting with oligodendrocytes at recurrence (Figure 3D).

Of all the normal brain cells, astrocytes were found to significantly interact most frequently and significantly with the cancer cells, though this is mostly specifically at recurrence. In fact, aside from “self” interactions which were consistent through treatment, normal astrocytes only formed significant interactions during recurrence.

Crosstalk between microglia and macrophages is known to induce reactive astrocyte phenotypes, which are crucial for the brain’s wound healing process - a key aspect in GBM ^33,34^. Moreover, the MES phenotypes, as described by Wang et al. and Neftel et al., have been shown to overlap significantly with the presence of reactive astrocytes, indicating that these cells may migrate to injury sites after resection as part of the healing process ^28,29^. In our samples we found significant interactions between normal neurons and astrocytes, suggesting the activation of cellular programmes that could restore normal tissue function (Figure 3E and F).

### Alterations in cellular neighbourhoods through treatment in GBM

Cell interactions within the GBM TME are heavily influenced by the spatial context, as GBM tumours consist of distinct anatomical regions ^35^. To generalise groups of interacting cell types we defined cellular neighbourhoods (CNs) using a nearest neighbours approach (Figure 4A). This method defined 12 distinct cellular neighbourhoods that provided a different level of structure from that observed based just on individual cells (as exemplified in Figure 4B). As expected, owing to the fact that each cell significantly associates with itself in both the primary and recurrent tumours (Figure 3B) we found that most cell neighbourhoods are dominated by a specific type (Figure 4C). CNs capture multiple cells in close proximity (Figure 4A) so are akin to the information captured by spot-based spatial technologies such as the 10X Visium platform. Our finding of dominance of a given cell type in each defined CN agrees with Greenwald et al’s recently published results from application of the Visium platform to primary GBM samples ^12^. Extending these results using imaging mass cytometry, which provides single cell resolution, we can further see that this dominance rarely equates to more than 50% of the cell types in a given CN, meaning there is clear ad-mixture and heterogeneity in interacting cells even when signal fomr one type predominates (Figure 4C).

**Figure 4.**
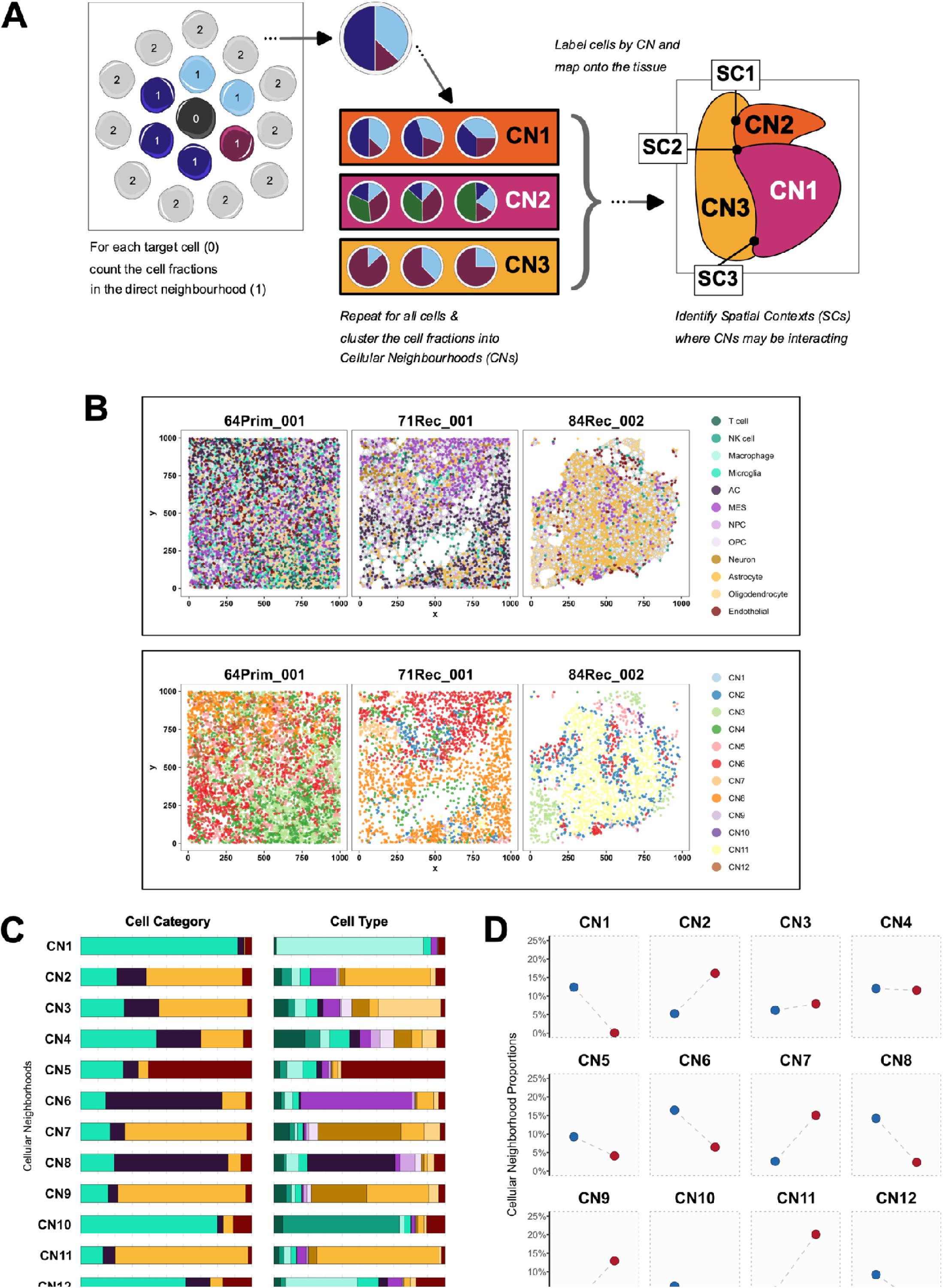
Identifying distinct cellular neighbourhoods present across GBM samples through treatment. **A)** Schematic showing the process of identifying cellular neighbourhoods (CNs): each cells direct neighbourhood (as defined by an interaction graph) cell fraction is aggregated and clustered across each patient/surgery. The resulting CN cluster labels are then mapped to each cell object. The spatial contexts (SCs) are then defined as locations where the most dominant CNs interactions are also interacting. **B)** Plots showing the single-cell spatial locations of three representative patient/surgery sample regions of interest visualised as nodes on a 2-dimensional plane, with cell-cell interactions shown in the form of undirected edges between nodes (top). The nodes are coloured according to the cell type label (top) and also by the cellular neighbourhoods they belong to (bottom). **C)** Stacked bar charts showing the proportion of each cell category (left) and cell type (right) that is present across each CN (rows). **D)** Dot plots showing the relative proportion of cells in each of the CNs (facets) across each surgery type.

Greenwald et al proceeded to cluster their spot-based gene expression profiles into 16 ‘metaprograms’ (MPs). Our CNs map to these MPs (Table 1 and Supplementary Table 11), though with some differences due to the level of cellular resolution and the differences in dimensionality and modality between the two studies. Specifically, Visium spots capture signals from 1-35 cells, so some MPs result from more than just nearest neighbours; and MPs are derived from gene expression (typically 7000 parameters our CNs derive from protein expression (34 parameters). It is worth noting that we aligned both CN4 (T-cell dominated) and CN10 (NK cell dominated) with the T-cell MP, owing to the functional similarities between T- and NK-cells ^36^.

**Table 1.**
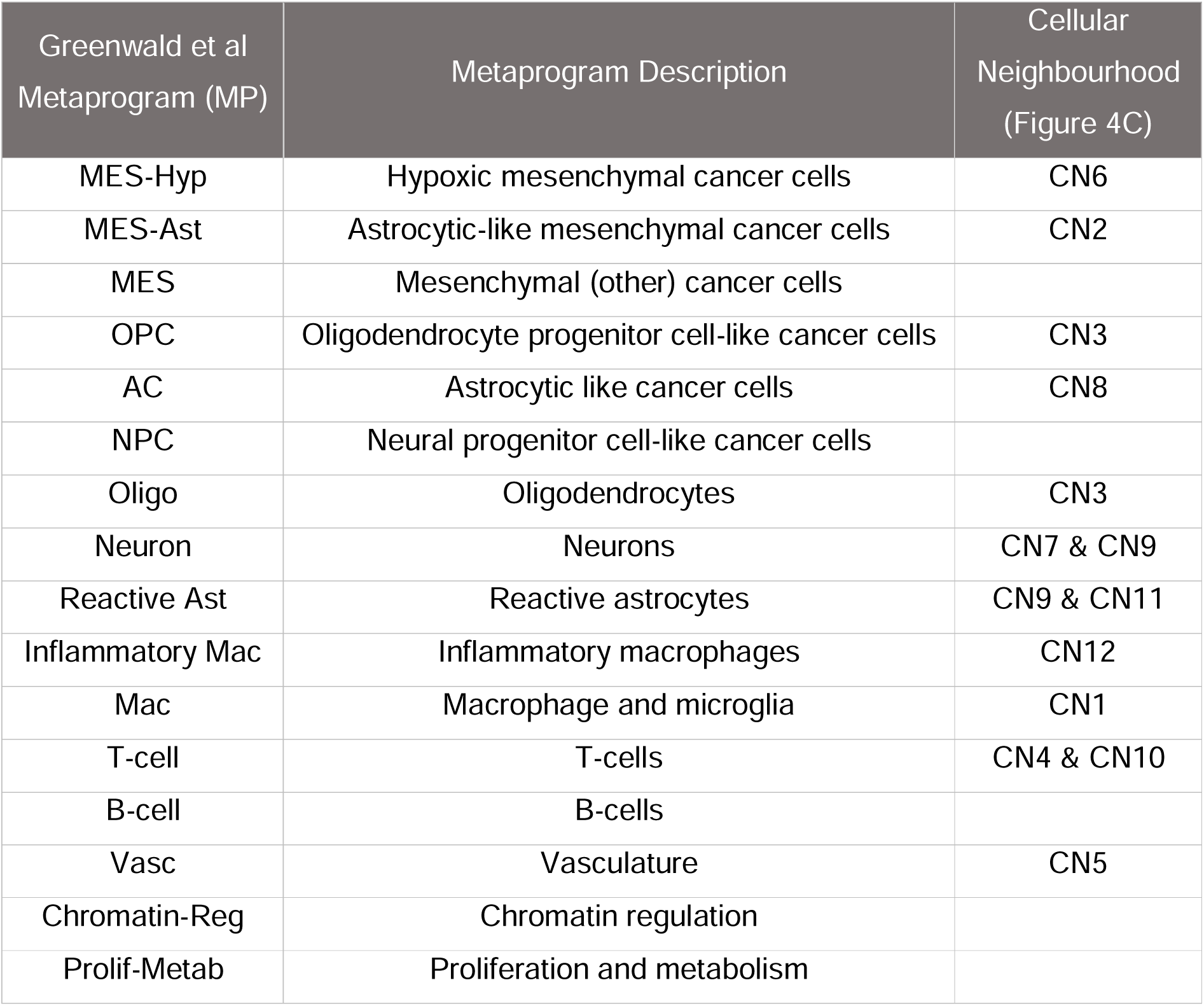
Mapping of previously defined spatial GBM metaprograms to the cellular neighbourhoods defined in this study.

Having aligned with previous findings from GBM tumours at a single time point, we wished to see how the prevalence of CNs change over time. We see that certain CNs increased in abundance from primary to recurrenct tumours, and others decrease (Figure 4D). Primary samples were enriched in neighbourhoods that included immune cells, particularly macrophages (CN1 and CN12) and lymphocytes (CN10), vasculature (CN5), hypoxic MES (CN6) and AC cancer cells (CN8). In contrast, the recurrent surgery samples were enriched in neighbourhoods dominated by normal brain cells: astrocytes (CN9 and CN11); neurons (CN7); and oligodendrocytes (CN3). Interestingly, we found that whilst hypoxic mesenchymal-driven CN6 decreased, astrocytic like mesenchymal-driven CN2 was increased from primary to recurrence.

Ultimately these results reconfirm what was seen when looking at cell type or category prevalences in isolation (Figure 2) i.e. that immune cell-driven (CN1, CN10 and CN12), vascular-cell driven (CN5) and cancer cell-driven (CN6 and CN8) neighbourhoods decreased from primary to recurrence, whereas normal brain cell driven (CN2, CN3, CN7, CN9 and CN11) neighbourhoods increased. CN4, which was dominated by T-cells, changed least in prevalence over time.

Greenwald et al’s seminal finding was that, in some primary GBMs, MPs form organised layers that result in a global tumour architecture, which is seemingly driven by the presence of hypoxic niches. We, therefore, proceeded to investigate whether this organisation was evident in our primary samples and whether it was maintained post-treatment ^12^.

### Alterations in spatial organisation through treatment in GBM

To better understand higher order structuring of our cellular neighbourhoods, we classified spatial contexts (SCs); locations where distinct cellular neighbourhoods were found to consistently interact (Figure 4A). When considering the most dominant CN interactions present across primary and recurrent surgeries our results reproduce similar ordered layers to those reported in Greenwald et.al ^12^. However, the prevalence and importance of states which make up the layers differs greatly through treatment, as revealed by the structure and parameters of the calculated CN interaction networks (Figure 5).

**Figure 5.**
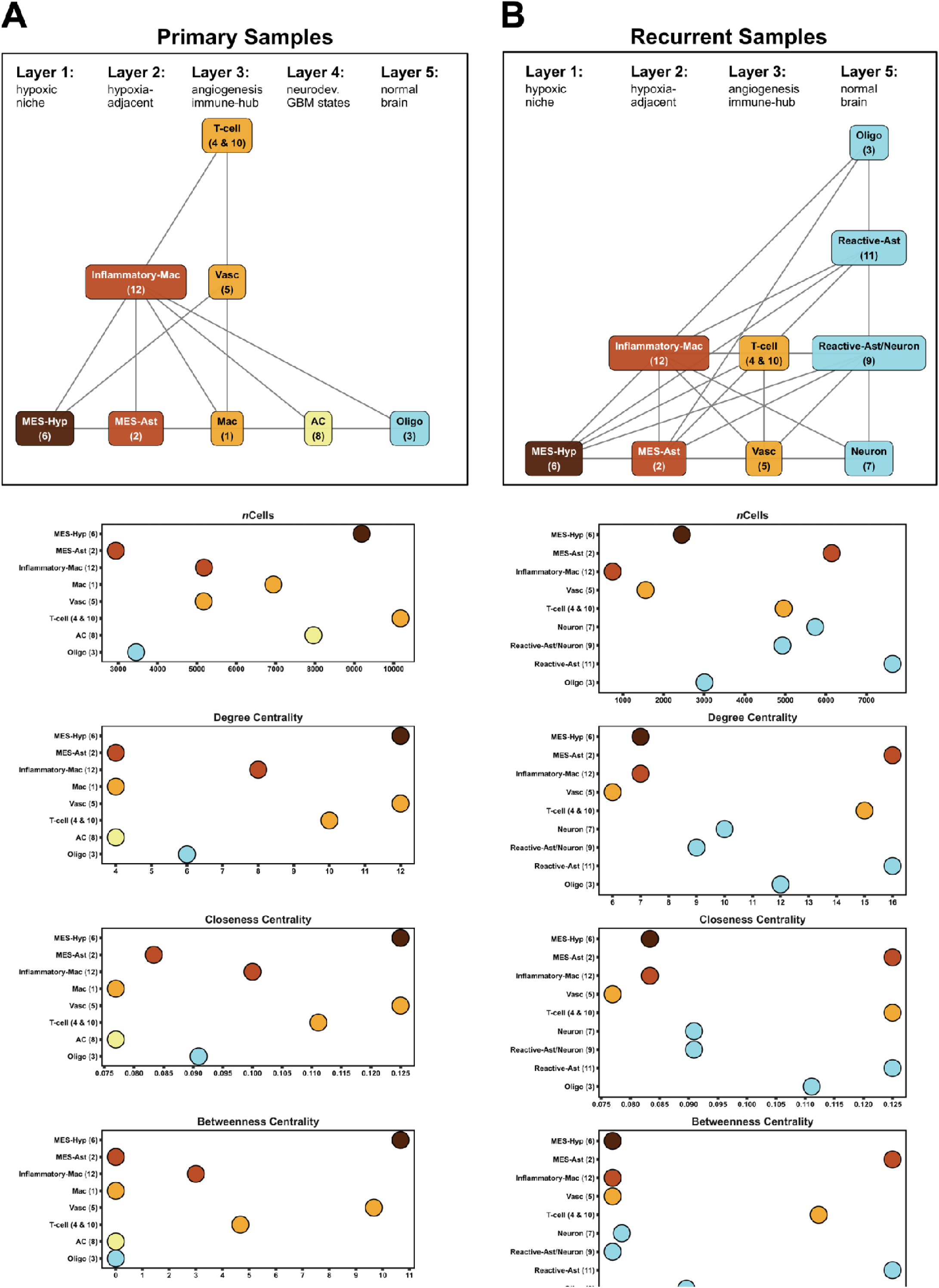
Spatial organisation of cellular neighbourhoods across surgeries. A-B) Top: network graphs with nodes labelled according to the cell metaprograms identified in Greenwald et.al, 2024 along with their corresponding cellular neighbourhoods (shown in the brackets below). The edges represent the most dominant interactions present across primary surgery (A) and recurrent surgery (B) regions of interest (ROI), respectively. Bottom: dot plots showing the number of cells present across each surgery specific cellular neighbourhood and also three network-specific centrality measures: degree; closeness and betweenness. Both network graphs and their corresponding metrics are coloured and ordered according to the structured GBM spatial layers described in Greenwald et.al, 2024.

In the primary samples the most influential and prevalent cellular neighbourhoods were those characterised by layers 1 and layers 3, which denote the hypoxic/necrotic core niche and the angiogenesis-immune hub, respectively (Figure 5A). These layers were comprised of CNs with high network centrality scores across all three measures (degree, closeness and betweenness), indicating their importance for communication between other neighbourhoods and layers. This concurs with previous findings suggesting that hypoxia potentially drives the presence of the organised layers owing to phenotypic consequences of reduced oxygen, especially at the tumour core ^12^.

Conversely, in recurrent samples there were many more significant interactions between CNs in different layers (Figure 5B) in agreement with our findings from pairwise cellular interaction analysis (Figure 3B). Additionally, the most influential and prevalent cellular neighbourhoods in recurrent samples were mostly in layers 2 and 5, which represented the hypoxia-adjacent and normal infiltrative brain layers (Figure 5B). This suggests a reduced global structure with less well organised layers, potentially owing to a reduction in the presence of hypoxic niches in recurrent versus primary GBM.

Worth noting is that T-cell dominated CN4, which remained the most stably prevalent between primary and recurrent samples (Figure 4D), in combination with CN10 (together these CNs align to the previously denoted T-cell MP: Table 1) maintain high network parameters in both primary and recurrent GBMs (Figure 5), implying they are important in driving spatial contexts both pre- and post-treatment.

## Discussion

We note that this was a small study (n=5), and technical limitations of protein-based expression profiling confined the number of markers that could be used to assign cell types (n=34). This meant that some cells (e.g. B-cells) were not included, and it also reduced our ability to definitively discern between neoplastic cell types and those of the normal brain, which they closely mirror ^6^. However, the benefit of protein markers is that they are less stochastically expressed than gene expression markers, and that dropout of signal for these parameters (i.e. false negatives) is substantially less than in single-cell sequencing approaches.

Cumulatively our results reveal an influx of normal brain cells into the GBM microenvironment post-treatment, with a concurrent reduction in vascular cells (Figure 2A). The latter has been shown previously and is an expected result given that surgery aims to debulk the core of the primary tumour, which is known to be highly vascularised ^37,38^. However, the reduction of endothelial cells in recurrent GBM also suggests a reduced functional reliance on vasculature which may explain the failure of angiogenic treatments such as bevacizumab (branded as Avastin) at clinical trials ^39^.

The increased presence of oligodendrocytes at recurrence has been shown in several large cohort studies of paired GBM tumours, which deconvoluted cellular signals from bulk RNAseq, and single cell analysis of pre- and post-treatment GBMs ^18,40,41^. Herein we confirm this finding (Figure 2B) but additionally show that these cells integrate into the GBM TME, as the number of significant interactions between oligodendrocytes and other cells increases at recurrence (Figure 3B).

Oligodendrocytes play a crucial role in maintaining cerebral homeostasis by helping to regulate neuronal activity via axon myelination ^30^. We used the myelin oligodendrocyte glycoprotein (MOG) marker to characterise oligodendrocytes (Figure 1B) meaning that our results suggest that it is myelinating oligodendrocytes that increased in number and are integrating into the tissue, suggesting a potential functional role for myelination within the GBM TME post-treatment. Interestingly, oligodendrocyte integration and additional interactions that we identified in recurrent tumours occur primarily with non-neoplastic cells. In terms of cancer cells, the recurrence-specific interactions containing oligodendrocytes are restricted to MES neoplastic cells (Figure 3B). Oligodendrocytes have been shown to upregulate the invasive capacity of GBM cancer cells via Angiopoietin-2 signalling, and MES are the most invasive neoplastic subtype ^31,42^.

We found that all neoplastic GBM cells showed an increase in markers of epithelial to mesenchymal transition (EMT) at recurrence (Figure 2D) and that the cellular neighbourhood dominated by oligodendrocytes (CN3) had higher closeness and degree centrality at recurrence (Figure 5), meaning a heightened proximity and this ability to interact with all other CNs. We propose that the role of oligodendrocytes in driving post-treatment recovery of GBM is worthy of further exploration.

The largest increase of normal brain cells within the recurrent GBM TME is observed for astrocytes (Figure 2B), and these also have the highest number of recurrence specific interactions, particularly those including neoplastic cells (Figure 3B). Within the healthy brain parenchyma, astrocytes are crucial for neuronal cell homeostasis and also help drive the brain’s injury response by acquiring a reactive phenotype. Consistent with this role, cellular neighbourhoods that map to the previously defined reactive astrocytic metaprogramme (CN9 and CN11) are increased, alongside that mapping to the astrocytic mesenchymal metaprogramme (CN2), at recurrence (Figure 4D and Table 1). Astrocytes also exhibit resistance to apoptosis triggered by death receptors during inflammation, such as apoptosis antigen 1 and TNF-related apoptosis-inducing ligands (FAS, TRAIL), indicating their resilience under inflammatory conditions. Together, this suggests a phenotypic response within the (infiltrating) astrocytic population of the TME that could serve to protect neoplastic cells.

Previous research has indicated that there is a shift towards a more mesenchymal state in bulk tumours at recurrence ^18^. The advent of single cell analysis enabled this to be inspected at cellular resolution showing that, whilst some GBMs do show an increase in MES cancer cells post-treatment, this is not universal. Other tumours instead show an increase in the more proneural (OPC and NPC) cells at recurrence ^43,44^. The results herein agree that there is not a significant, consistent change in a specific type of cancer cell at recurrence but do show a universal increase in EMT markers in all the neoplastic cell types (Figure 2D). This potentially explains the shift to mesenchymal expression signatures observed from bulk tumour profiling ^18^.

AC-like cancer cells appeared to reduce the most consistently at recurrence, in keeping with our previous findings ^19^. However, those AC-like cells that remained had elevated levels protein indicating hypoxia (Figure 2B and D). In contrast, hypoxia markers within the MES and NPC cell populations decreased. Hypoxia can induce a reactive astrocyte phenotype within the tumour microenvironment, which may extend to AC-like cancer cells, potentially even promoting plastic conversion to this neoplastic subtype ^45,46^.

Overall, we found that changes in cellular diversity between primary and recurrent GBM are not consistent (Figure 2C), suggesting that there is a maintenance of cellular heterogeneity but with greater interaction between differing cell types post-treatment (greater admixture, Figure 3 and Figure 4). A recent spatial profiling study of primary GBM tumours by Greenwald et al. concluded that hypoxia drives organisation of a GBM architecture, composed of layers ^12^. Our findings concur with theirs for primary GBM tumours but expand upon these results to reveal that this layering is less structured post-treatment (Figure 5). The decrease in CN6, which maps to their hypoxic MES cancer cell metaprogramme (Table 1), but increase in CN2, which maps to their astrocytic MES cancer cell metaprogramme (Table 1), at recurrence suggests that an overall reduction in hypoxia post-treatment, could drive this increased disorder. The influx and integration of normal brain cells in the GBM TME at recurrence corresponds with these cells becoming much more influential in terms of the interaction between cellular layers, particularly CN11 which map to the reactive astrocyte metaprogramme of Greenwald et al. (Figure 5B).

Whilst we see no difference in the abundance of lymphocytes (Figure 2B) from primary to recurrent GBM, we do find that the cellular neighbourhoods (CN4 and CN10) that map to the T-cell metaprogramme (Table 1) become much more influential in the recurrent GBM CN interaction networks (Figure 5B). T-cells and, specifically, tertiary lymphoid structures (cellular regions enriched in lymphocytes, akin to niches captured by CN4 and CN10) have been shown to increase in subsets of paired primary and recurrent GBM ^41,47^, which has renewed interest in understanding if and how this could impact immunotherapies for these tumours. In support of this, a study into GBM MES cancer cell states showed that activated T-cells associate specifically with astrocytic MES; and it is this subtype that we find to increase in recurrent tumours ^20^.

This study has revealed some prominent changes in the cellular landscape of GBM, post-treatment, that offer novel insight into the importance of specific interactions between GBM cancer cells and the TME during tumour survival and regrowth. Further work to characterise these interactions could lead to therapeutic targets to overcome resistance to standard treatment in GBM, or for treating recurrent tumours.

## Materials & Methods

The data acquisition for these samples, and their initial processing, are described in our previous publication where we used them to validate our bulk RNAseq deconvolution tool ^48^.

### Ethics Statement

Tumour samples used in this study were obtained from patients at the Walton Centre, UK, that provided informed consent in writing for the use of their tissue in research. The inclusion of these samples in this project was following approval by the UK National Health Service’s Research Ethics Service Committee South Central - Oxford A (Research Ethics Code: 13/SC/0509).

### Patients & Samples

Tumour samples from five patients were used in this study (Supplementary Table 1). All tumours were de novo primary IDHwt GBM that had been stored in formalin-fixed, paraffin-embedded blocks, and their matched locally recurrent tumour samples. All tumours were obtained following an initial debulking surgery and treatment with radiation and Temozolomide chemotherapy. Two matched (primary and recurrent surgeries) tumour cores were punched for each patient and from each respective core, three 1000µm^2^ regions with varying tumour localization were selected based on the expression of histochemical markers denoting areas of high Immune infiltration (CD45+); hypoxia (HIF1A+) and proliferation (Ki67+). In total 30 regions of interest were acquired (Supplementary Table 2 and Supplementary Figure 3).

### Antibody Conjugation & Validation

An antibody panel of 34 proteins was designed, including markers of immune, normal, cancer (neoplastic) and vasculature cells, in addition to markers for various cellular states such as proliferation, hypoxia and quiescence (Supplementary Table 3). The neoplastic GBM and immune cell markers were selected based on single-cell analysis of GBM samples in addition to literature searches and manufacturer websites, as previously described ^48^.

Antibodies were selected according to a specific order and using the following criteria: 1.) available in pre-conjugated format for IMC and previously used in IMC of GBM or normal brain; 2.) previously used in IMC of GBM or normal brain via bespoke conjugation; available in carrier free format and had been validated for use in IHC or ICC in brain or GBM; 3.) available in carrier free format. A set of panel-wide control tissues was determined: spleen, brain, tonsil, prostate, bone marrow, skin, and uterus. Control tissue samples from at least two individuals were amalgamated into a multi-tissue formalin fixed, paraffin embedded block. Multi-tissue block sections were used in IHC validation and testing of three antibody concentrations at, above and below those recommended by the manufacturer. The chosen antibody concentrations and control tissue(s) relevant to each antibody are in Supplementary Table 3.

Antibody conjugation, staining and IMC took place at the Flow Cytometry Core Facility at Newcastle University. Conjugation was performed using MaxPar metal labelling kits using X8 polymer according to standard manufacturers protocols (with the exception of Gd157 which was obtained by Trace Sciences International and was diluted to 0.1M prior to use with MaxPar reagents). Conjugations were validated by capture on Thermo AbC beads prior to acquisition on a Helios mass cytometer.

### Preparation & Staining

5µm sections, taken consecutively from the same blocks that underwent bulk RNA sequencing (antibody conjugation & validation section), were stained with a cocktail of all 33 conjugated antibodies after dewaxing (Xylene) and HIER antigen retrieval in Tris-EDTA (pH9) with 0.5% Tween 20. Sections were incubated for 30 minutes in 0.3 µM iridium to counterstain the nuclei prior to air drying. Images were generated on the Hyperion Tissue Imaging cytometer by ablation of the ROI at a 200Hz frequency with a 1-micron diameter laser. The raw MCD files were created and exported from MCD Viewer software (Fluidigm).

### IMC Image Processing

All image processing and downstream analysis steps were performed using the R statistical software package (>= version 4.3.0) and Python (version 3.11.3) ^49^. Further details of packages and specific versions are provided in the sections below. The name of each statistical test used, and level of significance achieved, is included within the results where the finding from each hypothesis test is confirmed. All plots were generated using ggplot2 (version 3.5.1) ^50^.

### Cell Segmentation

The Steinbock ^51^ python package (version 0.13.5) and accompanying command line tool was used to convert the raw MCD files obtained from the MCD Viewer software into multi-channel tiff images stacks using the “preprocess” command with default settings.

The multi-channel tiff images (n = 30) were processed using a custom python script where each multi-channel image stack was read using the tifffile (v2023.4.12) module and the signals from specific nuclear (Ir191, Ir193) and cytoplastic channels (Sm149, Eu153, Dy164, Yb171) were combined to generate single RGB images using pandas (v2.0.3) and numpy (v1.24.0). Each combined RGB image was cropped into smaller, random-sized sections (100µm x 100µm) and saved as a tiff image to be used as training data for deep-learning-based cell segmentation. The Cellpose (version 2.0), human-in-the-loop training method was used to segment each of the 30 ROIs as previously described ^52^. Briefly, all of the 100×100µm RGB random crops were locally loaded into the Cellpose graphical user interface and initially segmented with the pre-trained cytoplasm model (“cyto2”), independently using the both the cytoplasmic and nuclear channels defined above. The resulting segmentation masks were then manually refined in an iterative fashion where the base model was updated after re-annotating each ROI, and the mean pixel diameter was automatically adjusted in order to account for the differences in cell shapes/sizes present across all samples. The final refined model was applied to segment the full ROI’s (1000µm^2^) using the “cellpose_train” command with default settings and the segmentation masks were saved as tiff files.

### Cell Quantification

The “measure intensities” function from Steinbock was used to the extract signal intensities for each of the objects identified in the segmentation process. The pixels belonging to an object were aggregated by taking the mean signal intensities across each of the channels. Spatial object properties such as the area, centroid, eccentricity and the major/minor axis lengths were calculated for each object using the “measure regionprops” function from Steinbock (Supplementary Figure 4). Following quantification, the single-cell objects was read into R using “read_steinbock” from the imcRtools R package (version 1.2.3) and the expression counts were transformed using an inverse hyperbolic sine (asinh) function (cofactor = 5).

### Spillover Compensation

Spillover compensation was performed to account for any overlapping signals present across the image channels as previously described ^53^. Briefly, an agarose-coated slide was generated, and an array of trypan blue dots was spotted on the slide (0.3 µl each). For each antibody used, 0.2 µl of the same antibody conjugate was spotted onto one trypan blue spot. For each RNA probe, 0.3 µl of the 10 µM stock of the metal-conjugated probe was spotted onto a trypan blue spot. For each spot, 10 lines of 200 pixels were acquired and named according to the metal tag to be compensated. The resulting .txt files were used to generate a spillover matrix using the CATALYST R package(version 1.20.1) ^54^. This spillover matrix was also exported as a csv file and used to correct both the images and single-cell data generated following cell-segmentation and single cell quantification.

### Batch Correction

All spillover-compensated, single-cell objects were integrated to correct for unwanted sources of variation present across each patient using the harmony algorithm (Supplementary Figures 1-2) ^55^. Briefly, we performed PCA (number of PC’s = 30) using the asinh-transformed counts which were scaled (min-max) across each marker and then corrected using “RunHarmony” function from the harmony R package (version 1.2.0), setting the “group.by.vars” argument to the sample patient id’s (n = 5). The harmony corrected, low-dimensional coordinates for each cell were visualised in two dimensions (Supplementary Figures 1-2) using the UMAP implementation from the scater R package (version 1.32.0), with “n_neighbors” = 35.

### Annotating Cell-Types

Cells with high expression (>90^th^ percentile) across multiple markers (>50% of marker) were excluded from cell phenotyping. The following markers were used to define each cell type: endothelial cell (SMA^+^ CD31^+^); NK cell (CD45^+^ NKP46^+^); T cell (CD45^+^ CD3^+^); macrophage (CD45^+^ IBA1^+^); microglia (CD45^+^ IBA1^+^ TMEM119^+^); neuron (NeuN_FOX3^+^ NKP46^+^ CD3^+^); astrocyte (GFAP^+^); oligodendrocyte (MOG^+^); astrocyte-like (SLC1A3_EAAT1^+^ HOPX^+^); mesenchymal-like (SOD2^+^ CHI3L1^+^ ANEXIN_A2^+^ ANXA1^+^); neural progenitor cell-like (DLL3^+^ BCAN^+^); oligodendrocyte progenitor-like (SCD5^+^ OLIG1^+^).

The asinh-transformed expression counts were scaled (z-score) across each marker and then each cell was ranked according to its marker expression from low (1) – high (20). Cells were assigned a phenotype label based on a logical gating criterion based on whether their marker expression rank exceeded a user-defined threshold, for e.g., cells were labelled as endothelial if their endothelial cell type marker (SMA and CD31) ranks were both greater than an empirically determined threshold.

The expression rank thresholds were determined separately for each cell-type in order to account for the relative differences in marker abundance: endothelial cell (>=15); NK cell (>=14); T cell (>=11); macrophage (>=15); microglia (>=12); neuron (>=16); astrocyte (>=17); oligodendrocyte (>=18); astrocyte-like (>=17); mesenchymal-like (>=17); neural progenitor cell-like (>=15); oligodendrocyte progenitor-like (>=15).

Some cell-type markers were challenging to label based solely on their marker expression rankings. This difficulty arose either because related cell types such as normal brain and cancer shared the same markers, but at different levels, or because the distinction between cell types depended on multiple markers with varying relative abundances. This was addressed by applying different thresholds to each marker and then by combining the intersecting cells to generate a single label.

### Annotating Cell-States

The following markers were used to define each cell state: hypoxia (HIF1A^+^); proliferative (Ki67^+^ SOX2^+^); quiescence (TGFBeta^+^ TNC^+^); epithelial to mesenchymal transition - EMT (SNAI1^+^).

The asinh-transformed expression counts were scaled (z-score) across each cell-state and states with multiple markers were summed after scaling to obtain a single score per cell-state. Each cell was then scored for low (< -1.2) and high (> 1.2) state expression based on the summarised z-scores values.

### Measurement of Intra-patient heterogeneity

Each ROI represents a mixture including immune, cancer, normal and vasculature cells. We measured intra-patient heterogeneity using Shannon entropy (*H*) and the annotated cell type labels. In order to account for the differences in cell frequencies between ROI’s, we sub-sampled 1000 cells in each group (*i*). This was repeated over ten rounds, and, in each round, we calculated the Shannon entropy of the labelled cell type frequencies (*P_c_*) as:

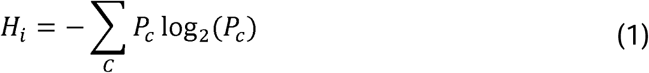

The Wilcoxon rank-sum test was used to compare the cell-type composition diversity across primary and recurrent surgeries using all of the data and also across each individual patient (n = 5).

### Spatial Cell Graphs

Undirected interaction graphs, in which each labelled cell-type corresponds to a node and where cells in spatial proximity are connected by an edge were generated for each ROI (n=30) using the Delaunay triangulation method, and the cell object centroids ^56^. The Delaunay method was favoured as it allowed us to better capture irregularly spaced and densely packed cells without setting an arbitrary distance thresholds or fixed neighbour counts. The “buildSpatialGraph” function from the imcRtools R package(version 1.10.0) was used build the graphs. In order to limit the number of spurious connections we also pruned the resulting graphs to only include edges that were within 50µm distance of each other by setting the “max_dist” argument.

### Testing Cell-Cell Interactions

We tested the cell-cell interactions defined in the spatial interaction graphs to see if cell types interacted more or less frequently than random, using a previously defined method ^57^. Briefly, each cell-cell interaction (edge) is summed and aggregated across a defined group (e.g., patient, surgery, cell-state etc.) and then divided by the number of cells of type A that have at least one neighbour of type B. The random component is derived by shuffling the cell-type labels across the same defined group, 1000 times, and for each iteration counting the interactions (as above), giving a null distribution which describes the interaction counts under spatial randomness. The observed interactions are then compared with the random distribution using two one-tailed permutation tests that define avoidance and interaction between cell types A and B for each group as:

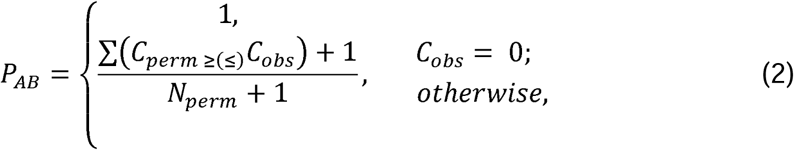

where *C_perm_* is the number of cell pairs (A, B) in each permutation, *C_obs_* is the actual number of cell pairs (A, B) given a defined distance, and *N_perm_* is the number of permutations. P values ≤ 0.01 were considered as significant interaction/avoidance between cell types. The method described above was implemented using the “testInteractions” function from the imcRtools R package.

### Cellular neighbourhoods

Cellular neighbourhoods (CN) were defined using a previously proposed method ^58,59^. Briefly, we aggregated the fraction of each cell type (using the annotated cell-type labels) within the direct (defined by their spatial graph) neighbourhood for every cell in each ROI (n=30) using the “aggregateNeighbors” function from the imcRtools R package. We then clustered the aggregated cell-type proportions using k-means clustering. The optimal k value was set as k=12, following a parameter sweep and visual inspection of a range of k values.

#### Spatial contexts

Spatial contexts (SCs) were identified using a previously proposed methodology ^60^. Briefly, for every cell in each ROI (n=30), we aggregated the fraction of cellular neighbourhood (identified above) present within a given cell’s direct neighbourhood (defined by their spatial graph) using the “aggregateNeighbors” function from the imcRtools R package. Then, for each cell we sorted (high-low) the CN fractions and assigned each cell an SC label as the set of CNs that cumulatively exceed a threshold of 90% (to capture the most dominant CNs) using the “detectSpatialContext” function from imcRtools R package. We also further filtered the SCs using the “filterSpatialContext” function from imcRtools R package to include only the most dominant SCs defined as: 1.) SCs that were present in at least three separate patients. 2.) SCs that contained a minimum number (>5% of the total cells) cells across each surgery type. We also assessed how the hypoxia and EMT cell-state protein marker abundance across each defined CN changed between primary and recurrent surgery samples (Supplementary Table 12).

## Authorship statement

Conception: LFS; Design: SA, AG, AAB, RAI, EW and LFS; Collection and assembly of data: SA, SP, GH, AC, AI, BH, AF and DM and LFS; Data analysis: SA; Interpretation of data: SA and LFS; Manuscript writing: SA; Final approval of manuscript: All authors; Accountable for all aspects of the study: All authors.

## Acknowledgements

This work was supported by grants from UK Research and Innovation [MR/T020504/1 to LFS], the Integrated Biological Imaging Network [IBIN4LS to LFS], Yorkshire’s Brain Tumour Charity and OSCARs Paediatric Brain Tumour Charity [Joint Infrastructure funding to LFS], the British Neuropathology Society [Small Grant Award to SA]. Research work in the Ihrie lab was supported by the Michael David Greene Brain Cancer Fund and the Ben & Catherine Ivy Foundation Emerging Leader Award. Tissue used in this study was accessed from the Sidney Driscol Neuroscience Foundation BTNW tissue bank. Tissue processing was possible through the Leeds Neuropathology Research Tissue Bank funded by Yorkshire’s Brain Tumour Charity and OSCARs Paediatric Brain Tumour Charity.

## Unpublished papers cited

No unpublished paper have been cited.

## Data availability

The data used in this study (including the raw high-dimensional TIFF images, spillover correction files; cell-object segmentation masks; patient and sample metadata; phenotype labelled single-cell data and the cell-object spatial information) are deposited and available online at Zenodo under the following DOI: https://doi.org/10.5281/zenodo.14679442

## Code availability

The analysis code that was used to process the IMC data and produce the results in this study can be found at: https://github.com/GliomaGenomics/GBM_IMC_Analysis

## References

1. Delgado-López, P. D. & Corrales-García, E. M. Survival in glioblastoma: a review on the impact of treatment modalities. Clin Transl Oncol 18, 1062–1071 (2016).

2. Stupp, R., Mason, W. P., Bent, M. J. van den, et al. Radiotherapy plus Concomitant and Adjuvant Temozolomide for Glioblastoma. New England Journal of Medicine 352, 987–996 (2005).

3. Barthel, F. P., Johnson, K. C., Varn, F. S., et al. Longitudinal molecular trajectories of diffuse glioma in adults. Nature 576, 112–120 (2019).

4. Körber, V., Yang, J., Barah, P., et al. Evolutionary Trajectories of IDHWT Glioblastomas Reveal a Common Path of Early Tumorigenesis Instigated Years ahead of Initial Diagnosis. Cancer Cell 35, 692–704.e12 (2019).

5. Wang, L., Babikir, H., Müller, S., et al. The Phenotypes of Proliferating Glioblastoma Cells Reside on a Single Axis of Variation. Cancer Discovery 9, 1708–1719 (2019).

6. Couturier, C. P., Ayyadhury, S., Le, P. U., et al. Single-cell RNA-seq reveals that glioblastoma recapitulates a normal neurodevelopmental hierarchy. Nat Commun 11, 3406 (2020).

7. Neftel, C., Laffy, J., Filbin, M. G., et al. An Integrative Model of Cellular States, Plasticity, and Genetics for Glioblastoma. Cell 178, 835–849.e21 (2019).

8. Hanahan, D. & Weinberg, R. A. Hallmarks of Cancer: The Next Generation. Cell 144, 646–674 (2011).

9. Neuroscience. (Sinauer Associates, Sunderland, Mass, 2001).

10. Quail, D. F. & Joyce, J. A. The Microenvironmental Landscape of Brain Tumors. Cancer Cell 31, 326–341 (2017).

11. Bressan, D., Battistoni, G. & Hannon, G. J. The dawn of spatial omics. *Science (New York*, N.Y.) 381, eabq4964 (2023).

12. Greenwald, A. C., Darnell, N. G., Hoefflin, R., et al. Integrative spatial analysis reveals a multi-layered organization of glioblastoma. Cell 187, 2485–2501.e26 (2024).

13. Manoharan, V. T., Abdelkareem, A., Gill, G., et al. Spatiotemporal modeling reveals high-resolution invasion states in glioblastoma. Genome Biology 25, 264 (2024).

14. Ravi, V. M., Will, P., Kueckelhaus, J., et al. Spatially resolved multi-omics deciphers bidirectional tumor-host interdependence in glioblastoma. Cancer Cell 40, 639–655.e13 (2022).

15. Giesen, C., Wang, H. A. O., Schapiro, D., et al. Highly multiplexed imaging of tumor tissues with subcellular resolution by mass cytometry. Nat Methods 11, 417–422 (2014).

16. Zhang, H., Zhou, Y., Cui, B., Liu, Z. & Shen, H. Novel insights into astrocyte-mediated signaling of proliferation, invasion and tumor immune microenvironment in glioblastoma. Biomedicine & Pharmacotherapy 126, 110086 (2020).

17. Gieryng, A., Pszczolkowska, D., Walentynowicz, K. A., Rajan, W. D. & Kaminska, B. Immune microenvironment of gliomas. Laboratory Investigation 97, 498–518 (2017).

18. Varn, F. S., Johnson, K. C., Martinek, J., et al. Glioma progression is shaped by genetic evolution and microenvironment interactions. Cell 185, 2184–2199.e16 (2022).

19. Tanner, G., Barrow, R., Ajaib, S., et al. IDHwt glioblastomas can be stratified by their transcriptional response to standard treatment, with implications for targeted therapy. Genome Biol 25, 45 (2024).

20. Chanoch-Myers, R., Wider, A., Suva, M. L. & Tirosh, I. Elucidating the diversity of malignant mesenchymal states in glioblastoma by integrative analysis. Genome Medicine 14, 106 (2022).

21. Gao, Z., Xu, J., Fan, Y., et al. PDIA3P1 promotes Temozolomide resistance in glioblastoma by inhibiting C/EBPβ degradation to facilitate proneural-to-mesenchymal transition. J Exp Clin Cancer Res 41, 223 (2022).

22. Yoon, S.-J., Shim, J.-K., Chang, J. H., et al. Tumor Mesenchymal Stem-Like Cell as a Prognostic Marker in Primary Glioblastoma. Stem Cells Int 2016, 6756983 (2016).

23. Carro, M. S., Lim, W. K., Alvarez, M. J., et al. The transcriptional network for mesenchymal transformation of brain tumours. Nature 463, 318–325 (2010).

24. Barriere, G., Fici, P., Gallerani, G., Fabbri, F. & Rigaud, M. Epithelial Mesenchymal Transition: a double-edged sword. Clinical and Translational Medicine 4, e14 (2015).

25. Spinelli, C., Adnani, L., Meehan, B., et al. Mesenchymal glioma stem cells trigger vasectasia—distinct neovascularization process stimulated by extracellular vesicles carrying EGFR. Nat Commun 15, 2865 (2024).

26. Cheng, J., Li, M., Motta, E., et al. Myeloid cells coordinately induce glioma cell-intrinsic and cell-extrinsic pathways for chemoresistance via GP130 signaling. Cell Reports Medicine 5, 101658 (2024).

27. Andreou, T., Williams, J., Brownlie, R. J., et al. Hematopoietic stem cell gene therapy targeting TGFβ enhances the efficacy of irradiation therapy in a preclinical glioblastoma model. J Immunother Cancer 9, e001143 (2021).

28. Albiach, A. M., Janusauskas, J., Kapustová, I., et al. Glioblastoma is spatially organized by neurodevelopmental programs and a glial-like wound healing response. 2023.09.01.555882 Preprint at 10.1101/2023.09.01.555882 (2023).

29. Brooks, L. J., Ragdale, H. S., Hill, C. S., Clements, M. & Parrinello, S. Injury programs shape glioblastoma. Trends in Neurosciences 45, 865–876 (2022).

30. Hide, T., Komohara, Y., Miyasato, Y., et al. Oligodendrocyte Progenitor Cells and Macrophages/Microglia Produce Glioma Stem Cell Niches at the Tumor Border. eBioMedicine 30, 94–104 (2018).

31. Kawashima, T., Yashiro, M., Kasashima, H., et al. Oligodendrocytes Up-regulate the Invasive Activity of Glioblastoma Cells via the Angiopoietin-2 Signaling Pathway. Anticancer Research 39, 577–584 (2019).

32. Huang, Y., Hoffman, C., Rajappa, P., et al. Oligodendrocyte Progenitor Cells Promote Neovascularization in Glioma by Disrupting the Blood–Brain Barrier. Cancer Research 74, 1011–1021 (2014).

33. Hortega, P. del R. & Penfield, W. Cerebral Cicatrix: The Reaction of Neuroglia and Microglia to Brain Wounds. (Johns Hopkins Hospital, 1927).

34. Escartin, C., Galea, E., Lakatos, A., et al. Reactive astrocyte nomenclature, definitions, and future directions. Nature neuroscience 24, 312 (2021).

35. Puchalski, R. B., Shah, N., Miller, J., et al. An anatomic transcriptional atlas of human glioblastoma. Science 360, 660–663 (2018).

36. Narni-Mancinelli, E., Vivier, E. & Kerdiles, Y. M. The ‘T-cell-ness’ of NK cells: unexpected similarities between NK cells and T cells. International Immunology 23, 427–431 (2011).

37. Maddison, K., Faulkner, S., Graves, M. C., et al. Vasculogenic Mimicry Occurs at Low Levels in Primary and Recurrent Glioblastoma. Cancers (Basel) 15, 3922 (2023).

38. Ahir, B. K., Engelhard, H. H. & Lakka, S. S. Tumor Development and Angiogenesis in Adult Brain Tumor: Glioblastoma. Mol Neurobiol 57, 2461–2478 (2020).

39. Wick, W., Brandes, AA., Gorlia, T., et al. LB-05PHASE III TRIAL EXPLORING THE COMBINATION OF BEVACIZUMAB AND LOMUSTINE IN PATIENTS WITH FIRST RECURRENCE OF A GLIOBLASTOMA: THE EORTC 26101 TRIAL. Neuro-Oncology 17, v1 (2015).

40. Hoogstrate, Y., Draaisma, K., Ghisai, S. A., et al. Transcriptome analysis reveals tumor microenvironment changes in glioblastoma. Cancer Cell 41, 678–692.e7 (2023).

41. Wang, L., Jung, J., Babikir, H., et al. A single-cell atlas of glioblastoma evolution under therapy reveals cell-intrinsic and cell-extrinsic therapeutic targets. Nat Cancer 3, 1534–1552 (2022).

42. Zhong, J., Paul, A., Kellie, S. J. & O’Neill, G. M. Mesenchymal Migration as a Therapeutic Target in Glioblastoma. J Oncol 2010, 430142 (2010).

43. van de Walle, T., Vaccaro, A., Ramachandran, M., et al. Tertiary Lymphoid Structures in the Central Nervous System: Implications for Glioblastoma. Front Immunol 12, 724739 (2021).

44. Stead, L. F. Treating glioblastoma often makes a MES. Nat Cancer 3, 1446–1448 (2022).

45. Pantazopoulou, V., Jeannot, P., Rosberg, R., Berg, T. J. & Pietras, A. Hypoxia-Induced Reactivity of Tumor-Associated Astrocytes Affects Glioma Cell Properties. Cells 10, 613 (2021).

46. Rosberg, R., Smolag, K. I., Sjölund, J., et al. Hypoxia-induced complement component 3 promotes aggressive tumor growth in the glioblastoma microenvironment. JCI Insight 9, (2024).

47. White, K., Connor, K., Meylan, M., et al. Identification, validation and biological characterisation of novel glioblastoma tumour microenvironment subtypes: implications for precision immunotherapy. Annals of Oncology 34, 300–314 (2023).

48. Ajaib, S., Lodha, D., Pollock, S., et al. GBMdeconvoluteR accurately infers proportions of neoplastic and immune cell populations from bulk glioblastoma transcriptomics data. Neuro-Oncology 25, 1236–1248 (2023).

49. R Core Team. R: A Language and Environment for Statistical Computing. (R Foundation for Statistical Computing, Vienna, Austria, 2022).

50. Wickham, H. Ggplot2: Elegant Graphics for Data Analysis. (Springer-Verlag New York, 2016).

51. Windhager, J., Zanotelli, V. R. T., Schulz, D., et al. An end-to-end workflow for multiplexed image processing and analysis. Nat Protoc 18, 3565–3613 (2023).

52. Pachitariu, M. & Stringer, C. Cellpose 2.0: how to train your own model. Nat Methods 19, 1634–1641 (2022).

53. Chevrier, S., Crowell, H. L., Zanotelli, V. R. T., et al. Compensation of Signal Spillover in Suspension and Imaging Mass Cytometry. Cell Syst 6, 612–620.e5 (2018).

54. Crowell, H. L., Chevrier, S., Jacobs, A., et al. An R-based reproducible and user-friendly preprocessing pipeline for CyTOF data. Preprint at 10.12688/f1000research.26073.2 (2022).

55. Korsunsky, I., Millard, N., Fan, J., et al. Fast, sensitive, and accurate integration of single cell data with Harmony. Nat Methods 16, 1289–1296 (2019).

56. Delaunay, B. Sur la sphère vide. A la mémoire de Georges Voronoï. Известия Российской академии наук. Серия математическая 793–800 (1934).

57. Schapiro, D., Jackson, H. W., Raghuraman, S., et al. histoCAT: analysis of cell phenotypes and interactions in multiplex image cytometry data. Nat Methods 14, 873–876 (2017).

58. Goltsev, Y., Samusik, N., Kennedy-Darling, J., et al. Deep Profiling of Mouse Splenic Architecture with CODEX Multiplexed Imaging. Cell 174, 968–981.e15 (2018).

59. Schürch, C. M., Bhate, S. S., Barlow, G. L., et al. Coordinated Cellular Neighborhoods Orchestrate Antitumoral Immunity at the Colorectal Cancer Invasive Front. Cell 182, 1341–1359.e19 (2020).

60. Bhate, S. S., Barlow, G. L., Schürch, C. M. & Nolan, G. P. Tissue schematics map the specialization of immune tissue motifs and their appropriation by tumors. cels 13, 109–130.e6 (2022).

