## Supplementary Information for "Spatial profiling of longitudinal glioblastoma reveals consistent changes in cellular architecture, post-treatment"

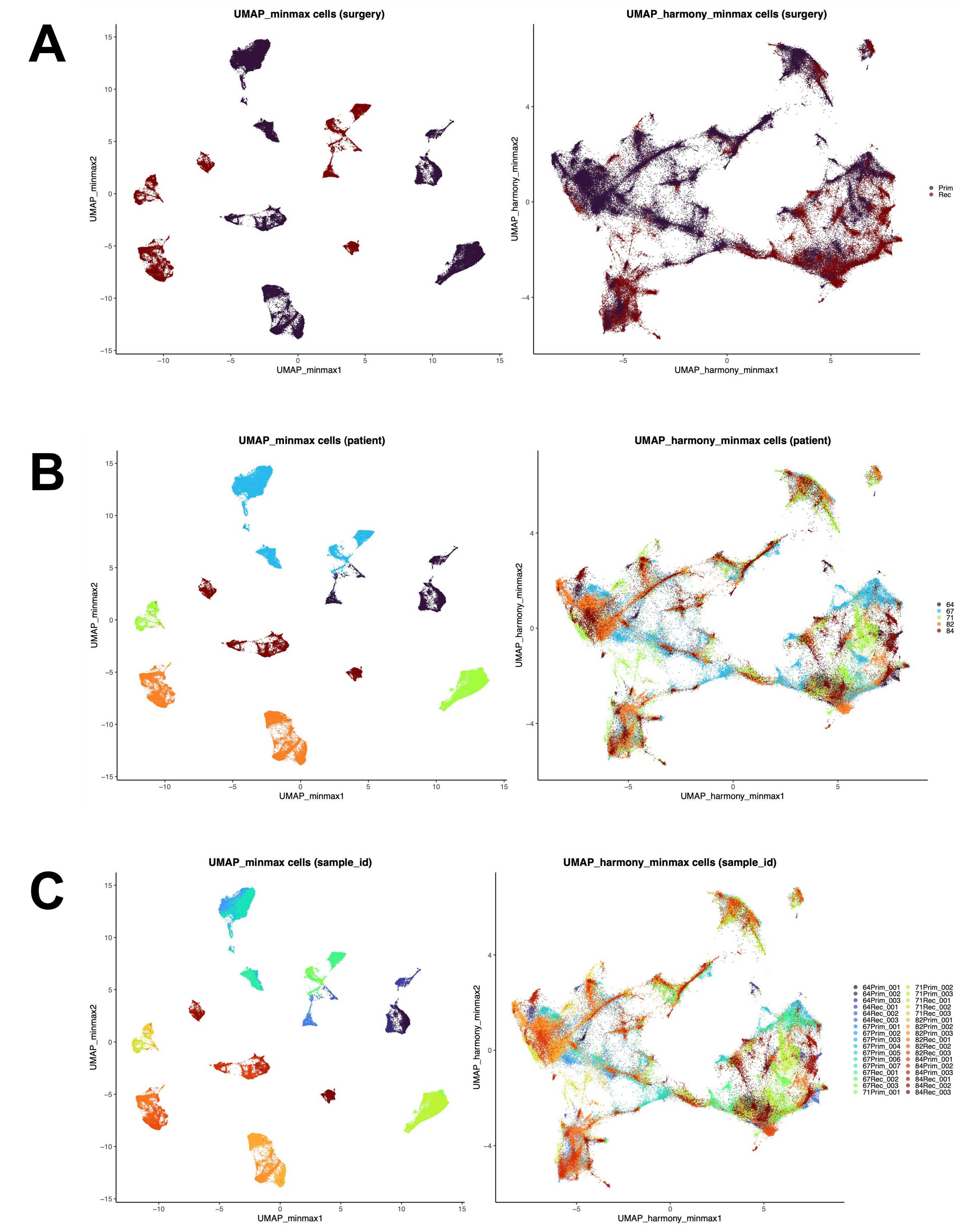


**Supplementary Figure 1. UMAP projections of all segmented single-cell objects before (left) and after (right) batch correction of min-max scaled protein marker abundances.** The projections are colored according to known sources of sample variation: **A)** surgery type; **B)** patient from which samples were obtained; **C)** patient and surgery-specific regions of interest (ROI).

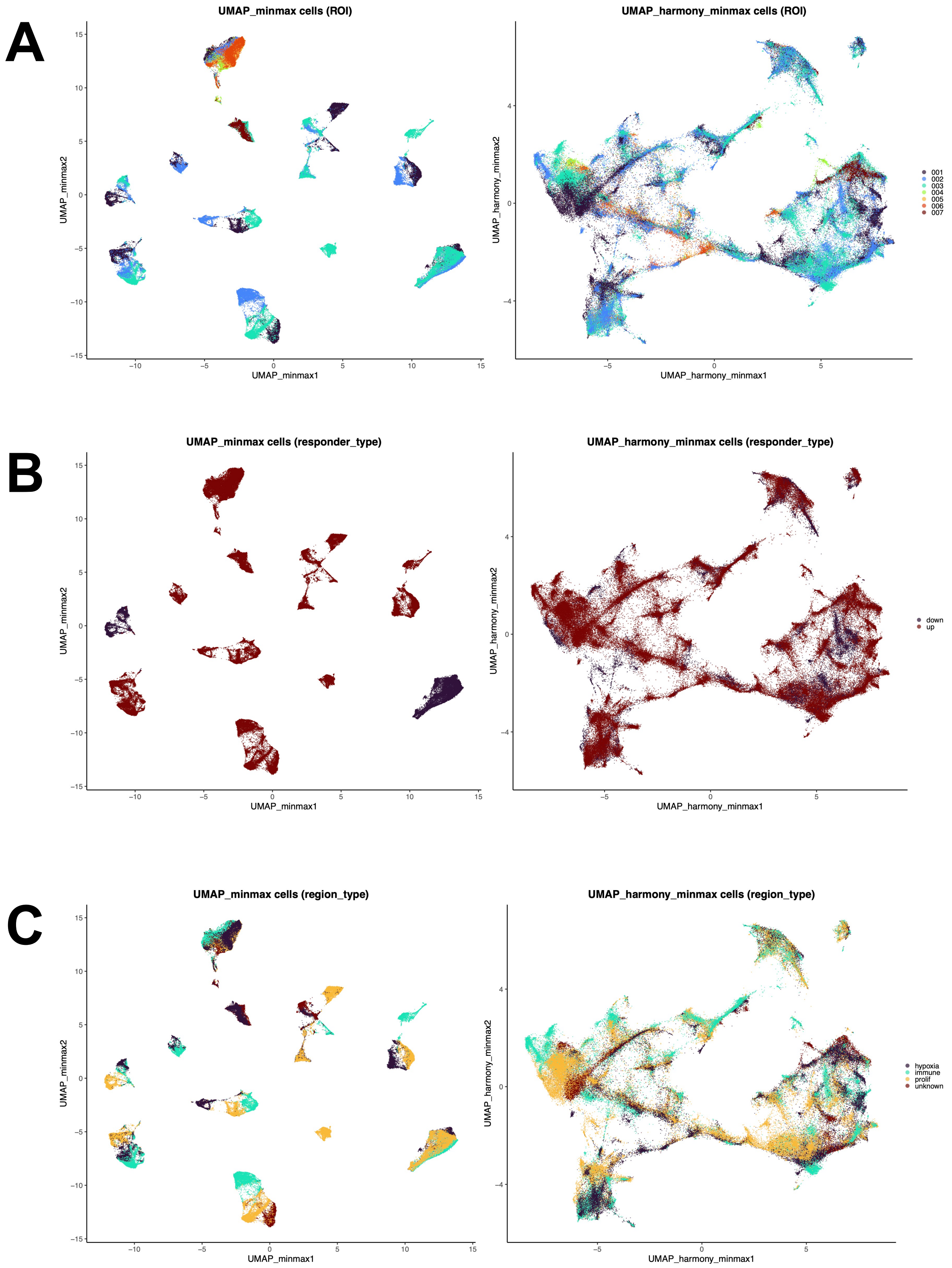


**Supplementary Figure 2. UMAP projections of all segmented single-cell objects before (left) and after (right) batch correction of min-max scaled protein marker abundances.** The projections are colored according to the following known sources of sample variation: **A)** regions of interest (ROI); **B)** responder types as defined in Tanner et al. *Genome Biology* 25, no. 1 (7 February 2024): 45; **C)** Immunohistochemical annotation of each ROI based on areas which have high levels of hypoxia, proliferation, and immune cells.

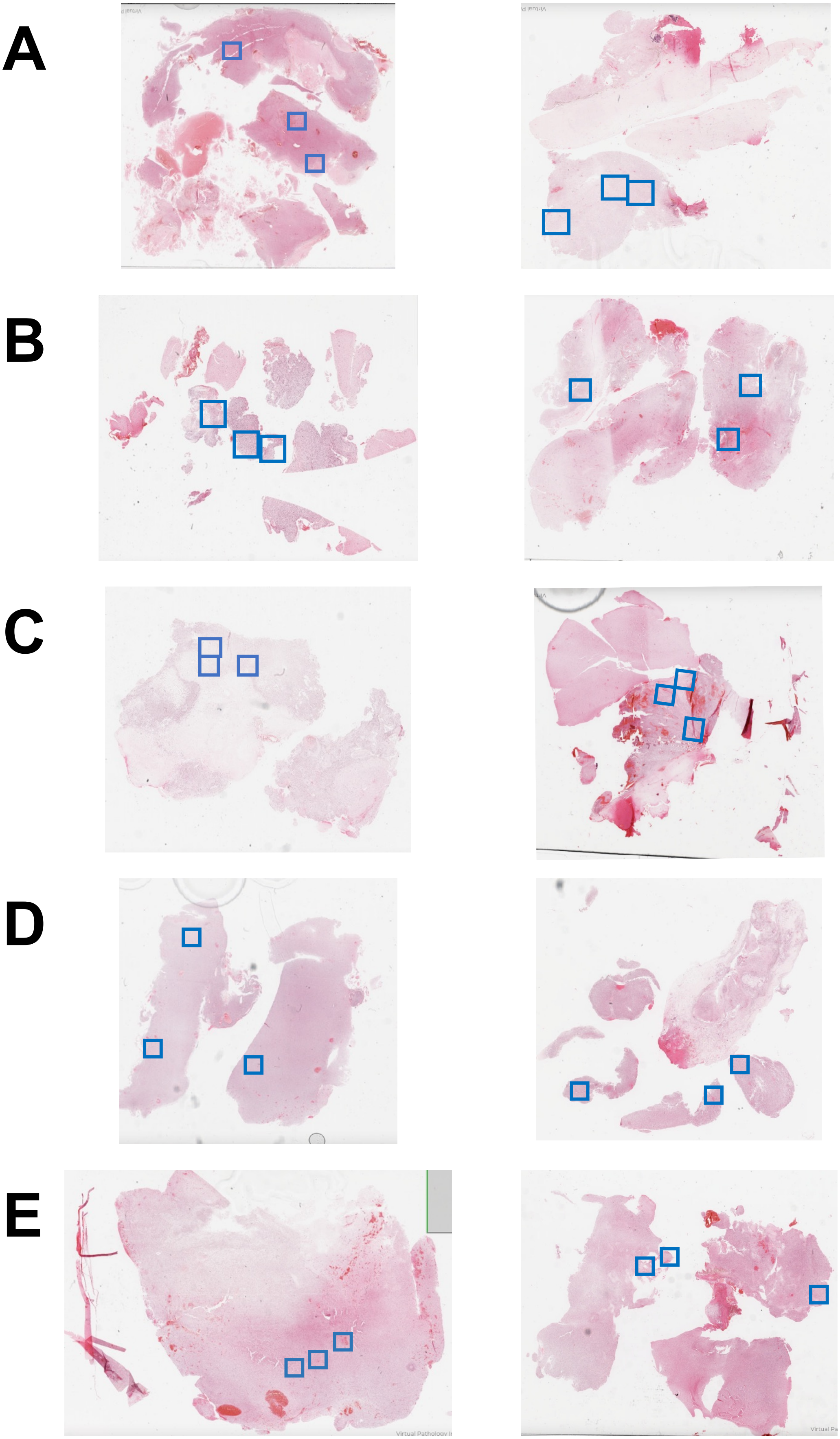


**Supplementary Figure 3. H&E stained FFPE sections of the five matched, primary (P) and recurrent (R) IDHwt glioblastoma tumour samples used in the study.** The blue demarcate the 1mm$^2$ regions of interest (ROI) that underwent imaging mass cytometry (IMC). The FFPE section correspond (from left to right) to the following patient/surgeries: **A)** 64P and 64R; **B)** 67P and 67R; **C)** 71P and 71R; **D)** 82P and 82R; **E)** 84P and 84R.

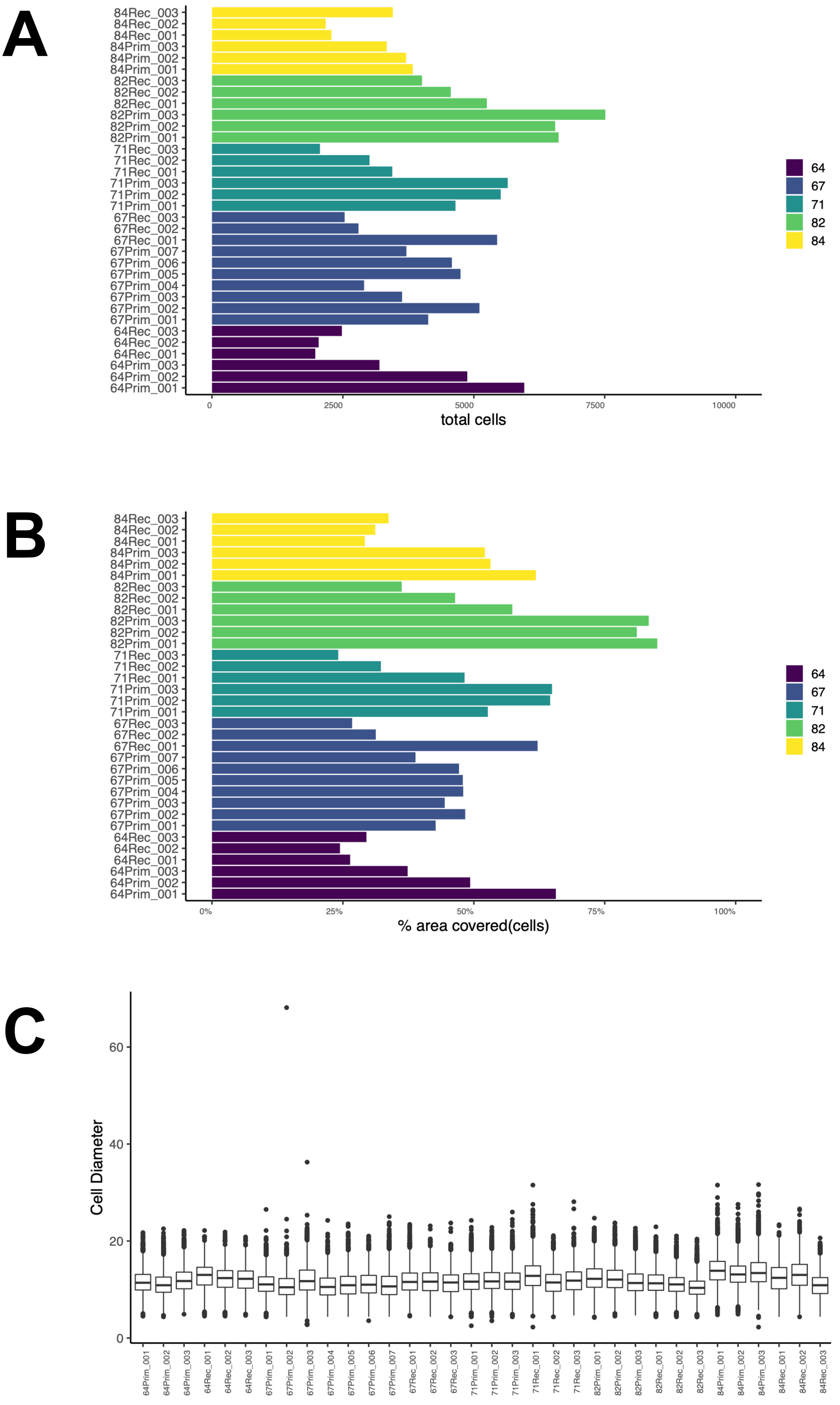


**Supplementary Figure 4. Segmented single-cell object metrics. A)** The total number of cells present in each region of interest (ROI) colored by patient; **B)** The percentage area covered by the segmented cell objects present across each ROI colored by patient; **C)** The distribution of cell diameters corresponding to the segmented cell objects present in each patient/surgery ROI.

**Supplementary Table 1. Clinical and molecular data of patients included in the study.**

| patient | sex | primary tumour location | diagnosis age | PFS (months) | OS (months) | status (1=deceased;0=alive) | responder type |
|---|---|---|---|---|---|---|---|
| 64 | Male | Frontal | 57 | 22.55 | 48.76 | 1.00 | up |
| 67 | Male | Temporal | 51 | 40.73 | 75.19 | 1.00 | up |
| 71 | Female | Temporal | 60 | 17.10 | 39.12 | 1.00 | down |
| 82 | Female | Parietal | 59 | 23.24 | 33.21 | 1.00 | up |
| 84 | Female | Parietal | 72 | 17.49 | 45.76 | 1.00 | up |

**Supplementary Table 2. Imaging mass cytometry (IMC) regions of interest (ROI) analysed in this study.**

| image id | patient | surgery | ROI | IHC annotation |
|---|---|---|---|---|
| 64Prim_001 | 64 | primary | 001 | prolif |
| 64Prim_002 | 64 | primary | 002 | hypoxia |
| 64Prim_003 | 64 | primary | 003 | immune |
| 64Rec_001 | 64 | recurrent | 001 | immune |
| 64Rec_002 | 64 | recurrent | 002 | hypoxia |
| 64Rec_003 | 64 | recurrent | 003 | prolif |
| 67Prim_001 | 67 | primary | 001 | immune |
| 67Prim_002 | 67 | primary | 002 | immune |
| 67Prim_003 | 67 | primary | 003 | unknown |
| 67Rec_001 | 67 | recurrent | 001 | prolif |
| 67Rec_002 | 67 | recurrent | 002 | hypoxia |
| 67Rec_003 | 67 | recurrent | 003 | unknown |
| 71Prim_001 | 71 | primary | 001 | hypoxia |
| 71Prim_002 | 71 | primary | 002 | immune |
| 71Prim_003 | 71 | primary | 003 | prolif |
| 71Rec_001 | 71 | recurrent | 001 | prolif |
| 71Rec_002 | 71 | recurrent | 002 | immune |
| 71Rec_003 | 71 | recurrent | 003 | hypoxia |
| 82Prim_001 | 82 | primary | 001 | unknown |
| 82Prim_002 | 82 | primary | 002 | immune |
| 82Prim_003 | 82 | primary | 003 | prolif |
| 82Rec_001 | 82 | recurrent | 001 | prolif |
| 82Rec_002 | 82 | recurrent | 002 | hypoxia |
| 82Rec_003 | 82 | recurrent | 003 | immune |
| 84Prim_001 | 84 | primary | 001 | prolif |
| 84Prim_002 | 84 | primary | 002 | hypoxia |
| 84Prim_003 | 84 | primary | 003 | immune |
| 84Rec_001 | 84 | recurrent | 001 | hypoxia |
| 84Rec_002 | 84 | recurrent | 002 | immune |
| 84Rec_003 | 84 | recurrent | 003 | prolif |

**Supplementary Table 3. Imaging mass cytometry (IMC) marker panel used in this study, including justification of marker/antibody selection.** Astrocyte-like (AC-like); bovine serum albumin (BSA); Cytometry by time of flight (CyTOF); Endoplasmic reticulum (ER); Epithelial to mesenchymal transition (EMT); Immunocytochemistry (ICC); Immunohistochemistry (IHC); immunofluorescence (IF); Knock-out (KO); Mesenchymal-like (MES-like); neural progenitor-like (NPC-like); oligodendrocyte progenitor-like (OPC-like); Phosphate buffered saline (PBS); PMID (PubMed Identifier).

| marker | cell category | cell type | cell state | location 1 | location 2 | marker justification (PMIDs) | manufacturer (antibody clone) | antibody justification | antibody concentration (µg/mL) | metal-isotope | control tissue |
|---|---|---|---|---|---|---|---|---|---|---|---|
| HOPX | cancer | AC-like | | cytoplasm | | 31327527; 31554641; 32641768 | abcam (ab230544) | | 100 | Yb171 | Brain, Tonsil |
| SLC1A3_EAAT1 | cancer | AC-like | | membrane | | 31327527; 31554641; 32641768 | abcam(ab240235) | BSA and azide free | 1000 | Gd158 | Brain |
| GFAP | normal | astrocyte | | cytoplasm | | 25726916 | abcam (ab218309) | PMID: 34174183 | 100 | Sm149 | Brain |
| CD56 | normal | neuron | | membrane | extracellular | 28791027 | biolegend (318345) | PMID: 28369679 | 100 | Dy162 | Brain |
| IBA1 | immune | macrophage | | cytoplasm | | 32848611 | abcam (ab220815) | PMID: 34174183 | 200 | Eu153 | Spleen, Tonsil |
| NeuN_FOX3 | normal | neuron | | nucleus | | 20452351 | biolegend (834502) | PMID: 34174183 | 400 | Sm147 | Brain |
| ANXA_A1 | cancer | MES-like | | cytoplasm | membrane | 31327527; 31554641; 32641768 | abcam (ab222398) | BSA and azide free | | Yb172 | Tonsil |
| ANXA_A2 | cancer | MES-like | | cytoplasm | membrane | 31327527; 31554641; 32641768 | rndsystems (mab3928) | In PBS with Trehalose | 8-25 | Er166 | Prostate, Tonsil |
| CHI3L1 | cancer | MES-like | | cytoplasm | extracellular | 31327527; 31554641; 32641768 | abcam (ab255864) | BSA and azide free | 250 | Sm154 | Spleen, Brain |
| SOD2 | cancer | MES-like | | mitochondria | | 31327527; 31554641; 32641768 | abcam (ab227846) | BSA and azide free, used in IHC | 100 | Nd146 | Prostate |
| P2Y12R | immune | microglia | | membrane | | 32848611 | abcam (ab274386) | BSA and azide free | 1000 | Lu175 | Brain |
| TMEM119 | immune | microglia | | cytoplasm | membrane | 32848611 | sigmaaldrich (HPA051870) | PMID: 31740814 | 500-1000 | Gd155 | Brain, Tonsil |
| NKP46 | immune | NK cell | | membrane | | 31784984 | rndsystems (mab1850) | PMID: 36689332 | 5-25 | Nd144 | Spleen |
| BCAN | cancer | NPC-like | | extracellular | nucleus | 31327527; 31554641; 32641768 | thermofisher (MA5-27639) | BSA free, used in ICC | 50 | Gd160 | Brain |
| DLL3 | cancer | NPC-like | | membrane | | 31327527; 31554641; 32641768 | abcam (ab255694) | BSA and azide free | 100 | Nd148 | Brain |
| MOG | normal | oligodendrocyte | | membrane | | 2649509 | rndsystems (mab1850) | | 5-25 | Gd157 | Brain |
| OLIG1 | cancer | OPC-like | | nucleus | | 31327527; 31554641; 32641768 | rndsystems (mab2417) | carrier free, used in IHC | 8-25 | Yb174 | Skin |
| SCD5 | cancer | OPC-like | | ER | | 31327527; 31554641; 32641768 | thermofisher (PA5-59963) | used in IHC | 50 | Tm169 | Brain |
| CD3 | immune | T cell | | membrane | | 29768164 | fluidigm (3170019D) | PMID: 36689332 | 75-200 | Er170 | Spleen, Tonsil |
| CD8 | immune | T cell | | membrane | | 29768164 | biolegend (344727) | PMID: 28369679 | 200 | Ho165 | Spleen, Tonsil |
| DNA1 | DNA intercalator | | | nucleus | | | fluidigm (201192B) | Preconugated to 191Ir | | Ir191 | |
| DNA2 | DNA intercalator | | | nucleus | | | fluidigm (201192B) | Preconugated to 193Ir | | Ir193 | |
| CD45 | immune | | | membrane | | 12414720 | fluidigm (91H029152) | Preconugated to 152Sm | 300 | Sm152 | Spleen, Tonsil |
| CD31 | vasculature | | | membrane | | 27055047 | fluidigm (3151025D) | Preconugated to 151Eu | | Eu151 | Skin, Tonsil, Prostate |
| SMA | vasculature | | | cytoplasm | | 19929197 | rndsystems (mab1420) | used in Cytof | 8-25 | Dy164 | Prostate, Skin, Tonsil |
| EZH2 | | | transcript respressive | nucleus | | 23720055 | abcam (ab231165) | BSA and azide free | 250 | Nd145 | Tonsil |
| HIF1A | | | hypoxia | cytoplasm | nucleus | 11606368 | thermofisher (700505) | PMID: 32868913 | 400 | Dy161 | Bone marrow |
| JARID2_C_Terminus | | | active | nucleus | | 30573669 | developed in house | | | Nd143 | Brain |
| JARID2_N_Terminus | | | repressed | nucleus | | 30573669 | abcam (ab251123) | BSA free version, validated (by KO) | | Yb173 | Brain |
| Ki67 | | | proliferating | nucleus | | 29322240 | fluidigm (3168001B) | Preconugated to 168Er | | Er168 | Skin, Tonsil |
| SNAI1 | | | EMT | nucleus | cytoplasm | 33806868 | rndsystems (af3639) | BSA and azide free, used in IHC | 5-15 | Tb159 | Ubiquitous |
| SOX2 | | | proliferating | nucleus | | 30952620 | fluidigm (3150019B) | Preconugated to 150Nd | | Nd150 | Brain, Tonsil |
| TGFBeta | | | quiescent | extracellular | | 30952620 | fluidigm (3163010B) | Pre-conjugated to 163Dy | | Dy163 | Spleen, Bone marrow, Prostate |
| TNC | | | quiescent | extracellular | | 30952620 | rndsystems (mab2138) | used in IF | 8-25 | Gd156 | Uterus |

**Supplementary Table 4. ANOVA results comparing intra-tumour (across patient regions of interest, ROI) and inter-tumor (within patient and surgery sample) heterogeneity of cell categories.** The p value significance levels are denoted using the following symbols: **** ($p < 0.0001$); *** ($p < 0.001$); ** ($p < 0.01$); * ($p < 0.05$); n.s (not significant).

| cell category | source of variation | effect size (F statistic) | p value | p significance |
|---|---|---|---|---|
| Immune | patient & surgery | 5.82 | 7.62E-04 | *** |
| Cancer | patient & surgery | 9.47 | 3.33E-05 | **** |
| Normal | patient & surgery | 10.11 | 2.11E-05 | **** |
| Vasculature | patient & surgery | 4.56 | 3.01E-03 | ** |
| Immune | within tumor (ROI) | 0.32 | 7.29E-01 | n.s |
| Cancer | within tumor (ROI) | 2.63 | 9.97E-02 | n.s |
| Normal | within tumor (ROI) | 1.23 | 3.15E-01 | n.s |
| Vasculature | within tumor (ROI) | 1.75 | 2.02E-01 | n.s |

**Supplementary Table 5. Comparison of cell category prevalence between primary and recurrent samples.** Statistical significance was assessed using an unpaired Wilcoxon test, with adjusted p-values calculated using the false discovery rate (FDR) method.

| cell category | comparison groups | n (per comparison group) | p value | adjusted p value |
|---|---|---|---|---|
| Immune | Prim vs Rec | 15 | 9.75E-02 | 9.75E-02 |
| Cancer | Prim vs Rec | 15 | 6.13E-02 | 8.17E-02 |
| Normal | Prim vs Rec | 15 | 1.13E-04 | 4.52E-04 |
| Vasculature | Prim vs Rec | 15 | 4.94E-03 | 9.88E-03 |

**Supplementary Table 6. Changes in cell type prevalence between primary and recurrent samples.** Statistical significance was assessed using the unpaired Wilcoxon test, with adjusted p-values calculated using the false discovery rate (FDR) method.

| cell type | comparison groups | n (per comparison group) | p value | adjusted p value |
|---|---|---|---|---|
| T cell | Prim vs Rec | 15 | 5.12E-01 | 6.83E-01 |
| NK cell | Prim vs Rec | 15 | 2.02E-01 | 3.72E-01 |
| Macrophage | Prim vs Rec | 15 | 2.17E-01 | 3.72E-01 |
| Microglia | Prim vs Rec | 15 | 2.50E-01 | 3.75E-01 |
| AC | Prim vs Rec | 15 | 6.53E-02 | 1.57E-01 |
| MES | Prim vs Rec | 15 | 9.17E-01 | 9.17E-01 |
| NPC | Prim vs Rec | 15 | 8.33E-01 | 9.09E-01 |
| OPC | Prim vs Rec | 15 | 8.14E-01 | 9.09E-01 |
| Neuron | Prim vs Rec | 15 | 4.17E-03 | 1.98E-02 |
| Astrocyte | Prim vs Rec | 15 | 3.02E-03 | 1.98E-02 |
| Oligodendrocyte | Prim vs Rec | 15 | 2.25E-02 | 6.75E-02 |
| Endothelial | Prim vs Rec | 15 | 4.94E-03 | 1.98E-02 |

**Supplementary Table 7. Comparison of Shannon's entropy (H) between primary and recurrent samples, quantifying intra-tumour cellular heterogeneity.** Statistical significance was assessed using the unpaired, Wilcoxon test, with adjusted p-values calculated using the false discovery rate (FDR) method.

| patient(s) | comparision group | n (per comparison group) | p value | adjusted p value |
|---|---|---|---|---|
| All | Prim vs Rec | 150 | 3.93E-03 | 3.93E-03 |
| 64 | Prim vs Rec | 30 | 1.43E-04 | 3.58E-04 |
| 67 | Prim vs Rec | 30 | 2.63E-02 | 2.63E-02 |
| 71 | Prim vs Rec | 30 | 1.69E-17 | 8.45E-17 |
| 82 | Prim vs Rec | 30 | 1.52E-03 | 1.90E-03 |
| 84 | Prim vs Rec | 30 | 4.26E-04 | 7.10E-04 |

**Supplementary Table 8. Changes in cellular states (hypoxia and Epithelial to mesenchymal transition - EMT) across neoplastic GBM cancer cell types in primary and recurrent samples.** Statistical significance was assessed using the unpaired, Wilcoxon test, with adjusted p-values calculated using the false discovery rate (FDR) method. The p value significance levels are denoted using the following symbols: **** (p < 0.0001); *** (p < 0.001); ** (p < 0.01); * (p < 0.05); n.s (not significant).

| cancer cell type | cellular state | comparison groups | n cells (primary) | n cells (recurrent) | p value | adjusted p value | p significance |
|---|---|---|---|---|---|---|---|
| AC | hypoxia | Prim vs Rec | 5522 | 686 | 4.98E-115 | 9.96E-115 | **** |
| MES | hypoxia | Prim vs Rec | 9141 | 3119 | 9.62E-125 | 3.85E-124 | **** |
| NPC | hypoxia | Prim vs Rec | 1664 | 613 | 5.49E-70 | 7.32E-70 | **** |
| OPC | hypoxia | Prim vs Rec | 1743 | 616 | 7.41E-01 | 7.41E-01 | n.s |
| AC | EMT | Prim vs Rec | 5522 | 686 | 5.29E-161 | 1.06E-160 | **** |
| MES | EMT | Prim vs Rec | 9141 | 3119 | 1.03E-07 | 1.37E-07 | **** |
| NPC | EMT | Prim vs Rec | 1664 | 613 | 2.32E-183 | 9.28E-183 | **** |
| OPC | EMT | Prim vs Rec | 1743 | 616 | 4.40E-03 | 4.40E-03 | ** |

**Supplementary Table 9. Patient-specific, observed, cell-cell interactions compared to a null model of spatial randomness.** The results are separated by patient and surgery type (primary or recurrent). Statistical significance and direction is determined using a permutation test, with p-values indicating interactions more or less likely than random: 1 (significant positive interactions); -1 (significant avoidance interactions); 0 (neutral and/or non- statistically significant interactions).

| row | patient | surgery | from cell type | to cell type | observed count | permutations (greater than) | permutations (less than) | p value | interaction significance/direction |
|---|---|---|---|---|---|---|---|---|---|
| 1 | 64 | primary | AC | AC | 2.96 | 9.99E-04 | 1.00E+00 | 9.99E-04 | 1 |
| 2 | 64 | primary | AC | Astrocyte | 1.05 | 2.90E-02 | 9.72E-01 | 2.90E-02 | 0 |
| 3 | 64 | primary | AC | Endothelial | 1.42 | 9.99E-04 | 1.00E+00 | 9.99E-04 | 1 |
| 4 | 64 | primary | AC | MES | 1.62 | 1.00E+00 | 9.99E-04 | 9.99E-04 | -1 |
| 5 | 64 | primary | AC | Macrophage | 1.49 | 9.99E-04 | 1.00E+00 | 9.99E-04 | 1 |
| 6 | 64 | primary | AC | Microglia | 1.24 | 9.16E-01 | 8.49E-02 | 8.49E-02 | 0 |
| 7 | 64 | primary | AC | NK cell | 1.16 | 9.99E-04 | 1.00E+00 | 9.99E-04 | 1 |
| 8 | 64 | primary | AC | NPC | 1.00 | 1.00E+00 | 9.91E-01 | 9.91E-01 | 0 |
| 9 | 64 | primary | AC | Neuron | 1.06 | 1.10E-02 | 9.92E-01 | 1.10E-02 | 0 |
| 10 | 64 | primary | AC | OPC | 1.00 | 9.57E-01 | 9.99E-01 | 9.57E-01 | 0 |
| 11 | 64 | primary | AC | Oligodendrocyte | 1.33 | 1.31E-01 | 8.70E-01 | 1.31E-01 | 0 |
| 12 | 64 | primary | AC | T cell | 1.31 | 2.00E-03 | 9.99E-01 | 2.00E-03 | 1 |
| 13 | 64 | primary | Astrocyte | AC | 2.24 | 9.99E-04 | 1.00E+00 | 9.99E-04 | 1 |
| 14 | 64 | primary | Astrocyte | Astrocyte | 1.08 | 1.80E-02 | 9.83E-01 | 1.80E-02 | 0 |
| 15 | 64 | primary | Astrocyte | Endothelial | 1.53 | 1.70E-02 | 9.84E-01 | 1.70E-02 | 0 |
| 16 | 64 | primary | Astrocyte | MES | 2.60 | 9.99E-04 | 1.00E+00 | 9.99E-04 | 1 |
| 17 | 64 | primary | Astrocyte | Macrophage | 1.58 | 2.90E-02 | 9.73E-01 | 2.90E-02 | 0 |
| 18 | 64 | primary | Astrocyte | Microglia | 1.14 | 9.21E-01 | 8.19E-02 | 8.19E-02 | 0 |
| 19 | 64 | primary | Astrocyte | NK cell | 1.00 | 9.98E-01 | 7.54E-01 | 7.54E-01 | 0 |
| 20 | 64 | primary | Astrocyte | NPC | 1.00 | 1.53E-01 | 1.00E+00 | 1.53E-01 | 0 |
| 21 | 64 | primary | Astrocyte | Neuron | 1.00 | 4.21E-01 | 1.00E+00 | 4.21E-01 | 0 |
| 22 | 64 | primary | Astrocyte | OPC | 0.00 | 1.00E+00 | 9.34E-01 | 9.34E-01 | 0 |
| 23 | 64 | primary | Astrocyte | Oligodendrocyte | 1.19 | 9.00E-01 | 1.06E-01 | 1.06E-01 | 0 |
| 24 | 64 | primary | Astrocyte | T cell | 1.10 | 9.74E-01 | 2.80E-02 | 2.80E-02 | 0 |
| 25 | 64 | primary | Endothelial | AC | 1.88 | 9.99E-04 | 1.00E+00 | 9.99E-04 | 1 |
| 26 | 64 | primary | Endothelial | Astrocyte | 1.09 | 9.99E-03 | 9.91E-01 | 9.99E-03 | 1 |
| 27 | 64 | primary | Endothelial | Endothelial | 2.06 | 9.99E-04 | 1.00E+00 | 9.99E-04 | 1 |
| 28 | 64 | primary | Endothelial | MES | 1.90 | 9.99E-04 | 1.00E+00 | 9.99E-04 | 1 |
| 29 | 64 | primary | Endothelial | Macrophage | 1.44 | 2.20E-02 | 9.79E-01 | 2.20E-02 | 0 |
| 30 | 64 | primary | Endothelial | Microglia | 1.35 | 9.99E-04 | 1.00E+00 | 9.99E-04 | 1 |
| 31 | 64 | primary | Endothelial | NK cell | 1.24 | 9.99E-04 | 1.00E+00 | 9.99E-04 | 1 |
| 32 | 64 | primary | Endothelial | NPC | 1.00 | 9.54E-01 | 9.98E-01 | 9.54E-01 | 0 |
| 33 | 64 | primary | Endothelial | Neuron | 1.00 | 1.00E+00 | 9.71E-01 | 9.71E-01 | 0 |
| 34 | 64 | primary | Endothelial | OPC | 1.00 | 7.01E-01 | 1.00E+00 | 7.01E-01 | 0 |
| 35 | 64 | primary | Endothelial | Oligodendrocyte | 1.39 | 9.99E-04 | 1.00E+00 | 9.99E-04 | 1 |
| 36 | 64 | primary | Endothelial | T cell | 1.33 | 9.99E-04 | 1.00E+00 | 9.99E-04 | 1 |
| 37 | 64 | primary | MES | AC | 1.77 | 4.80E-02 | 9.53E-01 | 4.80E-02 | 0 |
| 38 | 64 | primary | MES | Astrocyte | 1.07 | 5.00E-03 | 9.96E-01 | 5.00E-03 | 1 |
| 39 | 64 | primary | MES | Endothelial | 1.38 | 9.99E-04 | 1.00E+00 | 9.99E-04 | 1 |
| 40 | 64 | primary | MES | MES | 3.59 | 9.99E-04 | 1.00E+00 | 9.99E-04 | 1 |
| 41 | 64 | primary | MES | Macrophage | 1.30 | 1.00E+00 | 9.99E-04 | 9.99E-04 | -1 |
| 42 | 64 | primary | MES | Microglia | 1.28 | 7.79E-02 | 9.23E-01 | 7.79E-02 | 0 |
| 43 | 64 | primary | MES | NK cell | 1.25 | 9.99E-04 | 1.00E+00 | 9.99E-04 | 1 |
| 44 | 64 | primary | MES | NPC | 1.00 | 9.98E-01 | 9.99E-01 | 9.98E-01 | 0 |
| 45 | 64 | primary | MES | Neuron | 1.00 | 1.00E+00 | 9.46E-01 | 9.46E-01 | 0 |
| 46 | 64 | primary | MES | OPC | 0.00 | 1.00E+00 | 5.00E-02 | 5.00E-02 | 0 |
| 47 | 64 | primary | MES | Oligodendrocyte | 1.59 | 9.99E-04 | 1.00E+00 | 9.99E-04 | 1 |
| 48 | 64 | primary | MES | T cell | 1.33 | 9.99E-04 | 1.00E+00 | 9.99E-04 | 1 |
| 49 | 64 | primary | Macrophage | AC | 2.60 | 9.99E-04 | 1.00E+00 | 9.99E-04 | 1 |
| 50 | 64 | primary | Macrophage | Astrocyte | 1.12 | 9.99E-04 | 1.00E+00 | 9.99E-04 | 1 |
| 51 | 64 | primary | Macrophage | Endothelial | 1.50 | 9.99E-04 | 1.00E+00 | 9.99E-04 | 1 |
| 52 | 64 | primary | Macrophage | MES | 2.42 | 9.99E-04 | 1.00E+00 | 9.99E-04 | 1 |
| 53 | 64 | primary | Macrophage | Macrophage | 1.59 | 9.99E-04 | 1.00E+00 | 9.99E-04 | 1 |
| 54 | 64 | primary | Macrophage | Microglia | 1.48 | 9.99E-04 | 1.00E+00 | 9.99E-04 | 1 |
| 55 | 64 | primary | Macrophage | NK cell | 1.19 | 9.99E-04 | 1.00E+00 | 9.99E-04 | 1 |
| 56 | 64 | primary | Macrophage | NPC | 1.00 | 9.86E-01 | 9.97E-01 | 9.86E-01 | 0 |
| 57 | 64 | primary | Macrophage | Neuron | 1.00 | 1.00E+00 | 9.66E-01 | 9.66E-01 | 0 |
| 58 | 64 | primary | Macrophage | OPC | 1.00 | 8.16E-01 | 9.99E-01 | 8.16E-01 | 0 |
| 59 | 64 | primary | Macrophage | Oligodendrocyte | 1.46 | 9.99E-04 | 1.00E+00 | 9.99E-04 | 1 |
| 60 | 64 | primary | Macrophage | T cell | 1.32 | 2.00E-03 | 9.99E-01 | 2.00E-03 | 1 |
| 61 | 64 | primary | Microglia | AC | 2.40 | 9.99E-04 | 1.00E+00 | 9.99E-04 | 1 |
| 62 | 64 | primary | Microglia | Astrocyte | 1.09 | 7.99E-03 | 9.93E-01 | 7.99E-03 | 1 |
| 63 | 64 | primary | Microglia | Endothelial | 1.52 | 9.99E-04 | 1.00E+00 | 9.99E-04 | 1 |
| 64 | 64 | primary | Microglia | MES | 1.94 | 9.99E-04 | 1.00E+00 | 9.99E-04 | 1 |
| 65 | 64 | primary | Microglia | Macrophage | 1.52 | 9.99E-04 | 1.00E+00 | 9.99E-04 | 1 |
| 66 | 64 | primary | Microglia | Microglia | 1.72 | 9.99E-04 | 1.00E+00 | 9.99E-04 | 1 |
| 67 | 64 | primary | Microglia | NK cell | 1.44 | 9.99E-04 | 1.00E+00 | 9.99E-04 | 1 |
| 68 | 64 | primary | Microglia | NPC | 1.00 | 9.43E-01 | 9.98E-01 | 9.43E-01 | 0 |
| 69 | 64 | primary | Microglia | Neuron | 0.00 | 1.00E+00 | 9.99E-04 | 9.99E-04 | -1 |
| 70 | 64 | primary | Microglia | OPC | 0.00 | 1.00E+00 | 3.25E-01 | 3.25E-01 | 0 |
| 71 | 64 | primary | Microglia | Oligodendrocyte | 1.50 | 9.99E-04 | 1.00E+00 | 9.99E-04 | 1 |
| 72 | 64 | primary | Microglia | T cell | 1.49 | 9.99E-04 | 1.00E+00 | 9.99E-04 | 1 |
| 73 | 64 | primary | NK cell | AC | 2.17 | 9.99E-04 | 1.00E+00 | 9.99E-04 | 1 |
| 74 | 64 | primary | NK cell | Astrocyte | 1.00 | 9.98E-01 | 8.84E-01 | 8.84E-01 | 0 |
| 75 | 64 | primary | NK cell | Endothelial | 1.49 | 2.00E-03 | 9.99E-01 | 2.00E-03 | 1 |
| 76 | 64 | primary | NK cell | MES | 1.38 | 1.00E+00 | 9.99E-04 | 9.99E-04 | -1 |
| 77 | 64 | primary | NK cell | Macrophage | 1.46 | 1.05E-01 | 8.96E-01 | 1.05E-01 | 0 |
| 78 | 64 | primary | NK cell | Microglia | 1.79 | 9.99E-04 | 1.00E+00 | 9.99E-04 | 1 |
| 79 | 64 | primary | NK cell | NK cell | 1.78 | 9.99E-04 | 1.00E+00 | 9.99E-04 | 1 |
| 80 | 64 | primary | NK cell | NPC | 0.00 | 1.00E+00 | 5.77E-01 | 5.77E-01 | 0 |
| 81 | 64 | primary | NK cell | Neuron | 1.00 | 7.89E-01 | 9.98E-01 | 7.89E-01 | 0 |
| 82 | 64 | primary | NK cell | OPC | 1.00 | 1.97E-01 | 1.00E+00 | 1.97E-01 | 0 |
| 83 | 64 | primary | NK cell | Oligodendrocyte | 1.06 | 1.00E+00 | 9.99E-04 | 9.99E-04 | -1 |
| 84 | 64 | primary | NK cell | T cell | 1.70 | 9.99E-04 | 1.00E+00 | 9.99E-04 | 1 |
| 85 | 64 | primary | NPC | AC | 2.33 | 1.03E-01 | 9.15E-01 | 1.03E-01 | 0 |
| 86 | 64 | primary | NPC | Astrocyte | 1.00 | 1.53E-01 | 9.94E-01 | 1.53E-01 | 0 |
| 87 | 64 | primary | NPC | Endothelial | 2.00 | 1.13E-01 | 9.81E-01 | 1.13E-01 | 0 |
| 88 | 64 | primary | NPC | MES | 1.50 | 7.03E-01 | 4.05E-01 | 4.05E-01 | 0 |
| 89 | 64 | primary | NPC | Macrophage | 1.00 | 9.86E-01 | 3.71E-01 | 3.71E-01 | 0 |
| 90 | 64 | primary | NPC | Microglia | 1.50 | 2.40E-01 | 8.69E-01 | 2.40E-01 | 0 |
| 91 | 64 | primary | NPC | NK cell | 0.00 | 1.00E+00 | 5.77E-01 | 5.77E-01 | 0 |
| 92 | 64 | primary | NPC | NPC | 0.00 | 1.00E+00 | 9.99E-01 | 9.99E-01 | 0 |
| 93 | 64 | primary | NPC | Neuron | 0.00 | 1.00E+00 | 9.67E-01 | 9.67E-01 | 0 |
| 94 | 64 | primary | NPC | OPC | 0.00 | 1.00E+00 | 9.96E-01 | 9.96E-01 | 0 |
| 95 | 64 | primary | NPC | Oligodendrocyte | 0.00 | 1.00E+00 | 4.30E-02 | 4.30E-02 | 0 |
| 96 | 64 | primary | NPC | T cell | 1.33 | 3.73E-01 | 7.19E-01 | 3.73E-01 | 0 |
| 97 | 64 | primary | Neuron | AC | 2.83 | 2.00E-03 | 9.99E-01 | 2.00E-03 | 1 |
| 98 | 64 | primary | Neuron | Astrocyte | 1.00 | 4.21E-01 | 9.90E-01 | 4.21E-01 | 0 |
| 99 | 64 | primary | Neuron | Endothelial | 1.50 | 1.63E-01 | 8.96E-01 | 1.63E-01 | 0 |
| 100 | 64 | primary | Neuron | MES | 4.50 | 9.99E-04 | 1.00E+00 | 9.99E-04 | 1 |

| row | patient | surgery | from cell type | to cell tye | observed count | permutations (greater than) | permutations (less than) | p value | interaction significance/direction |
|---|---|---|---|---|---|---|---|---|---|
| 101 | 64 | primary | Neuron | Macrophage | 1.33 | 6.02E-01 | 4.83E-01 | 4.83E-01 | 0 |
| 102 | 64 | primary | Neuron | Microglia | 0.00 | 1.00E+00 | 9.99E-04 | 9.99E-04 | -1 |
| 103 | 64 | primary | Neuron | NK cell | 1.00 | 7.89E-01 | 9.36E-01 | 7.89E-01 | 0 |
| 104 | 64 | primary | Neuron | NPC | 0.00 | 1.00E+00 | 9.67E-01 | 9.67E-01 | 0 |
| 105 | 64 | primary | Neuron | Neuron | 0.00 | 1.00E+00 | 9.55E-01 | 9.55E-01 | 0 |
| 106 | 64 | primary | Neuron | OPC | 0.00 | 1.00E+00 | 9.85E-01 | 9.85E-01 | 0 |
| 107 | 64 | primary | Neuron | Oligodendrocyte | 1.50 | 2.30E-01 | 8.49E-01 | 2.30E-01 | 0 |
| 108 | 64 | primary | Neuron | T cell | 1.00 | 9.99E-01 | 2.35E-01 | 2.35E-01 | 0 |
| 109 | 64 | primary | OPC | AC | 4.00 | 1.80E-02 | 9.97E-01 | 1.80E-02 | 0 |
| 110 | 64 | primary | OPC | Astrocyte | 0.00 | 1.00E+00 | 9.34E-01 | 9.34E-01 | 0 |
| 111 | 64 | primary | OPC | Endothelial | 1.00 | 7.01E-01 | 7.93E-01 | 7.01E-01 | 0 |
| 112 | 64 | primary | OPC | MES | 0.00 | 1.00E+00 | 5.00E-02 | 5.00E-02 | 0 |
| 113 | 64 | primary | OPC | Macrophage | 2.00 | 1.99E-01 | 9.60E-01 | 1.99E-01 | 0 |
| 114 | 64 | primary | OPC | Microglia | 0.00 | 1.00E+00 | 3.25E-01 | 3.25E-01 | 0 |
| 115 | 64 | primary | OPC | NK cell | 1.00 | 1.97E-01 | 9.89E-01 | 1.97E-01 | 0 |
| 116 | 64 | primary | OPC | NPC | 0.00 | 1.00E+00 | 9.96E-01 | 9.96E-01 | 0 |
| 117 | 64 | primary | OPC | Neuron | 0.00 | 1.00E+00 | 9.85E-01 | 9.85E-01 | 0 |
| 118 | 64 | primary | OPC | OPC | 0.00 | 1.00E+00 | 1.00E+00 | 1.00E+00 | 0 |
| 119 | 64 | primary | OPC | Oligodendrocyte | 1.00 | 7.40E-01 | 7.57E-01 | 7.40E-01 | 0 |
| 120 | 64 | primary | OPC | T cell | 0.00 | 1.00E+00 | 3.13E-01 | 3.13E-01 | 0 |
| 121 | 64 | primary | Oligodendrocyte | AC | 2.31 | 9.99E-04 | 1.00E+00 | 9.99E-04 | 1 |
| 122 | 64 | primary | Oligodendrocyte | Astrocyte | 1.00 | 1.00E+00 | 5.58E-01 | 5.58E-01 | 0 |
| 123 | 64 | primary | Oligodendrocyte | Endothelial | 1.48 | 9.99E-04 | 1.00E+00 | 9.99E-04 | 1 |
| 124 | 64 | primary | Oligodendrocyte | MES | 1.84 | 9.99E-04 | 1.00E+00 | 9.99E-04 | 1 |
| 125 | 64 | primary | Oligodendrocyte | Macrophage | 1.38 | 5.84E-01 | 4.18E-01 | 4.18E-01 | 0 |
| 126 | 64 | primary | Oligodendrocyte | Microglia | 1.32 | 2.00E-03 | 9.99E-01 | 2.00E-03 | 1 |
| 127 | 64 | primary | Oligodendrocyte | NK cell | 1.09 | 2.50E-02 | 9.79E-01 | 2.50E-02 | 0 |
| 128 | 64 | primary | Oligodendrocyte | NPC | 0.00 | 1.00E+00 | 4.30E-02 | 4.30E-02 | 0 |
| 129 | 64 | primary | Oligodendrocyte | Neuron | 1.00 | 1.00E+00 | 9.70E-01 | 9.70E-01 | 0 |
| 130 | 64 | primary | Oligodendrocyte | OPC | 1.00 | 7.40E-01 | 1.00E+00 | 7.40E-01 | 0 |
| 131 | 64 | primary | Oligodendrocyte | Oligodendrocyte | 1.86 | 9.99E-04 | 1.00E+00 | 9.99E-04 | 1 |
| 132 | 64 | primary | Oligodendrocyte | T cell | 1.56 | 9.99E-04 | 1.00E+00 | 9.99E-04 | 1 |
| 133 | 64 | primary | T cell | AC | 2.13 | 9.99E-04 | 1.00E+00 | 9.99E-04 | 1 |
| 134 | 64 | primary | T cell | Astrocyte | 1.00 | 1.00E+00 | 5.98E-01 | 5.98E-01 | 0 |
| 135 | 64 | primary | T cell | Endothelial | 1.48 | 9.99E-04 | 1.00E+00 | 9.99E-04 | 1 |
| 136 | 64 | primary | T cell | MES | 2.04 | 9.99E-04 | 1.00E+00 | 9.99E-04 | 1 |
| 137 | 64 | primary | T cell | Macrophage | 1.37 | 6.53E-01 | 3.48E-01 | 3.48E-01 | 0 |
| 138 | 64 | primary | T cell | Microglia | 1.36 | 9.99E-04 | 1.00E+00 | 9.99E-04 | 1 |
| 139 | 64 | primary | T cell | NK cell | 1.48 | 9.99E-04 | 1.00E+00 | 9.99E-04 | 1 |
| 140 | 64 | primary | T cell | NPC | 1.00 | 9.30E-01 | 9.97E-01 | 9.30E-01 | 0 |
| 141 | 64 | primary | T cell | Neuron | 1.00 | 9.99E-01 | 9.71E-01 | 9.71E-01 | 0 |
| 142 | 64 | primary | T cell | OPC | 0.00 | 1.00E+00 | 3.13E-01 | 3.13E-01 | 0 |
| 143 | 64 | primary | T cell | Oligodendrocyte | 1.78 | 9.99E-04 | 1.00E+00 | 9.99E-04 | 1 |
| 144 | 64 | primary | T cell | T cell | 1.81 | 9.99E-04 | 1.00E+00 | 9.99E-04 | 1 |
| 145 | 64 | recurrent | AC | AC | | | | | |
| 146 | 64 | recurrent | AC | Astrocyte | | | | | |
| 147 | 64 | recurrent | AC | Endothelial | | | | | |
| 148 | 64 | recurrent | AC | MES | | | | | |
| 149 | 64 | recurrent | AC | Macrophage | | | | | |
| 150 | 64 | recurrent | AC | Microglia | | | | | |
| 151 | 64 | recurrent | AC | NK cell | | | | | |
| 152 | 64 | recurrent | AC | NPC | | | | | |
| 153 | 64 | recurrent | AC | Neuron | | | | | |
| 154 | 64 | recurrent | AC | OPC | | | | | |
| 155 | 64 | recurrent | AC | Oligodendrocyte | | | | | |
| 156 | 64 | recurrent | AC | T cell | | | | | |
| 157 | 64 | recurrent | Astrocyte | AC | | | | | |
| 158 | 64 | recurrent | Astrocyte | Astrocyte | 2.99 | 9.99E-04 | 1.00E+00 | 9.99E-04 | 1 |
| 159 | 64 | recurrent | Astrocyte | Endothelial | 1.15 | 4.00E-03 | 9.97E-01 | 4.00E-03 | 1 |
| 160 | 64 | recurrent | Astrocyte | MES | 1.24 | 3.00E-03 | 9.98E-01 | 3.00E-03 | 1 |
| 161 | 64 | recurrent | Astrocyte | Macrophage | 1.13 | 9.65E-01 | 3.60E-02 | 3.60E-02 | 0 |
| 162 | 64 | recurrent | Astrocyte | Microglia | 1.16 | 1.50E-02 | 9.87E-01 | 1.50E-02 | 0 |
| 163 | 64 | recurrent | Astrocyte | NK cell | 1.17 | 1.29E-01 | 8.72E-01 | 1.29E-01 | 0 |
| 164 | 64 | recurrent | Astrocyte | NPC | 1.34 | 9.99E-04 | 1.00E+00 | 9.99E-04 | 1 |
| 165 | 64 | recurrent | Astrocyte | Neuron | 1.20 | 9.99E-04 | 1.00E+00 | 9.99E-04 | 1 |
| 166 | 64 | recurrent | Astrocyte | OPC | 0.00 | 1.00E+00 | 1.42E-01 | 1.42E-01 | 0 |
| 167 | 64 | recurrent | Astrocyte | Oligodendrocyte | 1.91 | 9.99E-04 | 1.00E+00 | 9.99E-04 | 1 |
| 168 | 64 | recurrent | Astrocyte | T cell | 1.11 | 9.50E-01 | 5.09E-02 | 5.09E-02 | 0 |
| 169 | 64 | recurrent | Endothelial | AC | | | | | |
| 170 | 64 | recurrent | Endothelial | Astrocyte | 2.46 | 9.99E-04 | 1.00E+00 | 9.99E-04 | 1 |
| 171 | 64 | recurrent | Endothelial | Endothelial | 1.88 | 9.99E-04 | 1.00E+00 | 9.99E-04 | 1 |
| 172 | 64 | recurrent | Endothelial | MES | 1.17 | 5.55E-01 | 4.65E-01 | 4.65E-01 | 0 |
| 173 | 64 | recurrent | Endothelial | Macrophage | 1.28 | 3.20E-02 | 9.69E-01 | 3.20E-02 | 0 |
| 174 | 64 | recurrent | Endothelial | Microglia | 1.22 | 5.09E-02 | 9.50E-01 | 5.09E-02 | 0 |
| 175 | 64 | recurrent | Endothelial | NK cell | 1.83 | 9.99E-04 | 1.00E+00 | 9.99E-04 | 1 |
| 176 | 64 | recurrent | Endothelial | NPC | 1.30 | 2.40E-02 | 9.78E-01 | 2.40E-02 | 0 |
| 177 | 64 | recurrent | Endothelial | Neuron | 1.16 | 1.28E-01 | 8.73E-01 | 1.28E-01 | 0 |
| 178 | 64 | recurrent | Endothelial | OPC | 0.00 | 1.00E+00 | 8.17E-01 | 8.17E-01 | 0 |
| 179 | 64 | recurrent | Endothelial | Oligodendrocyte | 1.35 | 1.00E+00 | 9.99E-04 | 9.99E-04 | -1 |
| 180 | 64 | recurrent | Endothelial | T cell | 1.05 | 9.50E-01 | 5.29E-02 | 5.29E-02 | 0 |
| 181 | 64 | recurrent | MES | AC | | | | | |
| 182 | 64 | recurrent | MES | Astrocyte | 1.85 | 9.93E-01 | 7.99E-03 | 7.99E-03 | -1 |
| 183 | 64 | recurrent | MES | Endothelial | 1.17 | 3.50E-02 | 9.73E-01 | 3.50E-02 | 0 |
| 184 | 64 | recurrent | MES | MES | 1.70 | 9.99E-04 | 1.00E+00 | 9.99E-04 | 1 |
| 185 | 64 | recurrent | MES | Macrophage | 1.39 | 9.99E-04 | 1.00E+00 | 9.99E-04 | 1 |
| 186 | 64 | recurrent | MES | Microglia | 1.18 | 3.50E-02 | 9.67E-01 | 3.50E-02 | 0 |
| 187 | 64 | recurrent | MES | NK cell | 1.13 | 5.86E-01 | 4.19E-01 | 4.19E-01 | 0 |
| 188 | 64 | recurrent | MES | NPC | 1.43 | 9.99E-04 | 1.00E+00 | 9.99E-04 | 1 |
| 189 | 64 | recurrent | MES | Neuron | 1.00 | 1.00E+00 | 5.00E-03 | 5.00E-03 | -1 |
| 190 | 64 | recurrent | MES | OPC | 1.00 | 3.16E-01 | 1.00E+00 | 3.16E-01 | 0 |
| 191 | 64 | recurrent | MES | Oligodendrocyte | 3.02 | 9.99E-04 | 1.00E+00 | 9.99E-04 | 1 |
| 192 | 64 | recurrent | MES | T cell | 1.17 | 2.43E-01 | 7.61E-01 | 2.43E-01 | 0 |
| 193 | 64 | recurrent | Macrophage | AC | | | | | |
| 194 | 64 | recurrent | Macrophage | Astrocyte | 1.82 | 9.95E-01 | 5.99E-03 | 5.99E-03 | -1 |
| 195 | 64 | recurrent | Macrophage | Endothelial | 1.40 | 9.99E-04 | 1.00E+00 | 9.99E-04 | 1 |
| 196 | 64 | recurrent | Macrophage | MES | 1.44 | 9.99E-04 | 1.00E+00 | 9.99E-04 | 1 |
| 197 | 64 | recurrent | Macrophage | Macrophage | 1.50 | 9.99E-04 | 1.00E+00 | 9.99E-04 | 1 |
| 198 | 64 | recurrent | Macrophage | Microglia | 1.25 | 2.00E-03 | 9.99E-01 | 2.00E-03 | 1 |
| 199 | 64 | recurrent | Macrophage | NK cell | 1.47 | 9.99E-04 | 1.00E+00 | 9.99E-04 | 1 |
| 200 | 64 | recurrent | Macrophage | NPC | 1.35 | 2.00E-03 | 9.99E-01 | 2.00E-03 | 1 |
| 201 | 64 | recurrent | Macrophage | Neuron | 1.23 | 3.00E-03 | 9.98E-01 | 3.00E-03 | 1 |
| 202 | 64 | recurrent | Macrophage | OPC | 1.00 | 3.08E-01 | 1.00E+00 | 3.08E-01 | 0 |
| 203 | 64 | recurrent | Macrophage | Oligodendrocyte | 2.86 | 9.99E-04 | 1.00E+00 | 9.99E-04 | 1 |
| 204 | 64 | recurrent | Macrophage | T cell | 1.20 | 8.29E-02 | 9.18E-01 | 8.29E-02 | 0 |
| 205 | 64 | recurrent | Microglia | AC | | | | | |
| 206 | 64 | recurrent | Microglia | Astrocyte | 2.53 | 9.99E-04 | 1.00E+00 | 9.99E-04 | 1 |
| 207 | 64 | recurrent | Microglia | Endothelial | 1.36 | 9.99E-04 | 1.00E+00 | 9.99E-04 | 1 |
| 208 | 64 | recurrent | Microglia | MES | 1.40 | 9.99E-04 | 1.00E+00 | 9.99E-04 | 1 |
| 209 | 64 | recurrent | Microglia | Macrophage | 1.53 | 9.99E-04 | 1.00E+00 | 9.99E-04 | 1 |
| 210 | 64 | recurrent | Microglia | Microglia | 1.24 | 5.09E-02 | 9.51E-01 | 5.09E-02 | 0 |

| row | patient | surgery | from cell type | to cell tye | observed count | permutations (greater than) | permutations (less than) | p value | interaction significance/direction |
|---|---|---|---|---|---|---|---|---|---|
| 211 | 64 | recurrent | Microglia | NK cell | 1.33 | 2.00E-03 | 9.99E-01 | 2.00E-03 | 1 |
| 212 | 64 | recurrent | Microglia | NPC | 1.35 | 2.00E-03 | 9.99E-01 | 2.00E-03 | 1 |
| 213 | 64 | recurrent | Microglia | Neuron | 1.00 | 1.00E+00 | 4.90E-02 | 4.90E-02 | 0 |
| 214 | 64 | recurrent | Microglia | OPC | 0.00 | 1.00E+00 | 7.61E-01 | 7.61E-01 | 0 |
| 215 | 64 | recurrent | Microglia | Oligodendrocyte | 2.37 | 9.99E-04 | 1.00E+00 | 9.99E-04 | 1 |
| 216 | 64 | recurrent | Microglia | T cell | 1.15 | 4.23E-01 | 5.87E-01 | 4.23E-01 | 0 |
| 217 | 64 | recurrent | NK cell | AC | | | | | |
| 218 | 64 | recurrent | NK cell | Astrocyte | 2.56 | 9.99E-04 | 1.00E+00 | 9.99E-04 | 1 |
| 219 | 64 | recurrent | NK cell | Endothelial | 1.94 | 9.99E-04 | 1.00E+00 | 9.99E-04 | 1 |
| 220 | 64 | recurrent | NK cell | MES | 1.31 | 6.99E-03 | 9.95E-01 | 6.99E-03 | 1 |
| 221 | 64 | recurrent | NK cell | Macrophage | 1.49 | 9.99E-04 | 1.00E+00 | 9.99E-04 | 1 |
| 222 | 64 | recurrent | NK cell | Microglia | 1.26 | 2.00E-03 | 9.99E-01 | 2.00E-03 | 1 |
| 223 | 64 | recurrent | NK cell | NK cell | 1.81 | 9.99E-04 | 1.00E+00 | 9.99E-04 | 1 |
| 224 | 64 | recurrent | NK cell | NPC | 1.35 | 9.99E-04 | 1.00E+00 | 9.99E-04 | 1 |
| 225 | 64 | recurrent | NK cell | Neuron | 1.11 | 3.01E-01 | 7.20E-01 | 3.01E-01 | 0 |
| 226 | 64 | recurrent | NK cell | OPC | 0.00 | 1.00E+00 | 7.13E-01 | 7.13E-01 | 0 |
| 227 | 64 | recurrent | NK cell | Oligodendrocyte | 2.15 | 9.99E-04 | 1.00E+00 | 9.99E-04 | 1 |
| 228 | 64 | recurrent | NK cell | T cell | 1.20 | 8.49E-02 | 9.16E-01 | 8.49E-02 | 0 |
| 229 | 64 | recurrent | NPC | AC | | | | | |
| 230 | 64 | recurrent | NPC | Astrocyte | 2.69 | 9.99E-04 | 1.00E+00 | 9.99E-04 | 1 |
| 231 | 64 | recurrent | NPC | Endothelial | 1.17 | 4.40E-02 | 9.60E-01 | 4.40E-02 | 0 |
| 232 | 64 | recurrent | NPC | MES | 1.18 | 4.55E-01 | 5.54E-01 | 4.55E-01 | 0 |
| 233 | 64 | recurrent | NPC | Macrophage | 1.10 | 9.47E-01 | 5.59E-02 | 5.59E-02 | 0 |
| 234 | 64 | recurrent | NPC | Microglia | 1.11 | 4.76E-01 | 5.30E-01 | 4.76E-01 | 0 |
| 235 | 64 | recurrent | NPC | NK cell | 1.15 | 4.43E-01 | 5.64E-01 | 4.43E-01 | 0 |
| 236 | 64 | recurrent | NPC | NPC | 1.42 | 9.99E-04 | 1.00E+00 | 9.99E-04 | 1 |
| 237 | 64 | recurrent | NPC | Neuron | 1.23 | 4.00E-03 | 9.97E-01 | 4.00E-03 | 1 |
| 238 | 64 | recurrent | NPC | OPC | 0.00 | 1.00E+00 | 7.06E-01 | 7.06E-01 | 0 |
| 239 | 64 | recurrent | NPC | Oligodendrocyte | 1.11 | 1.00E+00 | 9.99E-04 | 9.99E-04 | -1 |
| 240 | 64 | recurrent | NPC | T cell | 1.04 | 9.98E-01 | 3.00E-03 | 3.00E-03 | -1 |
| 241 | 64 | recurrent | Neuron | AC | | | | | |
| 242 | 64 | recurrent | Neuron | Astrocyte | 2.70 | 9.99E-04 | 1.00E+00 | 9.99E-04 | 1 |
| 243 | 64 | recurrent | Neuron | Endothelial | 1.10 | 3.59E-01 | 6.62E-01 | 3.59E-01 | 0 |
| 244 | 64 | recurrent | Neuron | MES | 1.30 | 3.20E-02 | 9.71E-01 | 3.20E-02 | 0 |
| 245 | 64 | recurrent | Neuron | Macrophage | 1.07 | 9.59E-01 | 4.50E-02 | 4.50E-02 | 0 |
| 246 | 64 | recurrent | Neuron | Microglia | 1.05 | 8.64E-01 | 1.42E-01 | 1.42E-01 | 0 |
| 247 | 64 | recurrent | Neuron | NK cell | 1.11 | 7.00E-01 | 3.09E-01 | 3.09E-01 | 0 |
| 248 | 64 | recurrent | Neuron | NPC | 1.52 | 9.99E-04 | 1.00E+00 | 9.99E-04 | 1 |
| 249 | 64 | recurrent | Neuron | Neuron | 1.28 | 1.40E-02 | 9.87E-01 | 1.40E-02 | 0 |
| 250 | 64 | recurrent | Neuron | OPC | 0.00 | 1.00E+00 | 8.22E-01 | 8.22E-01 | 0 |
| 251 | 64 | recurrent | Neuron | Oligodendrocyte | 1.21 | 1.00E+00 | 9.99E-04 | 9.99E-04 | -1 |
| 252 | 64 | recurrent | Neuron | T cell | 1.10 | 7.54E-01 | 2.52E-01 | 2.52E-01 | 0 |
| 253 | 64 | recurrent | OPC | AC | | | | | |
| 254 | 64 | recurrent | OPC | Astrocyte | 0.00 | 1.00E+00 | 1.42E-01 | 1.42E-01 | 0 |
| 255 | 64 | recurrent | OPC | Endothelial | 0.00 | 1.00E+00 | 8.17E-01 | 8.17E-01 | 0 |
| 256 | 64 | recurrent | OPC | MES | 1.00 | 3.16E-01 | 9.49E-01 | 3.16E-01 | 0 |
| 257 | 64 | recurrent | OPC | Macrophage | 1.00 | 3.08E-01 | 9.60E-01 | 3.08E-01 | 0 |
| 258 | 64 | recurrent | OPC | Microglia | 0.00 | 1.00E+00 | 7.61E-01 | 7.61E-01 | 0 |
| 259 | 64 | recurrent | OPC | NK cell | 0.00 | 1.00E+00 | 7.13E-01 | 7.13E-01 | 0 |
| 260 | 64 | recurrent | OPC | NPC | 0.00 | 1.00E+00 | 7.06E-01 | 7.06E-01 | 0 |
| 261 | 64 | recurrent | OPC | Neuron | 0.00 | 1.00E+00 | 8.22E-01 | 8.22E-01 | 0 |
| 262 | 64 | recurrent | OPC | OPC | 0.00 | 1.00E+00 | 1.00E+00 | 1.00E+00 | 0 |
| 263 | 64 | recurrent | OPC | Oligodendrocyte | 5.00 | 5.00E-03 | 9.99E-01 | 5.00E-03 | 1 |
| 264 | 64 | recurrent | OPC | T cell | 0.00 | 1.00E+00 | 7.29E-01 | 7.29E-01 | 0 |
| 265 | 64 | recurrent | Oligodendrocyte | AC | | | | | |
| 266 | 64 | recurrent | Oligodendrocyte | Astrocyte | 1.36 | 1.00E+00 | 9.99E-04 | 9.99E-04 | -1 |
| 267 | 64 | recurrent | Oligodendrocyte | Endothelial | 1.41 | 9.99E-04 | 1.00E+00 | 9.99E-04 | 1 |
| 268 | 64 | recurrent | Oligodendrocyte | MES | 1.46 | 9.99E-04 | 1.00E+00 | 9.99E-04 | 1 |
| 269 | 64 | recurrent | Oligodendrocyte | Macrophage | 1.40 | 9.99E-04 | 1.00E+00 | 9.99E-04 | 1 |
| 270 | 64 | recurrent | Oligodendrocyte | Microglia | 1.17 | 8.99E-03 | 9.92E-01 | 8.99E-03 | 1 |
| 271 | 64 | recurrent | Oligodendrocyte | NK cell | 1.20 | 1.10E-02 | 9.90E-01 | 1.10E-02 | 0 |
| 272 | 64 | recurrent | Oligodendrocyte | NPC | 1.39 | 9.99E-04 | 1.00E+00 | 9.99E-04 | 1 |
| 273 | 64 | recurrent | Oligodendrocyte | Neuron | 1.12 | 1.34E-01 | 8.67E-01 | 1.34E-01 | 0 |
| 274 | 64 | recurrent | Oligodendrocyte | OPC | 1.00 | 7.66E-01 | 1.00E+00 | 7.66E-01 | 0 |
| 275 | 64 | recurrent | Oligodendrocyte | Oligodendrocyte | 3.31 | 9.99E-04 | 1.00E+00 | 9.99E-04 | 1 |
| 276 | 64 | recurrent | Oligodendrocyte | T cell | 1.21 | 4.00E-03 | 9.97E-01 | 4.00E-03 | 1 |
| 277 | 64 | recurrent | T cell | AC | | | | | |
| 278 | 64 | recurrent | T cell | Astrocyte | 2.46 | 9.99E-04 | 1.00E+00 | 9.99E-04 | 1 |
| 279 | 64 | recurrent | T cell | Endothelial | 1.44 | 9.99E-04 | 1.00E+00 | 9.99E-04 | 1 |
| 280 | 64 | recurrent | T cell | MES | 1.41 | 9.99E-04 | 1.00E+00 | 9.99E-04 | 1 |
| 281 | 64 | recurrent | T cell | Macrophage | 1.32 | 2.00E-03 | 9.99E-01 | 2.00E-03 | 1 |
| 282 | 64 | recurrent | T cell | Microglia | 1.17 | 9.99E-02 | 9.07E-01 | 9.99E-02 | 0 |
| 283 | 64 | recurrent | T cell | NK cell | 1.26 | 7.99E-03 | 9.93E-01 | 7.99E-03 | 1 |
| 284 | 64 | recurrent | T cell | NPC | 1.45 | 9.99E-04 | 1.00E+00 | 9.99E-04 | 1 |
| 285 | 64 | recurrent | T cell | Neuron | 1.26 | 3.00E-03 | 9.98E-01 | 3.00E-03 | 1 |
| 286 | 64 | recurrent | T cell | OPC | 0.00 | 1.00E+00 | 7.29E-01 | 7.29E-01 | 0 |
| 287 | 64 | recurrent | T cell | Oligodendrocyte | 2.52 | 9.99E-04 | 1.00E+00 | 9.99E-04 | 1 |
| 288 | 64 | recurrent | T cell | T cell | 1.47 | 9.99E-04 | 1.00E+00 | 9.99E-04 | 1 |
| 289 | 67 | primary | AC | AC | 1.00 | 4.58E-01 | 9.87E-01 | 4.58E-01 | 0 |
| 290 | 67 | primary | AC | Astrocyte | 1.14 | 4.35E-01 | 6.06E-01 | 4.35E-01 | 0 |
| 291 | 67 | primary | AC | Endothelial | 1.00 | 1.00E+00 | 2.10E-01 | 2.10E-01 | 0 |
| 292 | 67 | primary | AC | MES | 0.00 | 1.00E+00 | 8.79E-02 | 8.79E-02 | 0 |
| 293 | 67 | primary | AC | Macrophage | 1.00 | 9.99E-01 | 5.45E-01 | 5.45E-01 | 0 |
| 294 | 67 | primary | AC | Microglia | 1.26 | 3.92E-01 | 6.10E-01 | 3.92E-01 | 0 |
| 295 | 67 | primary | AC | NK cell | 1.33 | 1.00E+00 | 9.99E-04 | 9.99E-04 | -1 |
| 296 | 67 | primary | AC | NPC | 1.00 | 9.95E-01 | 7.84E-01 | 7.84E-01 | 0 |
| 297 | 67 | primary | AC | Neuron | 1.64 | 4.90E-02 | 9.54E-01 | 4.90E-02 | 0 |
| 298 | 67 | primary | AC | OPC | 1.74 | 4.00E-03 | 9.97E-01 | 4.00E-03 | 1 |
| 299 | 67 | primary | AC | Oligodendrocyte | 1.80 | 5.99E-03 | 9.95E-01 | 5.99E-03 | 1 |
| 300 | 67 | primary | AC | T cell | 1.13 | 6.03E-01 | 4.34E-01 | 4.34E-01 | 0 |
| 301 | 67 | primary | Astrocyte | AC | 1.00 | 1.00E+00 | 8.78E-01 | 8.78E-01 | 0 |
| 302 | 67 | primary | Astrocyte | Astrocyte | 1.49 | 9.99E-04 | 1.00E+00 | 9.99E-04 | 1 |
| 303 | 67 | primary | Astrocyte | Endothelial | 1.19 | 4.70E-02 | 9.56E-01 | 4.70E-02 | 0 |
| 304 | 67 | primary | Astrocyte | MES | 1.26 | 9.99E-04 | 1.00E+00 | 9.99E-04 | 1 |
| 305 | 67 | primary | Astrocyte | Macrophage | 1.10 | 2.26E-01 | 7.86E-01 | 2.26E-01 | 0 |
| 306 | 67 | primary | Astrocyte | Microglia | 1.40 | 9.99E-04 | 1.00E+00 | 9.99E-04 | 1 |
| 307 | 67 | primary | Astrocyte | NK cell | 1.76 | 1.00E+00 | 9.99E-04 | 9.99E-04 | -1 |
| 308 | 67 | primary | Astrocyte | NPC | 1.08 | 1.54E-01 | 8.54E-01 | 1.54E-01 | 0 |
| 309 | 67 | primary | Astrocyte | Neuron | 1.91 | 9.99E-04 | 1.00E+00 | 9.99E-04 | 1 |
| 310 | 67 | primary | Astrocyte | OPC | 1.48 | 9.99E-04 | 1.00E+00 | 9.99E-04 | 1 |
| 311 | 67 | primary | Astrocyte | Oligodendrocyte | 1.48 | 1.20E-02 | 9.89E-01 | 1.20E-02 | 0 |
| 312 | 67 | primary | Astrocyte | T cell | 1.17 | 4.07E-01 | 5.94E-01 | 4.07E-01 | 0 |
| 313 | 67 | primary | Endothelial | AC | 1.00 | 1.00E+00 | 8.69E-01 | 8.69E-01 | 0 |
| 314 | 67 | primary | Endothelial | Astrocyte | 1.23 | 3.00E-03 | 9.98E-01 | 3.00E-03 | 1 |
| 315 | 67 | primary | Endothelial | Endothelial | 2.38 | 9.99E-04 | 1.00E+00 | 9.99E-04 | 1 |
| 316 | 67 | primary | Endothelial | MES | 1.00 | 1.00E+00 | 5.87E-01 | 5.87E-01 | 0 |
| 317 | 67 | primary | Endothelial | Macrophage | 1.10 | 2.01E-01 | 8.06E-01 | 2.01E-01 | 0 |
| 318 | 67 | primary | Endothelial | Microglia | 1.23 | 6.12E-01 | 3.95E-01 | 3.95E-01 | 0 |
| 319 | 67 | primary | Endothelial | NK cell | 3.68 | 9.99E-04 | 1.00E+00 | 9.99E-04 | 1 |
| 320 | 67 | primary | Endothelial | NPC | 1.00 | 1.00E+00 | 5.79E-02 | 5.79E-02 | 0 |

| row | patient | surgery | from cell type | to cell tye | observed count | permutations (greater than) | permutations (less than) | p value | interaction significance/direction |
|---|---|---|---|---|---|---|---|---|---|
| 321 | 67 | primary | Endothelial | Neuron | 1.54 | 6.99E-03 | 9.94E-01 | 6.99E-03 | 1 |
| 322 | 67 | primary | Endothelial | OPC | 1.44 | 9.99E-04 | 1.00E+00 | 9.99E-04 | 1 |
| 323 | 67 | primary | Endothelial | Oligodendrocyte | 1.35 | 9.25E-01 | 7.59E-02 | 7.59E-02 | 0 |
| 324 | 67 | primary | Endothelial | T cell | 1.17 | 3.76E-01 | 6.25E-01 | 3.76E-01 | 0 |
| 325 | 67 | primary | MES | AC | 0.00 | 1.00E+00 | 8.79E-02 | 8.79E-02 | 0 |
| 326 | 67 | primary | MES | Astrocyte | 1.55 | 2.00E-03 | 9.99E-01 | 2.00E-03 | 1 |
| 327 | 67 | primary | MES | Endothelial | 1.50 | 9.99E-04 | 1.00E+00 | 9.99E-04 | 1 |
| 328 | 67 | primary | MES | MES | 1.68 | 3.00E-03 | 9.98E-01 | 3.00E-03 | 1 |
| 329 | 67 | primary | MES | Macrophage | 1.27 | 2.60E-02 | 9.78E-01 | 2.60E-02 | 0 |
| 330 | 67 | primary | MES | Microglia | 1.00 | 1.00E+00 | 9.99E-04 | 9.99E-04 | -1 |
| 331 | 67 | primary | MES | NK cell | 3.20 | 9.99E-04 | 1.00E+00 | 9.99E-04 | 1 |
| 332 | 67 | primary | MES | NPC | 0.00 | 1.00E+00 | 9.99E-04 | 9.99E-04 | -1 |
| 333 | 67 | primary | MES | Neuron | 1.00 | 1.00E+00 | 9.99E-04 | 9.99E-04 | -1 |
| 334 | 67 | primary | MES | OPC | 0.00 | 1.00E+00 | 9.99E-04 | 9.99E-04 | -1 |
| 335 | 67 | primary | MES | Oligodendrocyte | 1.29 | 9.11E-01 | 9.29E-02 | 9.29E-02 | 0 |
| 336 | 67 | primary | MES | T cell | 1.32 | 4.50E-02 | 9.57E-01 | 4.50E-02 | 0 |
| 337 | 67 | primary | Macrophage | AC | 1.00 | 9.99E-01 | 9.13E-01 | 9.13E-01 | 0 |
| 338 | 67 | primary | Macrophage | Astrocyte | 1.33 | 9.99E-04 | 1.00E+00 | 9.99E-04 | 1 |
| 339 | 67 | primary | Macrophage | Endothelial | 1.39 | 9.99E-04 | 1.00E+00 | 9.99E-04 | 1 |
| 340 | 67 | primary | Macrophage | MES | 1.27 | 2.00E-03 | 9.99E-01 | 2.00E-03 | 1 |
| 341 | 67 | primary | Macrophage | Macrophage | 1.04 | 7.80E-01 | 2.21E-01 | 2.21E-01 | 0 |
| 342 | 67 | primary | Macrophage | Microglia | 1.40 | 9.99E-04 | 1.00E+00 | 9.99E-04 | 1 |
| 343 | 67 | primary | Macrophage | NK cell | 2.28 | 9.99E-04 | 1.00E+00 | 9.99E-04 | 1 |
| 344 | 67 | primary | Macrophage | NPC | 1.19 | 6.99E-03 | 9.95E-01 | 6.99E-03 | 1 |
| 345 | 67 | primary | Macrophage | Neuron | 1.69 | 9.99E-04 | 1.00E+00 | 9.99E-04 | 1 |
| 346 | 67 | primary | Macrophage | OPC | 1.48 | 9.99E-04 | 1.00E+00 | 9.99E-04 | 1 |
| 347 | 67 | primary | Macrophage | Oligodendrocyte | 1.67 | 9.99E-04 | 1.00E+00 | 9.99E-04 | 1 |
| 348 | 67 | primary | Macrophage | T cell | 1.25 | 2.70E-02 | 9.74E-01 | 2.70E-02 | 0 |
| 349 | 67 | primary | Microglia | AC | 1.00 | 1.00E+00 | 7.64E-01 | 7.64E-01 | 0 |
| 350 | 67 | primary | Microglia | Astrocyte | 1.29 | 9.99E-04 | 1.00E+00 | 9.99E-04 | 1 |
| 351 | 67 | primary | Microglia | Endothelial | 1.26 | 9.99E-04 | 1.00E+00 | 9.99E-04 | 1 |
| 352 | 67 | primary | Microglia | MES | 1.25 | 9.99E-04 | 1.00E+00 | 9.99E-04 | 1 |
| 353 | 67 | primary | Microglia | Macrophage | 1.15 | 3.00E-03 | 9.98E-01 | 3.00E-03 | 1 |
| 354 | 67 | primary | Microglia | Microglia | 1.47 | 9.99E-04 | 1.00E+00 | 9.99E-04 | 1 |
| 355 | 67 | primary | Microglia | NK cell | 1.53 | 1.00E+00 | 9.99E-04 | 9.99E-04 | -1 |
| 356 | 67 | primary | Microglia | NPC | 1.06 | 3.52E-01 | 6.62E-01 | 3.52E-01 | 0 |
| 357 | 67 | primary | Microglia | Neuron | 1.71 | 9.99E-04 | 1.00E+00 | 9.99E-04 | 1 |
| 358 | 67 | primary | Microglia | OPC | 1.55 | 9.99E-04 | 1.00E+00 | 9.99E-04 | 1 |
| 359 | 67 | primary | Microglia | Oligodendrocyte | 1.66 | 9.99E-04 | 1.00E+00 | 9.99E-04 | 1 |
| 360 | 67 | primary | Microglia | T cell | 1.33 | 9.99E-04 | 1.00E+00 | 9.99E-04 | 1 |
| 361 | 67 | primary | NK cell | AC | 1.04 | 5.00E-02 | 9.54E-01 | 5.00E-02 | 0 |
| 362 | 67 | primary | NK cell | Astrocyte | 1.37 | 9.99E-04 | 1.00E+00 | 9.99E-04 | 1 |
| 363 | 67 | primary | NK cell | Endothelial | 1.73 | 9.99E-04 | 1.00E+00 | 9.99E-04 | 1 |
| 364 | 67 | primary | NK cell | MES | 1.38 | 9.99E-04 | 1.00E+00 | 9.99E-04 | 1 |
| 365 | 67 | primary | NK cell | Macrophage | 1.09 | 2.61E-01 | 7.40E-01 | 2.61E-01 | 0 |
| 366 | 67 | primary | NK cell | Microglia | 1.34 | 9.99E-04 | 1.00E+00 | 9.99E-04 | 1 |
| 367 | 67 | primary | NK cell | NK cell | 4.32 | 9.99E-04 | 1.00E+00 | 9.99E-04 | 1 |
| 368 | 67 | primary | NK cell | NPC | 1.13 | 9.99E-04 | 1.00E+00 | 9.99E-04 | 1 |
| 369 | 67 | primary | NK cell | Neuron | 1.61 | 9.99E-04 | 1.00E+00 | 9.99E-04 | 1 |
| 370 | 67 | primary | NK cell | OPC | 1.47 | 9.99E-04 | 1.00E+00 | 9.99E-04 | 1 |
| 371 | 67 | primary | NK cell | Oligodendrocyte | 1.56 | 9.99E-04 | 1.00E+00 | 9.99E-04 | 1 |
| 372 | 67 | primary | NK cell | T cell | 1.24 | 9.99E-04 | 1.00E+00 | 9.99E-04 | 1 |
| 373 | 67 | primary | NPC | AC | 1.00 | 9.95E-01 | 9.54E-01 | 9.54E-01 | 0 |
| 374 | 67 | primary | NPC | Astrocyte | 1.26 | 2.00E-02 | 9.81E-01 | 2.00E-02 | 0 |
| 375 | 67 | primary | NPC | Endothelial | 1.42 | 9.99E-04 | 1.00E+00 | 9.99E-04 | 1 |
| 376 | 67 | primary | NPC | MES | 0.00 | 1.00E+00 | 9.99E-04 | 9.99E-04 | -1 |
| 377 | 67 | primary | NPC | Macrophage | 1.19 | 2.70E-02 | 9.74E-01 | 2.70E-02 | 0 |
| 378 | 67 | primary | NPC | Microglia | 1.25 | 4.00E-01 | 6.03E-01 | 4.00E-01 | 0 |
| 379 | 67 | primary | NPC | NK cell | 1.28 | 1.00E+00 | 9.99E-04 | 9.99E-04 | -1 |
| 380 | 67 | primary | NPC | NPC | 1.12 | 1.54E-01 | 8.47E-01 | 1.54E-01 | 0 |
| 381 | 67 | primary | NPC | Neuron | 1.78 | 9.99E-04 | 1.00E+00 | 9.99E-04 | 1 |
| 382 | 67 | primary | NPC | OPC | 1.67 | 9.99E-04 | 1.00E+00 | 9.99E-04 | 1 |
| 383 | 67 | primary | NPC | Oligodendrocyte | 1.78 | 9.99E-04 | 1.00E+00 | 9.99E-04 | 1 |
| 384 | 67 | primary | NPC | T cell | 1.16 | 5.14E-01 | 4.90E-01 | 4.90E-01 | 0 |
| 385 | 67 | primary | Neuron | AC | 1.07 | 1.80E-02 | 9.83E-01 | 1.80E-02 | 0 |
| 386 | 67 | primary | Neuron | Astrocyte | 1.35 | 9.99E-04 | 1.00E+00 | 9.99E-04 | 1 |
| 387 | 67 | primary | Neuron | Endothelial | 1.30 | 9.99E-04 | 1.00E+00 | 9.99E-04 | 1 |
| 388 | 67 | primary | Neuron | MES | 1.00 | 1.00E+00 | 1.75E-01 | 1.75E-01 | 0 |
| 389 | 67 | primary | Neuron | Macrophage | 1.15 | 9.99E-04 | 1.00E+00 | 9.99E-04 | 1 |
| 390 | 67 | primary | Neuron | Microglia | 1.32 | 9.99E-04 | 1.00E+00 | 9.99E-04 | 1 |
| 391 | 67 | primary | Neuron | NK cell | 1.29 | 1.00E+00 | 9.99E-04 | 9.99E-04 | -1 |
| 392 | 67 | primary | Neuron | NPC | 1.15 | 9.99E-04 | 1.00E+00 | 9.99E-04 | 1 |
| 393 | 67 | primary | Neuron | Neuron | 1.98 | 9.99E-04 | 1.00E+00 | 9.99E-04 | 1 |
| 394 | 67 | primary | Neuron | OPC | 1.53 | 9.99E-04 | 1.00E+00 | 9.99E-04 | 1 |
| 395 | 67 | primary | Neuron | Oligodendrocyte | 1.70 | 9.99E-04 | 1.00E+00 | 9.99E-04 | 1 |
| 396 | 67 | primary | Neuron | T cell | 1.22 | 2.00E-03 | 9.99E-01 | 2.00E-03 | 1 |
| 397 | 67 | primary | OPC | AC | 1.02 | 2.56E-01 | 7.45E-01 | 2.56E-01 | 0 |
| 398 | 67 | primary | OPC | Astrocyte | 1.34 | 9.99E-04 | 1.00E+00 | 9.99E-04 | 1 |
| 399 | 67 | primary | OPC | Endothelial | 1.17 | 8.79E-02 | 9.14E-01 | 8.79E-02 | 0 |
| 400 | 67 | primary | OPC | MES | 0.00 | 1.00E+00 | 9.99E-04 | 9.99E-04 | -1 |
| 401 | 67 | primary | OPC | Macrophage | 1.12 | 4.50E-02 | 9.56E-01 | 4.50E-02 | 0 |
| 402 | 67 | primary | OPC | Microglia | 1.37 | 9.99E-04 | 1.00E+00 | 9.99E-04 | 1 |
| 403 | 67 | primary | OPC | NK cell | 1.29 | 1.00E+00 | 9.99E-04 | 9.99E-04 | -1 |
| 404 | 67 | primary | OPC | NPC | 1.10 | 1.50E-02 | 9.86E-01 | 1.50E-02 | 0 |
| 405 | 67 | primary | OPC | Neuron | 1.76 | 9.99E-04 | 1.00E+00 | 9.99E-04 | 1 |
| 406 | 67 | primary | OPC | OPC | 1.75 | 9.99E-04 | 1.00E+00 | 9.99E-04 | 1 |
| 407 | 67 | primary | OPC | Oligodendrocyte | 1.70 | 9.99E-04 | 1.00E+00 | 9.99E-04 | 1 |
| 408 | 67 | primary | OPC | T cell | 1.24 | 9.99E-04 | 1.00E+00 | 9.99E-04 | 1 |
| 409 | 67 | primary | Oligodendrocyte | AC | 1.05 | 6.39E-02 | 9.40E-01 | 6.39E-02 | 0 |
| 410 | 67 | primary | Oligodendrocyte | Astrocyte | 1.28 | 9.99E-04 | 1.00E+00 | 9.99E-04 | 1 |
| 411 | 67 | primary | Oligodendrocyte | Endothelial | 1.25 | 9.99E-04 | 1.00E+00 | 9.99E-04 | 1 |
| 412 | 67 | primary | Oligodendrocyte | MES | 1.50 | 9.99E-04 | 1.00E+00 | 9.99E-04 | 1 |
| 413 | 67 | primary | Oligodendrocyte | Macrophage | 1.14 | 9.99E-04 | 1.00E+00 | 9.99E-04 | 1 |
| 414 | 67 | primary | Oligodendrocyte | Microglia | 1.34 | 9.99E-04 | 1.00E+00 | 9.99E-04 | 1 |
| 415 | 67 | primary | Oligodendrocyte | NK cell | 1.37 | 1.00E+00 | 9.99E-04 | 9.99E-04 | -1 |
| 416 | 67 | primary | Oligodendrocyte | NPC | 1.11 | 2.00E-03 | 9.99E-01 | 2.00E-03 | 1 |
| 417 | 67 | primary | Oligodendrocyte | Neuron | 1.78 | 9.99E-04 | 1.00E+00 | 9.99E-04 | 1 |
| 418 | 67 | primary | Oligodendrocyte | OPC | 1.57 | 9.99E-04 | 1.00E+00 | 9.99E-04 | 1 |
| 419 | 67 | primary | Oligodendrocyte | Oligodendrocyte | 1.91 | 9.99E-04 | 1.00E+00 | 9.99E-04 | 1 |
| 420 | 67 | primary | Oligodendrocyte | T cell | 1.25 | 9.99E-04 | 1.00E+00 | 9.99E-04 | 1 |
| 421 | 67 | primary | T cell | AC | 1.06 | 1.06E-01 | 9.00E-01 | 1.06E-01 | 0 |
| 422 | 67 | primary | T cell | Astrocyte | 1.32 | 9.99E-04 | 1.00E+00 | 9.99E-04 | 1 |
| 423 | 67 | primary | T cell | Endothelial | 1.34 | 9.99E-04 | 1.00E+00 | 9.99E-04 | 1 |
| 424 | 67 | primary | T cell | MES | 1.57 | 9.99E-04 | 1.00E+00 | 9.99E-04 | 1 |
| 425 | 67 | primary | T cell | Macrophage | 1.12 | 6.89E-02 | 9.32E-01 | 6.89E-02 | 0 |
| 426 | 67 | primary | T cell | Microglia | 1.38 | 9.99E-04 | 1.00E+00 | 9.99E-04 | 1 |
| 427 | 67 | primary | T cell | NK cell | 2.11 | 4.80E-02 | 9.53E-01 | 4.80E-02 | 0 |
| 428 | 67 | primary | T cell | NPC | 1.09 | 5.79E-02 | 9.44E-01 | 5.79E-02 | 0 |
| 429 | 67 | primary | T cell | Neuron | 1.74 | 9.99E-04 | 1.00E+00 | 9.99E-04 | 1 |
| 430 | 67 | primary | T cell | OPC | 1.54 | 9.99E-04 | 1.00E+00 | 9.99E-04 | 1 |
| 431 | 67 | primary | T cell | Oligodendrocyte | 1.71 | 9.99E-04 | 1.00E+00 | 9.99E-04 | 1 |

| row | patient | surgery | from cell type | to cell tye | observed count | permutations (greater than) | permutations (less than) | p value | interaction significance/direction |
|---|---|---|---|---|---|---|---|---|---|
| 432 | 67 | primary | T cell | T cell | 1.34 | 9.99E-04 | 1.00E+00 | 9.99E-04 | 1 |
| 433 | 67 | recurrent | AC | AC | | | | | |
| 434 | 67 | recurrent | AC | Astrocyte | | | | | |
| 435 | 67 | recurrent | AC | Endothelial | | | | | |
| 436 | 67 | recurrent | AC | MES | | | | | |
| 437 | 67 | recurrent | AC | Macrophage | | | | | |
| 438 | 67 | recurrent | AC | Microglia | | | | | |
| 439 | 67 | recurrent | AC | NK cell | | | | | |
| 440 | 67 | recurrent | AC | NPC | | | | | |
| 441 | 67 | recurrent | AC | Neuron | | | | | |
| 442 | 67 | recurrent | AC | OPC | | | | | |
| 443 | 67 | recurrent | AC | Oligodendrocyte | | | | | |
| 444 | 67 | recurrent | AC | T cell | | | | | |
| 445 | 67 | recurrent | Astrocyte | AC | | | | | |
| 446 | 67 | recurrent | Astrocyte | Astrocyte | 3.83 | 9.99E-04 | 1.00E+00 | 9.99E-04 | 1 |
| 447 | 67 | recurrent | Astrocyte | Endothelial | 1.37 | 9.99E-04 | 1.00E+00 | 9.99E-04 | 1 |
| 448 | 67 | recurrent | Astrocyte | MES | 1.05 | 7.29E-02 | 9.28E-01 | 7.29E-02 | 0 |
| 449 | 67 | recurrent | Astrocyte | Macrophage | 1.25 | 2.00E-03 | 9.99E-01 | 2.00E-03 | 1 |
| 450 | 67 | recurrent | Astrocyte | Microglia | 1.30 | 9.99E-04 | 1.00E+00 | 9.99E-04 | 1 |
| 451 | 67 | recurrent | Astrocyte | NK cell | 1.27 | 9.99E-04 | 1.00E+00 | 9.99E-04 | 1 |
| 452 | 67 | recurrent | Astrocyte | NPC | 1.06 | 4.00E-03 | 9.97E-01 | 4.00E-03 | 1 |
| 453 | 67 | recurrent | Astrocyte | Neuron | 1.16 | 9.99E-04 | 1.00E+00 | 9.99E-04 | 1 |
| 454 | 67 | recurrent | Astrocyte | OPC | 1.07 | 4.00E-03 | 9.98E-01 | 4.00E-03 | 1 |
| 455 | 67 | recurrent | Astrocyte | Oligodendrocyte | 1.20 | 9.99E-04 | 1.00E+00 | 9.99E-04 | 1 |
| 456 | 67 | recurrent | Astrocyte | T cell | 1.20 | 9.99E-04 | 1.00E+00 | 9.99E-04 | 1 |
| 457 | 67 | recurrent | Endothelial | AC | | | | | |
| 458 | 67 | recurrent | Endothelial | Astrocyte | 2.10 | 1.00E+00 | 9.99E-04 | 9.99E-04 | -1 |
| 459 | 67 | recurrent | Endothelial | Endothelial | 3.57 | 9.99E-04 | 1.00E+00 | 9.99E-04 | 1 |
| 460 | 67 | recurrent | Endothelial | MES | 1.12 | 9.99E-04 | 1.00E+00 | 9.99E-04 | 1 |
| 461 | 67 | recurrent | Endothelial | Macrophage | 1.27 | 8.99E-03 | 9.92E-01 | 8.99E-03 | 1 |
| 462 | 67 | recurrent | Endothelial | Microglia | 1.38 | 9.99E-04 | 1.00E+00 | 9.99E-04 | 1 |
| 463 | 67 | recurrent | Endothelial | NK cell | 1.34 | 9.99E-04 | 1.00E+00 | 9.99E-04 | 1 |
| 464 | 67 | recurrent | Endothelial | NPC | 1.20 | 9.99E-04 | 1.00E+00 | 9.99E-04 | 1 |
| 465 | 67 | recurrent | Endothelial | Neuron | 1.12 | 1.56E-01 | 8.45E-01 | 1.56E-01 | 0 |
| 466 | 67 | recurrent | Endothelial | OPC | 1.00 | 9.99E-01 | 9.83E-01 | 9.83E-01 | 0 |
| 467 | 67 | recurrent | Endothelial | Oligodendrocyte | 1.14 | 5.00E-03 | 9.97E-01 | 5.00E-03 | 1 |
| 468 | 67 | recurrent | Endothelial | T cell | 1.20 | 5.49E-02 | 9.46E-01 | 5.49E-02 | 0 |
| 469 | 67 | recurrent | MES | AC | | | | | |
| 470 | 67 | recurrent | MES | Astrocyte | 3.42 | 3.00E-02 | 9.71E-01 | 3.00E-02 | 0 |
| 471 | 67 | recurrent | MES | Endothelial | 1.93 | 9.99E-04 | 1.00E+00 | 9.99E-04 | 1 |
| 472 | 67 | recurrent | MES | MES | 1.10 | 1.29E-01 | 8.72E-01 | 1.29E-01 | 0 |
| 473 | 67 | recurrent | MES | Macrophage | 1.27 | 1.48E-01 | 8.56E-01 | 1.48E-01 | 0 |
| 474 | 67 | recurrent | MES | Microglia | 1.31 | 1.74E-01 | 8.28E-01 | 1.74E-01 | 0 |
| 475 | 67 | recurrent | MES | NK cell | 1.38 | 6.99E-03 | 9.94E-01 | 6.99E-03 | 1 |
| 476 | 67 | recurrent | MES | NPC | 1.00 | 9.98E-01 | 8.90E-01 | 8.90E-01 | 0 |
| 477 | 67 | recurrent | MES | Neuron | 1.19 | 7.79E-02 | 9.26E-01 | 7.79E-02 | 0 |
| 478 | 67 | recurrent | MES | OPC | 0.00 | 1.00E+00 | 3.37E-01 | 3.37E-01 | 0 |
| 479 | 67 | recurrent | MES | Oligodendrocyte | 1.25 | 8.99E-03 | 9.96E-01 | 8.99E-03 | 1 |
| 480 | 67 | recurrent | MES | T cell | 1.33 | 1.80E-02 | 9.83E-01 | 1.80E-02 | 0 |
| 481 | 67 | recurrent | Macrophage | AC | | | | | |
| 482 | 67 | recurrent | Macrophage | Astrocyte | 2.95 | 1.00E+00 | 9.99E-04 | 9.99E-04 | -1 |
| 483 | 67 | recurrent | Macrophage | Endothelial | 1.65 | 9.99E-04 | 1.00E+00 | 9.99E-04 | 1 |
| 484 | 67 | recurrent | Macrophage | MES | 1.10 | 1.60E-02 | 9.85E-01 | 1.60E-02 | 0 |
| 485 | 67 | recurrent | Macrophage | Macrophage | 1.60 | 9.99E-04 | 1.00E+00 | 9.99E-04 | 1 |
| 486 | 67 | recurrent | Macrophage | Microglia | 1.56 | 9.99E-04 | 1.00E+00 | 9.99E-04 | 1 |
| 487 | 67 | recurrent | Macrophage | NK cell | 1.40 | 9.99E-04 | 1.00E+00 | 9.99E-04 | 1 |
| 488 | 67 | recurrent | Macrophage | NPC | 1.00 | 1.00E+00 | 6.29E-01 | 6.29E-01 | 0 |
| 489 | 67 | recurrent | Macrophage | Neuron | 1.15 | 3.80E-02 | 9.64E-01 | 3.80E-02 | 0 |
| 490 | 67 | recurrent | Macrophage | OPC | 1.00 | 9.98E-01 | 9.87E-01 | 9.87E-01 | 0 |
| 491 | 67 | recurrent | Macrophage | Oligodendrocyte | 1.15 | 2.00E-03 | 9.99E-01 | 2.00E-03 | 1 |
| 492 | 67 | recurrent | Macrophage | T cell | 1.19 | 8.99E-02 | 9.12E-01 | 8.99E-02 | 0 |
| 493 | 67 | recurrent | Microglia | AC | | | | | |
| 494 | 67 | recurrent | Microglia | Astrocyte | 2.87 | 1.00E+00 | 9.99E-04 | 9.99E-04 | -1 |
| 495 | 67 | recurrent | Microglia | Endothelial | 1.74 | 9.99E-04 | 1.00E+00 | 9.99E-04 | 1 |
| 496 | 67 | recurrent | Microglia | MES | 1.06 | 1.90E-01 | 8.11E-01 | 1.90E-01 | 0 |
| 497 | 67 | recurrent | Microglia | Macrophage | 1.41 | 9.99E-04 | 1.00E+00 | 9.99E-04 | 1 |
| 498 | 67 | recurrent | Microglia | Microglia | 1.64 | 9.99E-04 | 1.00E+00 | 9.99E-04 | 1 |
| 499 | 67 | recurrent | Microglia | NK cell | 1.41 | 9.99E-04 | 1.00E+00 | 9.99E-04 | 1 |
| 500 | 67 | recurrent | Microglia | NPC | 1.18 | 9.99E-04 | 1.00E+00 | 9.99E-04 | 1 |
| 501 | 67 | recurrent | Microglia | Neuron | 1.13 | 8.39E-02 | 9.17E-01 | 8.39E-02 | 0 |
| 502 | 67 | recurrent | Microglia | OPC | 1.33 | 9.99E-04 | 1.00E+00 | 9.99E-04 | 1 |
| 503 | 67 | recurrent | Microglia | Oligodendrocyte | 1.30 | 9.99E-04 | 1.00E+00 | 9.99E-04 | 1 |
| 504 | 67 | recurrent | Microglia | T cell | 1.29 | 9.99E-04 | 1.00E+00 | 9.99E-04 | 1 |
| 505 | 67 | recurrent | NK cell | AC | | | | | |
| 506 | 67 | recurrent | NK cell | Astrocyte | 2.99 | 1.00E+00 | 9.99E-04 | 9.99E-04 | -1 |
| 507 | 67 | recurrent | NK cell | Endothelial | 1.75 | 9.99E-04 | 1.00E+00 | 9.99E-04 | 1 |
| 508 | 67 | recurrent | NK cell | MES | 1.06 | 2.11E-01 | 7.95E-01 | 2.11E-01 | 0 |
| 509 | 67 | recurrent | NK cell | Macrophage | 1.41 | 9.99E-04 | 1.00E+00 | 9.99E-04 | 1 |
| 510 | 67 | recurrent | NK cell | Microglia | 1.48 | 9.99E-04 | 1.00E+00 | 9.99E-04 | 1 |
| 511 | 67 | recurrent | NK cell | NK cell | 1.61 | 9.99E-04 | 1.00E+00 | 9.99E-04 | 1 |
| 512 | 67 | recurrent | NK cell | NPC | 1.00 | 1.00E+00 | 6.12E-01 | 6.12E-01 | 0 |
| 513 | 67 | recurrent | NK cell | Neuron | 1.07 | 7.96E-01 | 2.05E-01 | 2.05E-01 | 0 |
| 514 | 67 | recurrent | NK cell | OPC | 1.00 | 9.97E-01 | 9.80E-01 | 9.80E-01 | 0 |
| 515 | 67 | recurrent | NK cell | Oligodendrocyte | 1.26 | 9.99E-04 | 1.00E+00 | 9.99E-04 | 1 |
| 516 | 67 | recurrent | NK cell | T cell | 1.25 | 2.00E-03 | 9.99E-01 | 2.00E-03 | 1 |
| 517 | 67 | recurrent | NPC | AC | | | | | |
| 518 | 67 | recurrent | NPC | Astrocyte | 3.68 | 9.99E-04 | 1.00E+00 | 9.99E-04 | 1 |
| 519 | 67 | recurrent | NPC | Endothelial | 1.00 | 1.00E+00 | 3.00E-03 | 3.00E-03 | -1 |
| 520 | 67 | recurrent | NPC | MES | 1.00 | 9.98E-01 | 8.27E-01 | 8.27E-01 | 0 |
| 521 | 67 | recurrent | NPC | Macrophage | 1.28 | 2.05E-01 | 7.99E-01 | 2.05E-01 | 0 |
| 522 | 67 | recurrent | NPC | Microglia | 1.39 | 7.49E-02 | 9.31E-01 | 7.49E-02 | 0 |
| 523 | 67 | recurrent | NPC | NK cell | 1.14 | 7.22E-01 | 3.07E-01 | 3.07E-01 | 0 |
| 524 | 67 | recurrent | NPC | NPC | 1.00 | 7.44E-01 | 9.71E-01 | 7.44E-01 | 0 |
| 525 | 67 | recurrent | NPC | Neuron | 1.30 | 2.60E-02 | 9.80E-01 | 2.60E-02 | 0 |
| 526 | 67 | recurrent | NPC | OPC | 1.00 | 4.24E-01 | 9.99E-01 | 4.24E-01 | 0 |
| 527 | 67 | recurrent | NPC | Oligodendrocyte | 1.25 | 5.00E-02 | 9.67E-01 | 5.00E-02 | 0 |
| 528 | 67 | recurrent | NPC | T cell | 1.25 | 1.75E-01 | 8.48E-01 | 1.75E-01 | 0 |
| 529 | 67 | recurrent | Neuron | AC | | | | | |
| 530 | 67 | recurrent | Neuron | Astrocyte | 3.58 | 9.99E-04 | 1.00E+00 | 9.99E-04 | 1 |
| 531 | 67 | recurrent | Neuron | Endothelial | 1.56 | 9.99E-04 | 1.00E+00 | 9.99E-04 | 1 |
| 532 | 67 | recurrent | Neuron | MES | 1.12 | 3.10E-02 | 9.72E-01 | 3.10E-02 | 0 |
| 533 | 67 | recurrent | Neuron | Macrophage | 1.32 | 7.99E-03 | 9.93E-01 | 7.99E-03 | 1 |
| 534 | 67 | recurrent | Neuron | Microglia | 1.25 | 4.82E-01 | 5.19E-01 | 4.82E-01 | 0 |
| 535 | 67 | recurrent | Neuron | NK cell | 1.18 | 6.12E-01 | 3.89E-01 | 3.89E-01 | 0 |
| 536 | 67 | recurrent | Neuron | NPC | 1.00 | 1.00E+00 | 7.72E-01 | 7.72E-01 | 0 |
| 537 | 67 | recurrent | Neuron | Neuron | 1.18 | 6.89E-02 | 9.36E-01 | 6.89E-02 | 0 |
| 538 | 67 | recurrent | Neuron | OPC | 1.00 | 9.36E-01 | 9.89E-01 | 9.36E-01 | 0 |
| 539 | 67 | recurrent | Neuron | Oligodendrocyte | 1.15 | 1.30E-02 | 9.90E-01 | 1.30E-02 | 0 |
| 540 | 67 | recurrent | Neuron | T cell | 1.18 | 3.12E-01 | 6.92E-01 | 3.12E-01 | 0 |
| 541 | 67 | recurrent | OPC | AC | | | | | |
| 542 | 67 | recurrent | OPC | Astrocyte | 2.67 | 9.29E-01 | 7.59E-02 | 7.59E-02 | 0 |

| row | patient | surgery | from cell type | to cell tye | observed count | permutations (greater than) | permutations (less than) | p value | interaction significance/direction |
|---|---|---|---|---|---|---|---|---|---|
| 543 | 67 | recurrent | OPC | Endothelial | 3.50 | 9.99E-04 | 1.00E+00 | 9.99E-04 | 1 |
| 544 | 67 | recurrent | OPC | MES | 0.00 | 1.00E+00 | 3.37E-01 | 3.37E-01 | 0 |
| 545 | 67 | recurrent | OPC | Macrophage | 1.00 | 9.98E-01 | 4.19E-01 | 4.19E-01 | 0 |
| 546 | 67 | recurrent | OPC | Microglia | 1.00 | 9.98E-01 | 2.88E-01 | 2.88E-01 | 0 |
| 547 | 67 | recurrent | OPC | NK cell | 1.50 | 1.51E-01 | 9.24E-01 | 1.51E-01 | 0 |
| 548 | 67 | recurrent | OPC | NPC | 1.00 | 4.24E-01 | 9.92E-01 | 4.24E-01 | 0 |
| 549 | 67 | recurrent | OPC | Neuron | 1.33 | 1.39E-01 | 9.12E-01 | 1.39E-01 | 0 |
| 550 | 67 | recurrent | OPC | OPC | 0.00 | 1.00E+00 | 9.59E-01 | 9.59E-01 | 0 |
| 551 | 67 | recurrent | OPC | Oligodendrocyte | 0.00 | 1.00E+00 | 1.74E-01 | 1.74E-01 | 0 |
| 552 | 67 | recurrent | OPC | T cell | 1.20 | 3.66E-01 | 6.97E-01 | 3.66E-01 | 0 |
| 553 | 67 | recurrent | Oligodendrocyte | AC | | | | | |
| 554 | 67 | recurrent | Oligodendrocyte | Astrocyte | 3.25 | 2.64E-01 | 7.37E-01 | 2.64E-01 | 0 |
| 555 | 67 | recurrent | Oligodendrocyte | Endothelial | 1.90 | 9.99E-04 | 1.00E+00 | 9.99E-04 | 1 |
| 556 | 67 | recurrent | Oligodendrocyte | MES | 1.00 | 1.00E+00 | 5.24E-01 | 5.24E-01 | 0 |
| 557 | 67 | recurrent | Oligodendrocyte | Macrophage | 1.23 | 2.71E-01 | 7.30E-01 | 2.71E-01 | 0 |
| 558 | 67 | recurrent | Oligodendrocyte | Microglia | 1.36 | 3.70E-02 | 9.64E-01 | 3.70E-02 | 0 |
| 559 | 67 | recurrent | Oligodendrocyte | NK cell | 1.32 | 1.30E-02 | 9.88E-01 | 1.30E-02 | 0 |
| 560 | 67 | recurrent | Oligodendrocyte | NPC | 1.00 | 1.00E+00 | 8.61E-01 | 8.61E-01 | 0 |
| 561 | 67 | recurrent | Oligodendrocyte | Neuron | 1.36 | 9.99E-04 | 1.00E+00 | 9.99E-04 | 1 |
| 562 | 67 | recurrent | Oligodendrocyte | OPC | 0.00 | 1.00E+00 | 1.74E-01 | 1.74E-01 | 0 |
| 563 | 67 | recurrent | Oligodendrocyte | Oligodendrocyte | 1.50 | 9.99E-04 | 1.00E+00 | 9.99E-04 | 1 |
| 564 | 67 | recurrent | Oligodendrocyte | T cell | 1.21 | 1.60E-01 | 8.44E-01 | 1.60E-01 | 0 |
| 565 | 67 | recurrent | T cell | AC | | | | | |
| 566 | 67 | recurrent | T cell | Astrocyte | 3.30 | 2.80E-02 | 9.73E-01 | 2.80E-02 | 0 |
| 567 | 67 | recurrent | T cell | Endothelial | 1.48 | 9.99E-04 | 1.00E+00 | 9.99E-04 | 1 |
| 568 | 67 | recurrent | T cell | MES | 1.15 | 3.00E-03 | 9.99E-01 | 3.00E-03 | 1 |
| 569 | 67 | recurrent | T cell | Macrophage | 1.30 | 9.99E-04 | 1.00E+00 | 9.99E-04 | 1 |
| 570 | 67 | recurrent | T cell | Microglia | 1.47 | 9.99E-04 | 1.00E+00 | 9.99E-04 | 1 |
| 571 | 67 | recurrent | T cell | NK cell | 1.41 | 9.99E-04 | 1.00E+00 | 9.99E-04 | 1 |
| 572 | 67 | recurrent | T cell | NPC | 1.09 | 3.20E-02 | 9.79E-01 | 3.20E-02 | 0 |
| 573 | 67 | recurrent | T cell | Neuron | 1.24 | 9.99E-04 | 1.00E+00 | 9.99E-04 | 1 |
| 574 | 67 | recurrent | T cell | OPC | 1.00 | 9.93E-01 | 9.88E-01 | 9.88E-01 | 0 |
| 575 | 67 | recurrent | T cell | Oligodendrocyte | 1.31 | 9.99E-04 | 1.00E+00 | 9.99E-04 | 1 |
| 576 | 67 | recurrent | T cell | T cell | 1.26 | 1.30E-02 | 9.88E-01 | 1.30E-02 | 0 |
| 577 | 71 | primary | AC | AC | 1.00 | 9.99E-04 | 1.00E+00 | 9.99E-04 | 1 |
| 578 | 71 | primary | AC | Astrocyte | 0.00 | 1.00E+00 | 9.48E-01 | 9.48E-01 | 0 |
| 579 | 71 | primary | AC | Endothelial | 0.00 | 1.00E+00 | 1.10E-02 | 1.10E-02 | 0 |
| 580 | 71 | primary | AC | MES | 3.00 | 3.00E-03 | 1.00E+00 | 3.00E-03 | 1 |
| 581 | 71 | primary | AC | Macrophage | 4.33 | 3.46E-01 | 7.77E-01 | 3.46E-01 | 0 |
| 582 | 71 | primary | AC | Microglia | 0.00 | 1.00E+00 | 2.55E-01 | 2.55E-01 | 0 |
| 583 | 71 | primary | AC | NK cell | 0.00 | 1.00E+00 | 9.58E-01 | 9.58E-01 | 0 |
| 584 | 71 | primary | AC | NPC | | | | | |
| 585 | 71 | primary | AC | Neuron | | | | | |
| 586 | 71 | primary | AC | OPC | | | | | |
| 587 | 71 | primary | AC | Oligodendrocyte | 0.00 | 1.00E+00 | 9.85E-01 | 9.85E-01 | 0 |
| 588 | 71 | primary | AC | T cell | 0.00 | 1.00E+00 | 7.56E-01 | 7.56E-01 | 0 |
| 589 | 71 | primary | Astrocyte | AC | 0.00 | 1.00E+00 | 9.48E-01 | 9.48E-01 | 0 |
| 590 | 71 | primary | Astrocyte | Astrocyte | 1.30 | 5.00E-03 | 9.96E-01 | 5.00E-03 | 1 |
| 591 | 71 | primary | Astrocyte | Endothelial | 1.67 | 5.71E-01 | 4.63E-01 | 4.63E-01 | 0 |
| 592 | 71 | primary | Astrocyte | MES | 1.35 | 4.10E-02 | 9.60E-01 | 4.10E-02 | 0 |
| 593 | 71 | primary | Astrocyte | Macrophage | 3.13 | 1.00E+00 | 9.99E-04 | 9.99E-04 | -1 |
| 594 | 71 | primary | Astrocyte | Microglia | 1.38 | 1.07E-01 | 9.03E-01 | 1.07E-01 | 0 |
| 595 | 71 | primary | Astrocyte | NK cell | 1.00 | 4.64E-01 | 9.96E-01 | 4.64E-01 | 0 |
| 596 | 71 | primary | Astrocyte | NPC | | | | | |
| 597 | 71 | primary | Astrocyte | Neuron | | | | | |
| 598 | 71 | primary | Astrocyte | OPC | | | | | |
| 599 | 71 | primary | Astrocyte | Oligodendrocyte | 0.00 | 1.00E+00 | 7.43E-01 | 7.43E-01 | 0 |
| 600 | 71 | primary | Astrocyte | T cell | 1.00 | 9.82E-01 | 8.36E-01 | 8.36E-01 | 0 |
| 601 | 71 | primary | Endothelial | AC | 0.00 | 1.00E+00 | 1.10E-02 | 1.10E-02 | 0 |
| 602 | 71 | primary | Endothelial | Astrocyte | 1.33 | 9.99E-04 | 1.00E+00 | 9.99E-04 | 1 |
| 603 | 71 | primary | Endothelial | Endothelial | 3.60 | 9.99E-04 | 1.00E+00 | 9.99E-04 | 1 |
| 604 | 71 | primary | Endothelial | MES | 1.19 | 9.99E-04 | 1.00E+00 | 9.99E-04 | 1 |
| 605 | 71 | primary | Endothelial | Macrophage | 2.49 | 1.00E+00 | 9.99E-04 | 9.99E-04 | -1 |
| 606 | 71 | primary | Endothelial | Microglia | 1.41 | 9.99E-04 | 1.00E+00 | 9.99E-04 | 1 |
| 607 | 71 | primary | Endothelial | NK cell | 1.00 | 1.00E+00 | 7.58E-01 | 7.58E-01 | 0 |
| 608 | 71 | primary | Endothelial | NPC | | | | | |
| 609 | 71 | primary | Endothelial | Neuron | | | | | |
| 610 | 71 | primary | Endothelial | OPC | | | | | |
| 611 | 71 | primary | Endothelial | Oligodendrocyte | 1.00 | 1.00E+00 | 9.61E-01 | 9.61E-01 | 0 |
| 612 | 71 | primary | Endothelial | T cell | 1.07 | 5.00E-02 | 9.51E-01 | 5.00E-02 | 0 |
| 613 | 71 | primary | MES | AC | 1.00 | 5.14E-01 | 1.00E+00 | 5.14E-01 | 0 |
| 614 | 71 | primary | MES | Astrocyte | 1.35 | 9.99E-04 | 1.00E+00 | 9.99E-04 | 1 |
| 615 | 71 | primary | MES | Endothelial | 1.44 | 1.00E+00 | 9.99E-04 | 9.99E-04 | -1 |
| 616 | 71 | primary | MES | MES | 1.71 | 9.99E-04 | 1.00E+00 | 9.99E-04 | 1 |
| 617 | 71 | primary | MES | Macrophage | 3.96 | 5.00E-02 | 9.51E-01 | 5.00E-02 | 0 |
| 618 | 71 | primary | MES | Microglia | 1.33 | 9.99E-04 | 1.00E+00 | 9.99E-04 | 1 |
| 619 | 71 | primary | MES | NK cell | 1.00 | 1.00E+00 | 9.50E-01 | 9.50E-01 | 0 |
| 620 | 71 | primary | MES | NPC | | | | | |
| 621 | 71 | primary | MES | Neuron | | | | | |
| 622 | 71 | primary | MES | OPC | | | | | |
| 623 | 71 | primary | MES | Oligodendrocyte | 1.00 | 9.80E-01 | 9.82E-01 | 9.80E-01 | 0 |
| 624 | 71 | primary | MES | T cell | 1.17 | 9.99E-04 | 1.00E+00 | 9.99E-04 | 1 |
| 625 | 71 | primary | Macrophage | AC | 1.18 | 3.00E-03 | 9.99E-01 | 3.00E-03 | 1 |
| 626 | 71 | primary | Macrophage | Astrocyte | 1.21 | 9.99E-04 | 1.00E+00 | 9.99E-04 | 1 |
| 627 | 71 | primary | Macrophage | Endothelial | 1.80 | 9.99E-04 | 1.00E+00 | 9.99E-04 | 1 |
| 628 | 71 | primary | Macrophage | MES | 1.31 | 9.99E-04 | 1.00E+00 | 9.99E-04 | 1 |
| 629 | 71 | primary | Macrophage | Macrophage | 4.70 | 9.99E-04 | 1.00E+00 | 9.99E-04 | 1 |
| 630 | 71 | primary | Macrophage | Microglia | 1.31 | 9.99E-04 | 1.00E+00 | 9.99E-04 | 1 |
| 631 | 71 | primary | Macrophage | NK cell | 1.05 | 9.99E-04 | 1.00E+00 | 9.99E-04 | 1 |
| 632 | 71 | primary | Macrophage | NPC | | | | | |
| 633 | 71 | primary | Macrophage | Neuron | | | | | |
| 634 | 71 | primary | Macrophage | OPC | | | | | |
| 635 | 71 | primary | Macrophage | Oligodendrocyte | 1.03 | 3.50E-02 | 9.66E-01 | 3.50E-02 | 0 |
| 636 | 71 | primary | Macrophage | T cell | 1.10 | 9.99E-04 | 1.00E+00 | 9.99E-04 | 1 |
| 637 | 71 | primary | Microglia | AC | 0.00 | 1.00E+00 | 2.55E-01 | 2.55E-01 | 0 |
| 638 | 71 | primary | Microglia | Astrocyte | 1.22 | 9.99E-04 | 1.00E+00 | 9.99E-04 | 1 |
| 639 | 71 | primary | Microglia | Endothelial | 2.57 | 9.99E-04 | 1.00E+00 | 9.99E-04 | 1 |
| 640 | 71 | primary | Microglia | MES | 1.37 | 9.99E-04 | 1.00E+00 | 9.99E-04 | 1 |
| 641 | 71 | primary | Microglia | Macrophage | 3.35 | 1.00E+00 | 9.99E-04 | 9.99E-04 | -1 |
| 642 | 71 | primary | Microglia | Microglia | 1.67 | 9.99E-04 | 1.00E+00 | 9.99E-04 | 1 |
| 643 | 71 | primary | Microglia | NK cell | 1.08 | 2.40E-02 | 9.82E-01 | 2.40E-02 | 0 |
| 644 | 71 | primary | Microglia | NPC | | | | | |
| 645 | 71 | primary | Microglia | Neuron | | | | | |
| 646 | 71 | primary | Microglia | OPC | | | | | |
| 647 | 71 | primary | Microglia | Oligodendrocyte | 1.00 | 9.99E-01 | 9.78E-01 | 9.78E-01 | 0 |
| 648 | 71 | primary | Microglia | T cell | 1.10 | 5.99E-03 | 9.95E-01 | 5.99E-03 | 1 |
| 649 | 71 | primary | NK cell | AC | 0.00 | 1.00E+00 | 9.58E-01 | 9.58E-01 | 0 |
| 650 | 71 | primary | NK cell | Astrocyte | 1.00 | 4.64E-01 | 9.96E-01 | 4.64E-01 | 0 |
| 651 | 71 | primary | NK cell | Endothelial | 2.27 | 9.99E-04 | 1.00E+00 | 9.99E-04 | 1 |
| 652 | 71 | primary | NK cell | MES | 1.50 | 1.30E-02 | 9.94E-01 | 1.30E-02 | 0 |
| 653 | 71 | primary | NK cell | Macrophage | 3.44 | 9.68E-01 | 3.30E-02 | 3.30E-02 | 0 |

| row | patient | surgery | from cell type | to cell tye | observed count | permutations (greater than) | permutations (less than) | p value | interaction significance/direction |
|---|---|---|---|---|---|---|---|---|---|
| 654 | 71 | primary | NK cell | Microglia | 1.44 | 5.29E-02 | 9.50E-01 | 5.29E-02 | 0 |
| 655 | 71 | primary | NK cell | NK cell | 1.00 | 2.40E-01 | 9.96E-01 | 2.40E-01 | 0 |
| 656 | 71 | primary | NK cell | NPC | | | | | |
| 657 | 71 | primary | NK cell | Neuron | | | | | |
| 658 | 71 | primary | NK cell | OPC | | | | | |
| 659 | 71 | primary | NK cell | Oligodendrocyte | 0.00 | 1.00E+00 | 7.53E-01 | 7.53E-01 | 0 |
| 660 | 71 | primary | NK cell | T cell | 1.00 | 9.70E-01 | 8.62E-01 | 8.62E-01 | 0 |
| 661 | 71 | primary | NPC | AC | | | | | |
| 662 | 71 | primary | NPC | Astrocyte | | | | | |
| 663 | 71 | primary | NPC | Endothelial | | | | | |
| 664 | 71 | primary | NPC | MES | | | | | |
| 665 | 71 | primary | NPC | Macrophage | | | | | |
| 666 | 71 | primary | NPC | Microglia | | | | | |
| 667 | 71 | primary | NPC | NK cell | | | | | |
| 668 | 71 | primary | NPC | NPC | | | | | |
| 669 | 71 | primary | NPC | Neuron | | | | | |
| 670 | 71 | primary | NPC | OPC | | | | | |
| 671 | 71 | primary | NPC | Oligodendrocyte | | | | | |
| 672 | 71 | primary | NPC | T cell | | | | | |
| 673 | 71 | primary | Neuron | AC | | | | | |
| 674 | 71 | primary | Neuron | Astrocyte | | | | | |
| 675 | 71 | primary | Neuron | Endothelial | | | | | |
| 676 | 71 | primary | Neuron | MES | | | | | |
| 677 | 71 | primary | Neuron | Macrophage | | | | | |
| 678 | 71 | primary | Neuron | Microglia | | | | | |
| 679 | 71 | primary | Neuron | NK cell | | | | | |
| 680 | 71 | primary | Neuron | NPC | | | | | |
| 681 | 71 | primary | Neuron | Neuron | | | | | |
| 682 | 71 | primary | Neuron | OPC | | | | | |
| 683 | 71 | primary | Neuron | Oligodendrocyte | | | | | |
| 684 | 71 | primary | Neuron | T cell | | | | | |
| 685 | 71 | primary | OPC | AC | | | | | |
| 686 | 71 | primary | OPC | Astrocyte | | | | | |
| 687 | 71 | primary | OPC | Endothelial | | | | | |
| 688 | 71 | primary | OPC | MES | | | | | |
| 689 | 71 | primary | OPC | Macrophage | | | | | |
| 690 | 71 | primary | OPC | Microglia | | | | | |
| 691 | 71 | primary | OPC | NK cell | | | | | |
| 692 | 71 | primary | OPC | NPC | | | | | |
| 693 | 71 | primary | OPC | Neuron | | | | | |
| 694 | 71 | primary | OPC | OPC | | | | | |
| 695 | 71 | primary | OPC | Oligodendrocyte | | | | | |
| 696 | 71 | primary | OPC | T cell | | | | | |
| 697 | 71 | primary | Oligodendrocyte | AC | 0.00 | 1.00E+00 | 9.85E-01 | 9.85E-01 | 0 |
| 698 | 71 | primary | Oligodendrocyte | Astrocyte | 0.00 | 1.00E+00 | 7.43E-01 | 7.43E-01 | 0 |
| 699 | 71 | primary | Oligodendrocyte | Endothelial | 1.20 | 9.89E-01 | 1.30E-02 | 1.30E-02 | 0 |
| 700 | 71 | primary | Oligodendrocyte | MES | 1.50 | 6.99E-02 | 9.75E-01 | 6.99E-02 | 0 |
| 701 | 71 | primary | Oligodendrocyte | Macrophage | 4.73 | 5.00E-03 | 9.96E-01 | 5.00E-03 | 1 |
| 702 | 71 | primary | Oligodendrocyte | Microglia | 1.33 | 2.71E-01 | 7.94E-01 | 2.71E-01 | 0 |
| 703 | 71 | primary | Oligodendrocyte | NK cell | 0.00 | 1.00E+00 | 7.53E-01 | 7.53E-01 | 0 |
| 704 | 71 | primary | Oligodendrocyte | NPC | | | | | |
| 705 | 71 | primary | Oligodendrocyte | Neuron | | | | | |
| 706 | 71 | primary | Oligodendrocyte | OPC | | | | | |
| 707 | 71 | primary | Oligodendrocyte | Oligodendrocyte | 1.00 | 5.59E-02 | 1.00E+00 | 5.59E-02 | 0 |
| 708 | 71 | primary | Oligodendrocyte | T cell | 1.00 | 7.95E-01 | 9.46E-01 | 7.95E-01 | 0 |
| 709 | 71 | primary | T cell | AC | 0.00 | 1.00E+00 | 7.56E-01 | 7.56E-01 | 0 |
| 710 | 71 | primary | T cell | Astrocyte | 1.00 | 9.82E-01 | 9.75E-01 | 9.75E-01 | 0 |
| 711 | 71 | primary | T cell | Endothelial | 2.44 | 9.99E-04 | 1.00E+00 | 9.99E-04 | 1 |
| 712 | 71 | primary | T cell | MES | 1.63 | 9.99E-04 | 1.00E+00 | 9.99E-04 | 1 |
| 713 | 71 | primary | T cell | Macrophage | 3.53 | 1.00E+00 | 9.99E-04 | 9.99E-04 | -1 |
| 714 | 71 | primary | T cell | Microglia | 1.59 | 9.99E-04 | 1.00E+00 | 9.99E-04 | 1 |
| 715 | 71 | primary | T cell | NK cell | 1.33 | 1.20E-02 | 9.93E-01 | 1.20E-02 | 0 |
| 716 | 71 | primary | T cell | NPC | | | | | |
| 717 | 71 | primary | T cell | Neuron | | | | | |
| 718 | 71 | primary | T cell | OPC | | | | | |
| 719 | 71 | primary | T cell | Oligodendrocyte | 1.00 | 7.95E-01 | 9.96E-01 | 7.95E-01 | 0 |
| 720 | 71 | primary | T cell | T cell | 1.12 | 1.23E-01 | 8.84E-01 | 1.23E-01 | 0 |
| 721 | 71 | recurrent | AC | AC | 3.24 | 9.99E-04 | 1.00E+00 | 9.99E-04 | 1 |
| 722 | 71 | recurrent | AC | Astrocyte | 1.75 | 9.99E-04 | 1.00E+00 | 9.99E-04 | 1 |
| 723 | 71 | recurrent | AC | Endothelial | 1.00 | 1.00E+00 | 9.99E-04 | 9.99E-04 | -1 |
| 724 | 71 | recurrent | AC | MES | 1.61 | 9.99E-04 | 1.00E+00 | 9.99E-04 | 1 |
| 725 | 71 | recurrent | AC | Macrophage | 1.10 | 3.65E-01 | 6.38E-01 | 3.65E-01 | 0 |
| 726 | 71 | recurrent | AC | Microglia | 1.15 | 8.25E-01 | 1.77E-01 | 1.77E-01 | 0 |
| 727 | 71 | recurrent | AC | NK cell | 1.09 | 9.96E-01 | 5.00E-03 | 5.00E-03 | -1 |
| 728 | 71 | recurrent | AC | NPC | 1.33 | 9.99E-04 | 1.00E+00 | 9.99E-04 | 1 |
| 729 | 71 | recurrent | AC | Neuron | 1.38 | 2.00E-03 | 9.99E-01 | 2.00E-03 | 1 |
| 730 | 71 | recurrent | AC | OPC | 1.21 | 5.87E-01 | 4.15E-01 | 4.15E-01 | 0 |
| 731 | 71 | recurrent | AC | Oligodendrocyte | 1.00 | 1.00E+00 | 9.99E-04 | 9.99E-04 | -1 |
| 732 | 71 | recurrent | AC | T cell | 1.26 | 1.00E+00 | 9.99E-04 | 9.99E-04 | -1 |
| 733 | 71 | recurrent | Astrocyte | AC | 2.58 | 9.99E-04 | 1.00E+00 | 9.99E-04 | 1 |
| 734 | 71 | recurrent | Astrocyte | Astrocyte | 2.05 | 9.99E-04 | 1.00E+00 | 9.99E-04 | 1 |
| 735 | 71 | recurrent | Astrocyte | Endothelial | 1.22 | 4.00E-03 | 9.98E-01 | 4.00E-03 | 1 |
| 736 | 71 | recurrent | Astrocyte | MES | 2.29 | 9.99E-04 | 1.00E+00 | 9.99E-04 | 1 |
| 737 | 71 | recurrent | Astrocyte | Macrophage | 1.12 | 1.74E-01 | 8.32E-01 | 1.74E-01 | 0 |
| 738 | 71 | recurrent | Astrocyte | Microglia | 1.22 | 1.10E-01 | 8.91E-01 | 1.10E-01 | 0 |
| 739 | 71 | recurrent | Astrocyte | NK cell | 1.12 | 8.33E-01 | 1.71E-01 | 1.71E-01 | 0 |
| 740 | 71 | recurrent | Astrocyte | NPC | 1.28 | 9.99E-04 | 1.00E+00 | 9.99E-04 | 1 |
| 741 | 71 | recurrent | Astrocyte | Neuron | 1.34 | 1.07E-01 | 8.94E-01 | 1.07E-01 | 0 |
| 742 | 71 | recurrent | Astrocyte | OPC | 1.59 | 9.99E-04 | 1.00E+00 | 9.99E-04 | 1 |
| 743 | 71 | recurrent | Astrocyte | Oligodendrocyte | 1.48 | 9.99E-04 | 1.00E+00 | 9.99E-04 | 1 |
| 744 | 71 | recurrent | Astrocyte | T cell | 1.60 | 1.42E-01 | 8.59E-01 | 1.42E-01 | 0 |
| 745 | 71 | recurrent | Endothelial | AC | 1.50 | 5.00E-03 | 9.98E-01 | 5.00E-03 | 1 |
| 746 | 71 | recurrent | Endothelial | Astrocyte | 1.10 | 9.55E-01 | 5.00E-02 | 5.00E-02 | 0 |
| 747 | 71 | recurrent | Endothelial | Endothelial | 3.09 | 9.99E-04 | 1.00E+00 | 9.99E-04 | 1 |
| 748 | 71 | recurrent | Endothelial | MES | 1.50 | 1.53E-01 | 8.58E-01 | 1.53E-01 | 0 |
| 749 | 71 | recurrent | Endothelial | Macrophage | 1.23 | 8.99E-03 | 9.92E-01 | 8.99E-03 | 1 |
| 750 | 71 | recurrent | Endothelial | Microglia | 1.65 | 9.99E-04 | 1.00E+00 | 9.99E-04 | 1 |
| 751 | 71 | recurrent | Endothelial | NK cell | 1.45 | 9.99E-04 | 1.00E+00 | 9.99E-04 | 1 |
| 752 | 71 | recurrent | Endothelial | NPC | 1.00 | 1.00E+00 | 2.10E-02 | 2.10E-02 | 0 |
| 753 | 71 | recurrent | Endothelial | Neuron | 1.50 | 9.99E-04 | 1.00E+00 | 9.99E-04 | 1 |
| 754 | 71 | recurrent | Endothelial | OPC | 1.38 | 3.00E-03 | 9.98E-01 | 3.00E-03 | 1 |
| 755 | 71 | recurrent | Endothelial | Oligodendrocyte | 1.21 | 8.29E-02 | 9.18E-01 | 8.29E-02 | 0 |
| 756 | 71 | recurrent | Endothelial | T cell | 1.56 | 4.25E-01 | 5.76E-01 | 4.25E-01 | 0 |
| 757 | 71 | recurrent | MES | AC | 1.86 | 9.99E-04 | 1.00E+00 | 9.99E-04 | 1 |
| 758 | 71 | recurrent | MES | Astrocyte | 1.55 | 9.99E-04 | 1.00E+00 | 9.99E-04 | 1 |
| 759 | 71 | recurrent | MES | Endothelial | 1.34 | 9.99E-04 | 1.00E+00 | 9.99E-04 | 1 |
| 760 | 71 | recurrent | MES | MES | 3.00 | 9.99E-04 | 1.00E+00 | 9.99E-04 | 1 |
| 761 | 71 | recurrent | MES | Macrophage | 1.22 | 9.99E-04 | 1.00E+00 | 9.99E-04 | 1 |
| 762 | 71 | recurrent | MES | Microglia | 1.40 | 9.99E-04 | 1.00E+00 | 9.99E-04 | 1 |
| 763 | 71 | recurrent | MES | NK cell | 1.27 | 9.99E-04 | 1.00E+00 | 9.99E-04 | 1 |
| 764 | 71 | recurrent | MES | NPC | 1.39 | 9.99E-04 | 1.00E+00 | 9.99E-04 | 1 |

| row | patient | surgery | from cell type | to cell tye | observed count | permutations (greater than) | permutations (less than) | p value | interaction significance/direction |
|---|---|---|---|---|---|---|---|---|---|
| 765 | 71 | recurrent | MES | Neuron | 1.56 | 9.99E-04 | 1.00E+00 | 9.99E-04 | 1 |
| 766 | 71 | recurrent | MES | OPC | 1.44 | 9.99E-04 | 1.00E+00 | 9.99E-04 | 1 |
| 767 | 71 | recurrent | MES | Oligodendrocyte | 1.64 | 9.99E-04 | 1.00E+00 | 9.99E-04 | 1 |
| 768 | 71 | recurrent | MES | T cell | 1.77 | 9.99E-04 | 1.00E+00 | 9.99E-04 | 1 |
| 769 | 71 | recurrent | Macrophage | AC | 2.61 | 9.99E-04 | 1.00E+00 | 9.99E-04 | 1 |
| 770 | 71 | recurrent | Macrophage | Astrocyte | 1.74 | 9.99E-04 | 1.00E+00 | 9.99E-04 | 1 |
| 771 | 71 | recurrent | Macrophage | Endothelial | 1.94 | 9.99E-04 | 1.00E+00 | 9.99E-04 | 1 |
| 772 | 71 | recurrent | Macrophage | MES | 1.94 | 9.99E-04 | 1.00E+00 | 9.99E-04 | 1 |
| 773 | 71 | recurrent | Macrophage | Macrophage | 1.08 | 4.41E-01 | 5.60E-01 | 4.41E-01 | 0 |
| 774 | 71 | recurrent | Macrophage | Microglia | 1.61 | 9.99E-04 | 1.00E+00 | 9.99E-04 | 1 |
| 775 | 71 | recurrent | Macrophage | NK cell | 1.44 | 9.99E-04 | 1.00E+00 | 9.99E-04 | 1 |
| 776 | 71 | recurrent | Macrophage | NPC | 1.56 | 9.99E-04 | 1.00E+00 | 9.99E-04 | 1 |
| 777 | 71 | recurrent | Macrophage | Neuron | 1.42 | 1.70E-02 | 9.84E-01 | 1.70E-02 | 0 |
| 778 | 71 | recurrent | Macrophage | OPC | 1.58 | 9.99E-04 | 1.00E+00 | 9.99E-04 | 1 |
| 779 | 71 | recurrent | Macrophage | Oligodendrocyte | 1.36 | 3.00E-03 | 9.98E-01 | 3.00E-03 | 1 |
| 780 | 71 | recurrent | Macrophage | T cell | 1.98 | 9.99E-04 | 1.00E+00 | 9.99E-04 | 1 |
| 781 | 71 | recurrent | Microglia | AC | 2.50 | 9.99E-04 | 1.00E+00 | 9.99E-04 | 1 |
| 782 | 71 | recurrent | Microglia | Astrocyte | 1.42 | 9.99E-04 | 1.00E+00 | 9.99E-04 | 1 |
| 783 | 71 | recurrent | Microglia | Endothelial | 1.70 | 9.99E-04 | 1.00E+00 | 9.99E-04 | 1 |
| 784 | 71 | recurrent | Microglia | MES | 1.85 | 9.99E-04 | 1.00E+00 | 9.99E-04 | 1 |
| 785 | 71 | recurrent | Microglia | Macrophage | 1.24 | 2.00E-03 | 9.99E-01 | 2.00E-03 | 1 |
| 786 | 71 | recurrent | Microglia | Microglia | 1.95 | 9.99E-04 | 1.00E+00 | 9.99E-04 | 1 |
| 787 | 71 | recurrent | Microglia | NK cell | 1.52 | 9.99E-04 | 1.00E+00 | 9.99E-04 | 1 |
| 788 | 71 | recurrent | Microglia | NPC | 1.29 | 9.99E-04 | 1.00E+00 | 9.99E-04 | 1 |
| 789 | 71 | recurrent | Microglia | Neuron | 1.41 | 3.00E-03 | 9.98E-01 | 3.00E-03 | 1 |
| 790 | 71 | recurrent | Microglia | OPC | 1.42 | 9.99E-04 | 1.00E+00 | 9.99E-04 | 1 |
| 791 | 71 | recurrent | Microglia | Oligodendrocyte | 1.25 | 6.99E-03 | 9.94E-01 | 6.99E-03 | 1 |
| 792 | 71 | recurrent | Microglia | T cell | 2.22 | 9.99E-04 | 1.00E+00 | 9.99E-04 | 1 |
| 793 | 71 | recurrent | NK cell | AC | 2.00 | 9.99E-04 | 1.00E+00 | 9.99E-04 | 1 |
| 794 | 71 | recurrent | NK cell | Astrocyte | 1.33 | 9.99E-04 | 1.00E+00 | 9.99E-04 | 1 |
| 795 | 71 | recurrent | NK cell | Endothelial | 1.72 | 9.99E-04 | 1.00E+00 | 9.99E-04 | 1 |
| 796 | 71 | recurrent | NK cell | MES | 1.31 | 9.99E-01 | 2.00E-03 | 2.00E-03 | -1 |
| 797 | 71 | recurrent | NK cell | Macrophage | 1.13 | 1.57E-01 | 8.47E-01 | 1.57E-01 | 0 |
| 798 | 71 | recurrent | NK cell | Microglia | 1.73 | 9.99E-04 | 1.00E+00 | 9.99E-04 | 1 |
| 799 | 71 | recurrent | NK cell | NK cell | 1.87 | 9.99E-04 | 1.00E+00 | 9.99E-04 | 1 |
| 800 | 71 | recurrent | NK cell | NPC | 1.00 | 1.00E+00 | 5.99E-03 | 5.99E-03 | -1 |
| 801 | 71 | recurrent | NK cell | Neuron | 1.43 | 2.00E-03 | 9.99E-01 | 2.00E-03 | 1 |
| 802 | 71 | recurrent | NK cell | OPC | 1.38 | 9.99E-04 | 1.00E+00 | 9.99E-04 | 1 |
| 803 | 71 | recurrent | NK cell | Oligodendrocyte | 1.30 | 9.99E-04 | 1.00E+00 | 9.99E-04 | 1 |
| 804 | 71 | recurrent | NK cell | T cell | 2.13 | 9.99E-04 | 1.00E+00 | 9.99E-04 | 1 |
| 805 | 71 | recurrent | NPC | AC | 2.34 | 9.99E-04 | 1.00E+00 | 9.99E-04 | 1 |
| 806 | 71 | recurrent | NPC | Astrocyte | 1.77 | 9.99E-04 | 1.00E+00 | 9.99E-04 | 1 |
| 807 | 71 | recurrent | NPC | Endothelial | 1.50 | 9.99E-04 | 1.00E+00 | 9.99E-04 | 1 |
| 808 | 71 | recurrent | NPC | MES | 2.30 | 9.99E-04 | 1.00E+00 | 9.99E-04 | 1 |
| 809 | 71 | recurrent | NPC | Macrophage | 1.18 | 5.49E-02 | 9.48E-01 | 5.49E-02 | 0 |
| 810 | 71 | recurrent | NPC | Microglia | 1.15 | 6.85E-01 | 3.20E-01 | 3.20E-01 | 0 |
| 811 | 71 | recurrent | NPC | NK cell | 1.14 | 5.71E-01 | 4.46E-01 | 4.46E-01 | 0 |
| 812 | 71 | recurrent | NPC | NPC | 1.55 | 9.99E-04 | 1.00E+00 | 9.99E-04 | 1 |
| 813 | 71 | recurrent | NPC | Neuron | 1.76 | 9.99E-04 | 1.00E+00 | 9.99E-04 | 1 |
| 814 | 71 | recurrent | NPC | OPC | 1.40 | 2.00E-03 | 9.99E-01 | 2.00E-03 | 1 |
| 815 | 71 | recurrent | NPC | Oligodendrocyte | 1.00 | 1.00E+00 | 9.99E-04 | 9.99E-04 | -1 |
| 816 | 71 | recurrent | NPC | T cell | 1.08 | 1.00E+00 | 9.99E-04 | 9.99E-04 | -1 |
| 817 | 71 | recurrent | Neuron | AC | 1.84 | 9.99E-04 | 1.00E+00 | 9.99E-04 | 1 |
| 818 | 71 | recurrent | Neuron | Astrocyte | 1.45 | 9.99E-04 | 1.00E+00 | 9.99E-04 | 1 |
| 819 | 71 | recurrent | Neuron | Endothelial | 1.41 | 9.99E-04 | 1.00E+00 | 9.99E-04 | 1 |
| 820 | 71 | recurrent | Neuron | MES | 1.84 | 9.99E-04 | 1.00E+00 | 9.99E-04 | 1 |
| 821 | 71 | recurrent | Neuron | Macrophage | 1.12 | 1.56E-01 | 8.46E-01 | 1.56E-01 | 0 |
| 822 | 71 | recurrent | Neuron | Microglia | 1.18 | 4.54E-01 | 5.47E-01 | 4.54E-01 | 0 |
| 823 | 71 | recurrent | Neuron | NK cell | 1.20 | 8.59E-02 | 9.19E-01 | 8.59E-02 | 0 |
| 824 | 71 | recurrent | Neuron | NPC | 1.48 | 9.99E-04 | 1.00E+00 | 9.99E-04 | 1 |
| 825 | 71 | recurrent | Neuron | Neuron | 2.15 | 9.99E-04 | 1.00E+00 | 9.99E-04 | 1 |
| 826 | 71 | recurrent | Neuron | OPC | 1.60 | 9.99E-04 | 1.00E+00 | 9.99E-04 | 1 |
| 827 | 71 | recurrent | Neuron | Oligodendrocyte | 1.49 | 9.99E-04 | 1.00E+00 | 9.99E-04 | 1 |
| 828 | 71 | recurrent | Neuron | T cell | 1.78 | 9.99E-04 | 1.00E+00 | 9.99E-04 | 1 |
| 829 | 71 | recurrent | OPC | AC | 1.81 | 9.99E-04 | 1.00E+00 | 9.99E-04 | 1 |
| 830 | 71 | recurrent | OPC | Astrocyte | 1.33 | 9.99E-04 | 1.00E+00 | 9.99E-04 | 1 |
| 831 | 71 | recurrent | OPC | Endothelial | 1.29 | 9.99E-04 | 1.00E+00 | 9.99E-04 | 1 |
| 832 | 71 | recurrent | OPC | MES | 1.57 | 2.00E-03 | 9.99E-01 | 2.00E-03 | 1 |
| 833 | 71 | recurrent | OPC | Macrophage | 1.06 | 8.53E-01 | 1.54E-01 | 1.54E-01 | 0 |
| 834 | 71 | recurrent | OPC | Microglia | 1.22 | 1.11E-01 | 8.90E-01 | 1.11E-01 | 0 |
| 835 | 71 | recurrent | OPC | NK cell | 1.17 | 3.68E-01 | 6.37E-01 | 3.68E-01 | 0 |
| 836 | 71 | recurrent | OPC | NPC | 1.08 | 6.79E-01 | 3.26E-01 | 3.26E-01 | 0 |
| 837 | 71 | recurrent | OPC | Neuron | 1.84 | 9.99E-04 | 1.00E+00 | 9.99E-04 | 1 |
| 838 | 71 | recurrent | OPC | OPC | 2.26 | 9.99E-04 | 1.00E+00 | 9.99E-04 | 1 |
| 839 | 71 | recurrent | OPC | Oligodendrocyte | 1.45 | 9.99E-04 | 1.00E+00 | 9.99E-04 | 1 |
| 840 | 71 | recurrent | OPC | T cell | 1.74 | 9.99E-04 | 1.00E+00 | 9.99E-04 | 1 |
| 841 | 71 | recurrent | Oligodendrocyte | AC | 1.25 | 9.45E-01 | 6.09E-02 | 6.09E-02 | 0 |
| 842 | 71 | recurrent | Oligodendrocyte | Astrocyte | 1.21 | 2.39E-01 | 7.62E-01 | 2.39E-01 | 0 |
| 843 | 71 | recurrent | Oligodendrocyte | Endothelial | 1.68 | 9.99E-04 | 1.00E+00 | 9.99E-04 | 1 |
| 844 | 71 | recurrent | Oligodendrocyte | MES | 1.55 | 1.90E-02 | 9.82E-01 | 1.90E-02 | 0 |
| 845 | 71 | recurrent | Oligodendrocyte | Macrophage | 1.23 | 4.00E-03 | 9.97E-01 | 4.00E-03 | 1 |
| 846 | 71 | recurrent | Oligodendrocyte | Microglia | 1.48 | 9.99E-04 | 1.00E+00 | 9.99E-04 | 1 |
| 847 | 71 | recurrent | Oligodendrocyte | NK cell | 1.15 | 5.27E-01 | 4.78E-01 | 4.78E-01 | 0 |
| 848 | 71 | recurrent | Oligodendrocyte | NPC | 1.00 | 1.00E+00 | 7.99E-03 | 7.99E-03 | -1 |
| 849 | 71 | recurrent | Oligodendrocyte | Neuron | 1.70 | 9.99E-04 | 1.00E+00 | 9.99E-04 | 1 |
| 850 | 71 | recurrent | Oligodendrocyte | OPC | 1.59 | 9.99E-04 | 1.00E+00 | 9.99E-04 | 1 |
| 851 | 71 | recurrent | Oligodendrocyte | Oligodendrocyte | 2.09 | 9.99E-04 | 1.00E+00 | 9.99E-04 | 1 |
| 852 | 71 | recurrent | Oligodendrocyte | T cell | 1.67 | 1.10E-02 | 9.90E-01 | 1.10E-02 | 0 |
| 853 | 71 | recurrent | T cell | AC | 2.12 | 9.99E-04 | 1.00E+00 | 9.99E-04 | 1 |
| 854 | 71 | recurrent | T cell | Astrocyte | 1.37 | 9.99E-04 | 1.00E+00 | 9.99E-04 | 1 |
| 855 | 71 | recurrent | T cell | Endothelial | 1.43 | 9.99E-04 | 1.00E+00 | 9.99E-04 | 1 |
| 856 | 71 | recurrent | T cell | MES | 1.57 | 9.99E-04 | 1.00E+00 | 9.99E-04 | 1 |
| 857 | 71 | recurrent | T cell | Macrophage | 1.13 | 4.00E-02 | 9.61E-01 | 4.00E-02 | 0 |
| 858 | 71 | recurrent | T cell | Microglia | 1.37 | 9.99E-04 | 1.00E+00 | 9.99E-04 | 1 |
| 859 | 71 | recurrent | T cell | NK cell | 1.52 | 9.99E-04 | 1.00E+00 | 9.99E-04 | 1 |
| 860 | 71 | recurrent | T cell | NPC | 1.26 | 9.99E-04 | 1.00E+00 | 9.99E-04 | 1 |
| 861 | 71 | recurrent | T cell | Neuron | 1.47 | 9.99E-04 | 1.00E+00 | 9.99E-04 | 1 |
| 862 | 71 | recurrent | T cell | OPC | 1.57 | 9.99E-04 | 1.00E+00 | 9.99E-04 | 1 |
| 863 | 71 | recurrent | T cell | Oligodendrocyte | 1.49 | 9.99E-04 | 1.00E+00 | 9.99E-04 | 1 |
| 864 | 71 | recurrent | T cell | T cell | 2.74 | 9.99E-04 | 1.00E+00 | 9.99E-04 | 1 |
| 865 | 82 | primary | AC | AC | 3.30 | 9.99E-04 | 1.00E+00 | 9.99E-04 | 1 |
| 866 | 82 | primary | AC | Astrocyte | 1.03 | 1.71E-01 | 8.30E-01 | 1.71E-01 | 0 |
| 867 | 82 | primary | AC | Endothelial | 1.54 | 9.99E-04 | 1.00E+00 | 9.99E-04 | 1 |
| 868 | 82 | primary | AC | MES | 1.21 | 1.00E+00 | 9.99E-04 | 9.99E-04 | -1 |
| 869 | 82 | primary | AC | Macrophage | 1.02 | 1.00E+00 | 9.99E-04 | 9.99E-04 | -1 |
| 870 | 82 | primary | AC | Microglia | 1.14 | 8.41E-01 | 1.60E-01 | 1.60E-01 | 0 |
| 871 | 82 | primary | AC | NK cell | 1.06 | 5.02E-01 | 4.99E-01 | 4.99E-01 | 0 |
| 872 | 82 | primary | AC | NPC | 1.54 | 9.99E-04 | 1.00E+00 | 9.99E-04 | 1 |
| 873 | 82 | primary | AC | Neuron | 1.08 | 1.15E-01 | 8.86E-01 | 1.15E-01 | 0 |
| 874 | 82 | primary | AC | OPC | 1.39 | 9.99E-04 | 1.00E+00 | 9.99E-04 | 1 |
| 875 | 82 | primary | AC | Oligodendrocyte | 1.00 | 1.00E+00 | 8.30E-01 | 8.30E-01 | 0 |

| row | patient | surgery | from cell type | to cell tye | observed count | permutations (greater than) | permutations (less than) | p value | interaction significance/direction |
|---|---|---|---|---|---|---|---|---|---|
| 876 | 82 | primary | AC | T cell | 1.18 | 3.11E-01 | 6.90E-01 | 3.11E-01 | 0 |
| 877 | 82 | primary | Astrocyte | AC | 3.12 | 9.99E-04 | 1.00E+00 | 9.99E-04 | 1 |
| 878 | 82 | primary | Astrocyte | Astrocyte | 1.09 | 3.10E-02 | 9.70E-01 | 3.10E-02 | 0 |
| 879 | 82 | primary | Astrocyte | Endothelial | 1.58 | 5.99E-03 | 9.98E-01 | 5.99E-03 | 1 |
| 880 | 82 | primary | Astrocyte | MES | 2.67 | 9.99E-04 | 1.00E+00 | 9.99E-04 | 1 |
| 881 | 82 | primary | Astrocyte | Macrophage | 1.38 | 4.00E-03 | 9.97E-01 | 4.00E-03 | 1 |
| 882 | 82 | primary | Astrocyte | Microglia | 1.29 | 8.19E-02 | 9.24E-01 | 8.19E-02 | 0 |
| 883 | 82 | primary | Astrocyte | NK cell | 1.00 | 1.00E+00 | 6.39E-01 | 6.39E-01 | 0 |
| 884 | 82 | primary | Astrocyte | NPC | 1.74 | 7.99E-03 | 9.93E-01 | 7.99E-03 | 1 |
| 885 | 82 | primary | Astrocyte | Neuron | 1.22 | 5.49E-02 | 9.56E-01 | 5.49E-02 | 0 |
| 886 | 82 | primary | Astrocyte | OPC | 1.00 | 1.00E+00 | 6.09E-02 | 6.09E-02 | 0 |
| 887 | 82 | primary | Astrocyte | Oligodendrocyte | 1.00 | 6.49E-01 | 9.92E-01 | 6.49E-01 | 0 |
| 888 | 82 | primary | Astrocyte | T cell | 1.23 | 2.61E-01 | 7.48E-01 | 2.61E-01 | 0 |
| 889 | 82 | primary | Endothelial | AC | 2.26 | 9.99E-04 | 1.00E+00 | 9.99E-04 | 1 |
| 890 | 82 | primary | Endothelial | Astrocyte | 1.02 | 4.29E-01 | 5.85E-01 | 4.29E-01 | 0 |
| 891 | 82 | primary | Endothelial | Endothelial | 2.50 | 9.99E-04 | 1.00E+00 | 9.99E-04 | 1 |
| 892 | 82 | primary | Endothelial | MES | 2.24 | 9.99E-04 | 1.00E+00 | 9.99E-04 | 1 |
| 893 | 82 | primary | Endothelial | Macrophage | 1.13 | 4.00E-03 | 9.97E-01 | 4.00E-03 | 1 |
| 894 | 82 | primary | Endothelial | Microglia | 1.20 | 3.50E-02 | 9.66E-01 | 3.50E-02 | 0 |
| 895 | 82 | primary | Endothelial | NK cell | 1.14 | 9.99E-04 | 1.00E+00 | 9.99E-04 | 1 |
| 896 | 82 | primary | Endothelial | NPC | 1.48 | 5.59E-02 | 9.45E-01 | 5.59E-02 | 0 |
| 897 | 82 | primary | Endothelial | Neuron | 1.03 | 9.48E-01 | 5.29E-02 | 5.29E-02 | 0 |
| 898 | 82 | primary | Endothelial | OPC | 1.28 | 9.99E-04 | 1.00E+00 | 9.99E-04 | 1 |
| 899 | 82 | primary | Endothelial | Oligodendrocyte | 1.00 | 1.00E+00 | 9.05E-01 | 9.05E-01 | 0 |
| 900 | 82 | primary | Endothelial | T cell | 1.19 | 2.00E-01 | 8.03E-01 | 2.00E-01 | 0 |
| 901 | 82 | primary | MES | AC | 1.55 | 1.00E+00 | 9.99E-04 | 9.99E-04 | -1 |
| 902 | 82 | primary | MES | Astrocyte | 1.04 | 7.89E-02 | 9.23E-01 | 7.89E-02 | 0 |
| 903 | 82 | primary | MES | Endothelial | 1.36 | 6.99E-03 | 9.94E-01 | 6.99E-03 | 1 |
| 904 | 82 | primary | MES | MES | 4.29 | 9.99E-04 | 1.00E+00 | 9.99E-04 | 1 |
| 905 | 82 | primary | MES | Macrophage | 1.22 | 9.99E-04 | 1.00E+00 | 9.99E-04 | 1 |
| 906 | 82 | primary | MES | Microglia | 1.23 | 9.99E-04 | 1.00E+00 | 9.99E-04 | 1 |
| 907 | 82 | primary | MES | NK cell | 1.10 | 2.30E-02 | 9.78E-01 | 2.30E-02 | 0 |
| 908 | 82 | primary | MES | NPC | 1.24 | 1.00E+00 | 9.99E-04 | 9.99E-04 | -1 |
| 909 | 82 | primary | MES | Neuron | 1.03 | 9.83E-01 | 1.80E-02 | 1.80E-02 | 0 |
| 910 | 82 | primary | MES | OPC | 1.07 | 1.00E+00 | 9.99E-04 | 9.99E-04 | -1 |
| 911 | 82 | primary | MES | Oligodendrocyte | 1.02 | 1.66E-01 | 8.35E-01 | 1.66E-01 | 0 |
| 912 | 82 | primary | MES | T cell | 1.24 | 9.99E-04 | 1.00E+00 | 9.99E-04 | 1 |
| 913 | 82 | primary | Macrophage | AC | 2.40 | 9.99E-04 | 1.00E+00 | 9.99E-04 | 1 |
| 914 | 82 | primary | Macrophage | Astrocyte | 1.00 | 1.00E+00 | 8.32E-01 | 8.32E-01 | 0 |
| 915 | 82 | primary | Macrophage | Endothelial | 1.55 | 9.99E-04 | 1.00E+00 | 9.99E-04 | 1 |
| 916 | 82 | primary | Macrophage | MES | 2.85 | 9.99E-04 | 1.00E+00 | 9.99E-04 | 1 |
| 917 | 82 | primary | Macrophage | Macrophage | 1.17 | 5.19E-02 | 9.50E-01 | 5.19E-02 | 0 |
| 918 | 82 | primary | Macrophage | Microglia | 1.42 | 9.99E-04 | 1.00E+00 | 9.99E-04 | 1 |
| 919 | 82 | primary | Macrophage | NK cell | 1.39 | 9.99E-04 | 1.00E+00 | 9.99E-04 | 1 |
| 920 | 82 | primary | Macrophage | NPC | 1.70 | 9.99E-04 | 1.00E+00 | 9.99E-04 | 1 |
| 921 | 82 | primary | Macrophage | Neuron | 1.21 | 5.99E-03 | 9.95E-01 | 5.99E-03 | 1 |
| 922 | 82 | primary | Macrophage | OPC | 1.22 | 6.59E-02 | 9.36E-01 | 6.59E-02 | 0 |
| 923 | 82 | primary | Macrophage | Oligodendrocyte | 1.00 | 9.77E-01 | 9.82E-01 | 9.77E-01 | 0 |
| 924 | 82 | primary | Macrophage | T cell | 1.28 | 2.00E-02 | 9.81E-01 | 2.00E-02 | 0 |
| 925 | 82 | primary | Microglia | AC | 2.55 | 9.99E-04 | 1.00E+00 | 9.99E-04 | 1 |
| 926 | 82 | primary | Microglia | Astrocyte | 1.05 | 1.72E-01 | 8.53E-01 | 1.72E-01 | 0 |
| 927 | 82 | primary | Microglia | Endothelial | 1.51 | 9.99E-04 | 1.00E+00 | 9.99E-04 | 1 |
| 928 | 82 | primary | Microglia | MES | 2.88 | 9.99E-04 | 1.00E+00 | 9.99E-04 | 1 |
| 929 | 82 | primary | Microglia | Macrophage | 1.30 | 9.99E-04 | 1.00E+00 | 9.99E-04 | 1 |
| 930 | 82 | primary | Microglia | Microglia | 1.30 | 9.99E-04 | 1.00E+00 | 9.99E-04 | 1 |
| 931 | 82 | primary | Microglia | NK cell | 1.28 | 9.99E-04 | 1.00E+00 | 9.99E-04 | 1 |
| 932 | 82 | primary | Microglia | NPC | 1.65 | 9.99E-04 | 1.00E+00 | 9.99E-04 | 1 |
| 933 | 82 | primary | Microglia | Neuron | 1.14 | 1.30E-02 | 9.88E-01 | 1.30E-02 | 0 |
| 934 | 82 | primary | Microglia | OPC | 1.43 | 9.99E-04 | 1.00E+00 | 9.99E-04 | 1 |
| 935 | 82 | primary | Microglia | Oligodendrocyte | 1.00 | 1.00E+00 | 9.52E-01 | 9.52E-01 | 0 |
| 936 | 82 | primary | Microglia | T cell | 1.25 | 1.20E-02 | 9.89E-01 | 1.20E-02 | 0 |
| 937 | 82 | primary | NK cell | AC | 2.44 | 9.99E-04 | 1.00E+00 | 9.99E-04 | 1 |
| 938 | 82 | primary | NK cell | Astrocyte | 1.00 | 1.00E+00 | 8.47E-01 | 8.47E-01 | 0 |
| 939 | 82 | primary | NK cell | Endothelial | 1.65 | 9.99E-04 | 1.00E+00 | 9.99E-04 | 1 |
| 940 | 82 | primary | NK cell | MES | 2.62 | 9.99E-04 | 1.00E+00 | 9.99E-04 | 1 |
| 941 | 82 | primary | NK cell | Macrophage | 1.39 | 9.99E-04 | 1.00E+00 | 9.99E-04 | 1 |
| 942 | 82 | primary | NK cell | Microglia | 1.31 | 3.00E-03 | 9.98E-01 | 3.00E-03 | 1 |
| 943 | 82 | primary | NK cell | NK cell | 1.29 | 1.20E-02 | 9.89E-01 | 1.20E-02 | 0 |
| 944 | 82 | primary | NK cell | NPC | 1.75 | 9.99E-04 | 1.00E+00 | 9.99E-04 | 1 |
| 945 | 82 | primary | NK cell | Neuron | 1.15 | 4.90E-02 | 9.59E-01 | 4.90E-02 | 0 |
| 946 | 82 | primary | NK cell | OPC | 1.30 | 9.99E-03 | 9.91E-01 | 9.99E-03 | 1 |
| 947 | 82 | primary | NK cell | Oligodendrocyte | 1.00 | 9.39E-01 | 9.81E-01 | 9.39E-01 | 0 |
| 948 | 82 | primary | NK cell | T cell | 1.42 | 9.99E-04 | 1.00E+00 | 9.99E-04 | 1 |
| 949 | 82 | primary | NPC | AC | 2.68 | 9.99E-04 | 1.00E+00 | 9.99E-04 | 1 |
| 950 | 82 | primary | NPC | Astrocyte | 1.14 | 9.99E-04 | 1.00E+00 | 9.99E-04 | 1 |
| 951 | 82 | primary | NPC | Endothelial | 1.59 | 9.99E-04 | 1.00E+00 | 9.99E-04 | 1 |
| 952 | 82 | primary | NPC | MES | 3.15 | 9.99E-04 | 1.00E+00 | 9.99E-04 | 1 |
| 953 | 82 | primary | NPC | Macrophage | 1.16 | 9.99E-04 | 1.00E+00 | 9.99E-04 | 1 |
| 954 | 82 | primary | NPC | Microglia | 1.23 | 2.00E-03 | 9.99E-01 | 2.00E-03 | 1 |
| 955 | 82 | primary | NPC | NK cell | 1.08 | 8.19E-02 | 9.20E-01 | 8.19E-02 | 0 |
| 956 | 82 | primary | NPC | NPC | 2.14 | 9.99E-04 | 1.00E+00 | 9.99E-04 | 1 |
| 957 | 82 | primary | NPC | Neuron | 1.23 | 9.99E-04 | 1.00E+00 | 9.99E-04 | 1 |
| 958 | 82 | primary | NPC | OPC | 1.48 | 9.99E-04 | 1.00E+00 | 9.99E-04 | 1 |
| 959 | 82 | primary | NPC | Oligodendrocyte | 1.00 | 1.00E+00 | 8.92E-01 | 8.92E-01 | 0 |
| 960 | 82 | primary | NPC | T cell | 1.21 | 2.80E-02 | 9.73E-01 | 2.80E-02 | 0 |
| 961 | 82 | primary | Neuron | AC | 2.68 | 9.99E-04 | 1.00E+00 | 9.99E-04 | 1 |
| 962 | 82 | primary | Neuron | Astrocyte | 1.00 | 1.00E+00 | 8.65E-01 | 8.65E-01 | 0 |
| 963 | 82 | primary | Neuron | Endothelial | 1.40 | 6.79E-02 | 9.34E-01 | 6.79E-02 | 0 |
| 964 | 82 | primary | Neuron | MES | 2.78 | 9.99E-04 | 1.00E+00 | 9.99E-04 | 1 |
| 965 | 82 | primary | Neuron | Macrophage | 1.21 | 8.99E-03 | 9.92E-01 | 8.99E-03 | 1 |
| 966 | 82 | primary | Neuron | Microglia | 1.30 | 5.99E-03 | 9.95E-01 | 5.99E-03 | 1 |
| 967 | 82 | primary | Neuron | NK cell | 1.11 | 1.33E-01 | 8.76E-01 | 1.33E-01 | 0 |
| 968 | 82 | primary | Neuron | NPC | 1.90 | 9.99E-04 | 1.00E+00 | 9.99E-04 | 1 |
| 969 | 82 | primary | Neuron | Neuron | 1.37 | 3.00E-03 | 9.98E-01 | 3.00E-03 | 1 |
| 970 | 82 | primary | Neuron | OPC | 1.35 | 9.99E-04 | 1.00E+00 | 9.99E-04 | 1 |
| 971 | 82 | primary | Neuron | Oligodendrocyte | 1.00 | 9.47E-01 | 9.80E-01 | 9.47E-01 | 0 |
| 972 | 82 | primary | Neuron | T cell | 1.29 | 2.00E-02 | 9.81E-01 | 2.00E-02 | 0 |
| 973 | 82 | primary | OPC | AC | 2.61 | 9.99E-04 | 1.00E+00 | 9.99E-04 | 1 |
| 974 | 82 | primary | OPC | Astrocyte | 1.00 | 1.00E+00 | 6.84E-01 | 6.84E-01 | 0 |
| 975 | 82 | primary | OPC | Endothelial | 1.46 | 9.99E-04 | 1.00E+00 | 9.99E-04 | 1 |
| 976 | 82 | primary | OPC | MES | 2.67 | 9.99E-04 | 1.00E+00 | 9.99E-04 | 1 |
| 977 | 82 | primary | OPC | Macrophage | 1.22 | 9.99E-04 | 1.00E+00 | 9.99E-04 | 1 |
| 978 | 82 | primary | OPC | Microglia | 1.13 | 8.06E-01 | 2.03E-01 | 2.03E-01 | 0 |
| 979 | 82 | primary | OPC | NK cell | 1.07 | 3.44E-01 | 6.71E-01 | 3.44E-01 | 0 |
| 980 | 82 | primary | OPC | NPC | 1.67 | 9.99E-04 | 1.00E+00 | 9.99E-04 | 1 |
| 981 | 82 | primary | OPC | Neuron | 1.19 | 3.00E-03 | 9.98E-01 | 3.00E-03 | 1 |
| 982 | 82 | primary | OPC | OPC | 1.95 | 9.99E-04 | 1.00E+00 | 9.99E-04 | 1 |
| 983 | 82 | primary | OPC | Oligodendrocyte | 1.00 | 1.00E+00 | 9.55E-01 | 9.55E-01 | 0 |
| 984 | 82 | primary | OPC | T cell | 1.19 | 2.83E-01 | 7.19E-01 | 2.83E-01 | 0 |
| 985 | 82 | primary | Oligodendrocyte | AC | 1.88 | 6.42E-01 | 3.61E-01 | 3.61E-01 | 0 |
| 986 | 82 | primary | Oligodendrocyte | Astrocyte | 1.00 | 6.49E-01 | 9.89E-01 | 6.49E-01 | 0 |

| row | patient | surgery | from cell type | to cell tye | observed count | permutations (greater than) | permutations (less than) | p value | interaction significance/direction |
|---|---|---|---|---|---|---|---|---|---|
| 987 | 82 | primary | Oligodendrocyte | Endothelial | 1.29 | 5.48E-01 | 4.80E-01 | 4.80E-01 | 0 |
| 988 | 82 | primary | Oligodendrocyte | MES | 3.27 | 9.99E-04 | 1.00E+00 | 9.99E-04 | 1 |
| 989 | 82 | primary | Oligodendrocyte | Macrophage | 1.20 | 1.80E-01 | 8.51E-01 | 1.80E-01 | 0 |
| 990 | 82 | primary | Oligodendrocyte | Microglia | 1.11 | 5.83E-01 | 4.64E-01 | 4.64E-01 | 0 |
| 991 | 82 | primary | Oligodendrocyte | NK cell | 1.00 | 9.39E-01 | 8.66E-01 | 8.66E-01 | 0 |
| 992 | 82 | primary | Oligodendrocyte | NPC | 1.55 | 2.71E-01 | 7.40E-01 | 2.71E-01 | 0 |
| 993 | 82 | primary | Oligodendrocyte | Neuron | 1.00 | 9.47E-01 | 8.63E-01 | 8.63E-01 | 0 |
| 994 | 82 | primary | Oligodendrocyte | OPC | 1.40 | 7.59E-02 | 9.39E-01 | 7.59E-02 | 0 |
| 995 | 82 | primary | Oligodendrocyte | Oligodendrocyte | 0.00 | 1.00E+00 | 8.40E-01 | 8.40E-01 | 0 |
| 996 | 82 | primary | Oligodendrocyte | T cell | 1.33 | 1.83E-01 | 8.74E-01 | 1.83E-01 | 0 |
| 997 | 82 | primary | T cell | AC | 2.62 | 9.99E-04 | 1.00E+00 | 9.99E-04 | 1 |
| 998 | 82 | primary | T cell | Astrocyte | 1.04 | 2.29E-01 | 7.92E-01 | 2.29E-01 | 0 |
| 999 | 82 | primary | T cell | Endothelial | 1.59 | 9.99E-04 | 1.00E+00 | 9.99E-04 | 1 |
| 1000 | 82 | primary | T cell | MES | 3.27 | 9.99E-04 | 1.00E+00 | 9.99E-04 | 1 |
| 1001 | 82 | primary | T cell | Macrophage | 1.14 | 1.20E-02 | 9.89E-01 | 1.20E-02 | 0 |
| 1002 | 82 | primary | T cell | Microglia | 1.40 | 9.99E-04 | 1.00E+00 | 9.99E-04 | 1 |
| 1003 | 82 | primary | T cell | NK cell | 1.25 | 9.99E-04 | 1.00E+00 | 9.99E-04 | 1 |
| 1004 | 82 | primary | T cell | NPC | 1.64 | 9.99E-04 | 1.00E+00 | 9.99E-04 | 1 |
| 1005 | 82 | primary | T cell | Neuron | 1.08 | 2.91E-01 | 7.19E-01 | 2.91E-01 | 0 |
| 1006 | 82 | primary | T cell | OPC | 1.56 | 9.99E-04 | 1.00E+00 | 9.99E-04 | 1 |
| 1007 | 82 | primary | T cell | Oligodendrocyte | 1.00 | 9.99E-01 | 9.61E-01 | 9.61E-01 | 0 |
| 1008 | 82 | primary | T cell | T cell | 1.43 | 9.99E-04 | 1.00E+00 | 9.99E-04 | 1 |
| 1009 | 82 | recurrent | AC | AC | 1.00 | 2.48E-01 | 9.93E-01 | 2.48E-01 | 0 |
| 1010 | 82 | recurrent | AC | Astrocyte | 2.31 | 4.00E-03 | 9.97E-01 | 4.00E-03 | 1 |
| 1011 | 82 | recurrent | AC | Endothelial | 1.25 | 1.92E-01 | 8.57E-01 | 1.92E-01 | 0 |
| 1012 | 82 | recurrent | AC | MES | 1.00 | 3.90E-01 | 9.97E-01 | 3.90E-01 | 0 |
| 1013 | 82 | recurrent | AC | Macrophage | 1.00 | 9.56E-01 | 8.94E-01 | 8.94E-01 | 0 |
| 1014 | 82 | recurrent | AC | Microglia | 1.00 | 9.90E-01 | 7.73E-01 | 7.73E-01 | 0 |
| 1015 | 82 | recurrent | AC | NK cell | 1.00 | 1.00E+00 | 6.77E-01 | 6.77E-01 | 0 |
| 1016 | 82 | recurrent | AC | NPC | 1.00 | 5.51E-01 | 9.93E-01 | 5.51E-01 | 0 |
| 1017 | 82 | recurrent | AC | Neuron | 2.67 | 5.98E-01 | 4.18E-01 | 4.18E-01 | 0 |
| 1018 | 82 | recurrent | AC | OPC | 0.00 | 1.00E+00 | 5.09E-02 | 5.09E-02 | 0 |
| 1019 | 82 | recurrent | AC | Oligodendrocyte | 1.36 | 1.28E-01 | 8.80E-01 | 1.28E-01 | 0 |
| 1020 | 82 | recurrent | AC | T cell | 1.20 | 8.06E-01 | 2.11E-01 | 2.11E-01 | 0 |
| 1021 | 82 | recurrent | Astrocyte | AC | 1.03 | 5.00E-02 | 9.51E-01 | 5.00E-02 | 0 |
| 1022 | 82 | recurrent | Astrocyte | Astrocyte | 2.48 | 9.99E-04 | 1.00E+00 | 9.99E-04 | 1 |
| 1023 | 82 | recurrent | Astrocyte | Endothelial | 1.26 | 9.99E-04 | 1.00E+00 | 9.99E-04 | 1 |
| 1024 | 82 | recurrent | Astrocyte | MES | 1.02 | 1.61E-01 | 8.41E-01 | 1.61E-01 | 0 |
| 1025 | 82 | recurrent | Astrocyte | Macrophage | 1.08 | 8.99E-03 | 9.92E-01 | 8.99E-03 | 1 |
| 1026 | 82 | recurrent | Astrocyte | Microglia | 1.09 | 6.69E-02 | 9.34E-01 | 6.69E-02 | 0 |
| 1027 | 82 | recurrent | Astrocyte | NK cell | 1.12 | 5.99E-03 | 9.95E-01 | 5.99E-03 | 1 |
| 1028 | 82 | recurrent | Astrocyte | NPC | 1.04 | 5.59E-02 | 9.46E-01 | 5.59E-02 | 0 |
| 1029 | 82 | recurrent | Astrocyte | Neuron | 2.63 | 1.00E+00 | 9.99E-04 | 9.99E-04 | -1 |
| 1030 | 82 | recurrent | Astrocyte | OPC | 1.08 | 9.99E-04 | 1.00E+00 | 9.99E-04 | 1 |
| 1031 | 82 | recurrent | Astrocyte | Oligodendrocyte | 1.28 | 9.99E-04 | 1.00E+00 | 9.99E-04 | 1 |
| 1032 | 82 | recurrent | Astrocyte | T cell | 1.37 | 2.00E-03 | 9.99E-01 | 2.00E-03 | 1 |
| 1033 | 82 | recurrent | Endothelial | AC | 1.00 | 1.00E+00 | 9.24E-01 | 9.24E-01 | 0 |
| 1034 | 82 | recurrent | Endothelial | Astrocyte | 2.01 | 9.99E-04 | 1.00E+00 | 9.99E-04 | 1 |
| 1035 | 82 | recurrent | Endothelial | Endothelial | 1.96 | 9.99E-04 | 1.00E+00 | 9.99E-04 | 1 |
| 1036 | 82 | recurrent | Endothelial | MES | 1.00 | 1.00E+00 | 9.59E-01 | 9.59E-01 | 0 |
| 1037 | 82 | recurrent | Endothelial | Macrophage | 1.17 | 9.99E-04 | 1.00E+00 | 9.99E-04 | 1 |
| 1038 | 82 | recurrent | Endothelial | Microglia | 1.11 | 7.09E-02 | 9.32E-01 | 7.09E-02 | 0 |
| 1039 | 82 | recurrent | Endothelial | NK cell | 1.22 | 9.99E-04 | 1.00E+00 | 9.99E-04 | 1 |
| 1040 | 82 | recurrent | Endothelial | NPC | 1.00 | 1.00E+00 | 8.75E-01 | 8.75E-01 | 0 |
| 1041 | 82 | recurrent | Endothelial | Neuron | 2.37 | 1.00E+00 | 9.99E-04 | 9.99E-04 | -1 |
| 1042 | 82 | recurrent | Endothelial | OPC | 1.05 | 2.71E-01 | 7.43E-01 | 2.71E-01 | 0 |
| 1043 | 82 | recurrent | Endothelial | Oligodendrocyte | 1.31 | 9.99E-04 | 1.00E+00 | 9.99E-04 | 1 |
| 1044 | 82 | recurrent | Endothelial | T cell | 1.52 | 9.99E-04 | 1.00E+00 | 9.99E-04 | 1 |
| 1045 | 82 | recurrent | MES | AC | 1.00 | 3.90E-01 | 9.97E-01 | 3.90E-01 | 0 |
| 1046 | 82 | recurrent | MES | Astrocyte | 2.43 | 2.00E-03 | 9.99E-01 | 2.00E-03 | 1 |
| 1047 | 82 | recurrent | MES | Endothelial | 1.00 | 1.00E+00 | 4.05E-01 | 4.05E-01 | 0 |
| 1048 | 82 | recurrent | MES | MES | 1.00 | 1.51E-01 | 1.00E+00 | 1.51E-01 | 0 |
| 1049 | 82 | recurrent | MES | Macrophage | 1.00 | 8.94E-01 | 9.31E-01 | 8.94E-01 | 0 |
| 1050 | 82 | recurrent | MES | Microglia | 1.00 | 9.75E-01 | 7.93E-01 | 7.93E-01 | 0 |
| 1051 | 82 | recurrent | MES | NK cell | 1.00 | 9.96E-01 | 7.18E-01 | 7.18E-01 | 0 |
| 1052 | 82 | recurrent | MES | NPC | 0.00 | 1.00E+00 | 5.13E-01 | 5.13E-01 | 0 |
| 1053 | 82 | recurrent | MES | Neuron | 2.10 | 9.96E-01 | 5.00E-03 | 5.00E-03 | -1 |
| 1054 | 82 | recurrent | MES | OPC | 0.00 | 1.00E+00 | 1.13E-01 | 1.13E-01 | 0 |
| 1055 | 82 | recurrent | MES | Oligodendrocyte | 1.20 | 5.11E-01 | 5.55E-01 | 5.11E-01 | 0 |
| 1056 | 82 | recurrent | MES | T cell | 1.54 | 1.01E-01 | 9.08E-01 | 1.01E-01 | 0 |
| 1057 | 82 | recurrent | Macrophage | AC | 1.33 | 5.99E-03 | 9.99E-01 | 5.99E-03 | 1 |
| 1058 | 82 | recurrent | Macrophage | Astrocyte | 1.71 | 7.74E-01 | 2.27E-01 | 2.27E-01 | 0 |
| 1059 | 82 | recurrent | Macrophage | Endothelial | 1.36 | 9.99E-04 | 1.00E+00 | 9.99E-04 | 1 |
| 1060 | 82 | recurrent | Macrophage | MES | 1.00 | 8.94E-01 | 9.90E-01 | 8.94E-01 | 0 |
| 1061 | 82 | recurrent | Macrophage | Macrophage | 1.14 | 8.69E-02 | 9.30E-01 | 8.69E-02 | 0 |
| 1062 | 82 | recurrent | Macrophage | Microglia | 1.35 | 2.00E-03 | 9.99E-01 | 2.00E-03 | 1 |
| 1063 | 82 | recurrent | Macrophage | NK cell | 1.21 | 2.00E-02 | 9.81E-01 | 2.00E-02 | 0 |
| 1064 | 82 | recurrent | Macrophage | NPC | 1.25 | 1.70E-02 | 9.90E-01 | 1.70E-02 | 0 |
| 1065 | 82 | recurrent | Macrophage | Neuron | 2.81 | 2.05E-01 | 7.96E-01 | 2.05E-01 | 0 |
| 1066 | 82 | recurrent | Macrophage | OPC | 1.00 | 1.00E+00 | 6.54E-01 | 6.54E-01 | 0 |
| 1067 | 82 | recurrent | Macrophage | Oligodendrocyte | 1.48 | 9.99E-04 | 1.00E+00 | 9.99E-04 | 1 |
| 1068 | 82 | recurrent | Macrophage | T cell | 1.80 | 9.99E-04 | 1.00E+00 | 9.99E-04 | 1 |
| 1069 | 82 | recurrent | Microglia | AC | 1.00 | 9.90E-01 | 9.65E-01 | 9.65E-01 | 0 |
| 1070 | 82 | recurrent | Microglia | Astrocyte | 1.82 | 1.31E-01 | 8.70E-01 | 1.31E-01 | 0 |
| 1071 | 82 | recurrent | Microglia | Endothelial | 1.27 | 7.99E-03 | 9.93E-01 | 7.99E-03 | 1 |
| 1072 | 82 | recurrent | Microglia | MES | 1.33 | 5.99E-03 | 9.99E-01 | 5.99E-03 | 1 |
| 1073 | 82 | recurrent | Microglia | Macrophage | 1.11 | 6.89E-02 | 9.38E-01 | 6.89E-02 | 0 |
| 1074 | 82 | recurrent | Microglia | Microglia | 1.39 | 9.99E-04 | 1.00E+00 | 9.99E-04 | 1 |
| 1075 | 82 | recurrent | Microglia | NK cell | 1.44 | 9.99E-04 | 1.00E+00 | 9.99E-04 | 1 |
| 1076 | 82 | recurrent | Microglia | NPC | 1.00 | 9.99E-01 | 9.43E-01 | 9.43E-01 | 0 |
| 1077 | 82 | recurrent | Microglia | Neuron | 2.63 | 9.27E-01 | 7.39E-02 | 7.39E-02 | 0 |
| 1078 | 82 | recurrent | Microglia | OPC | 1.11 | 6.39E-02 | 9.47E-01 | 6.39E-02 | 0 |
| 1079 | 82 | recurrent | Microglia | Oligodendrocyte | 1.37 | 9.99E-04 | 1.00E+00 | 9.99E-04 | 1 |
| 1080 | 82 | recurrent | Microglia | T cell | 1.65 | 9.99E-04 | 1.00E+00 | 9.99E-04 | 1 |
| 1081 | 82 | recurrent | NK cell | AC | 1.00 | 1.00E+00 | 9.58E-01 | 9.58E-01 | 0 |
| 1082 | 82 | recurrent | NK cell | Astrocyte | 1.63 | 9.95E-01 | 5.99E-03 | 5.99E-03 | -1 |
| 1083 | 82 | recurrent | NK cell | Endothelial | 1.40 | 9.99E-04 | 1.00E+00 | 9.99E-04 | 1 |
| 1084 | 82 | recurrent | NK cell | MES | 1.00 | 9.96E-01 | 9.82E-01 | 9.82E-01 | 0 |
| 1085 | 82 | recurrent | NK cell | Macrophage | 1.09 | 8.69E-02 | 9.14E-01 | 8.69E-02 | 0 |
| 1086 | 82 | recurrent | NK cell | Microglia | 1.32 | 9.99E-04 | 1.00E+00 | 9.99E-04 | 1 |
| 1087 | 82 | recurrent | NK cell | NK cell | 1.46 | 9.99E-04 | 1.00E+00 | 9.99E-04 | 1 |
| 1088 | 82 | recurrent | NK cell | NPC | 1.00 | 1.00E+00 | 9.16E-01 | 9.16E-01 | 0 |
| 1089 | 82 | recurrent | NK cell | Neuron | 2.26 | 1.00E+00 | 9.99E-04 | 9.99E-04 | -1 |
| 1090 | 82 | recurrent | NK cell | OPC | 1.33 | 9.99E-04 | 1.00E+00 | 9.99E-04 | 1 |
| 1091 | 82 | recurrent | NK cell | Oligodendrocyte | 1.40 | 9.99E-04 | 1.00E+00 | 9.99E-04 | 1 |
| 1092 | 82 | recurrent | NK cell | T cell | 1.91 | 9.99E-04 | 1.00E+00 | 9.99E-04 | 1 |
| 1093 | 82 | recurrent | NPC | AC | 1.00 | 5.51E-01 | 9.94E-01 | 5.51E-01 | 0 |
| 1094 | 82 | recurrent | NPC | Astrocyte | 1.81 | 3.64E-01 | 6.38E-01 | 3.64E-01 | 0 |
| 1095 | 82 | recurrent | NPC | Endothelial | 1.30 | 8.99E-02 | 9.18E-01 | 8.99E-02 | 0 |
| 1096 | 82 | recurrent | NPC | MES | 0.00 | 1.00E+00 | 5.13E-01 | 5.13E-01 | 0 |
| 1097 | 82 | recurrent | NPC | Macrophage | 1.00 | 9.81E-01 | 8.56E-01 | 8.56E-01 | 0 |

| row | patient | surgery | from cell type | to cell tye | observed count | permutations (greater than) | permutations (less than) | p value | interaction significance/direction |
|---|---|---|---|---|---|---|---|---|---|
| 1098 | 82 | recurrent | NPC | Microglia | 1.00 | 9.99E-01 | 6.86E-01 | 6.86E-01 | 0 |
| 1099 | 82 | recurrent | NPC | NK cell | 1.20 | 1.52E-01 | 8.94E-01 | 1.52E-01 | 0 |
| 1100 | 82 | recurrent | NPC | NPC | 2.00 | 3.00E-03 | 1.00E+00 | 3.00E-03 | 1 |
| 1101 | 82 | recurrent | NPC | Neuron | 2.74 | 4.86E-01 | 5.26E-01 | 4.86E-01 | 0 |
| 1102 | 82 | recurrent | NPC | OPC | 1.33 | 3.60E-02 | 9.81E-01 | 3.60E-02 | 0 |
| 1103 | 82 | recurrent | NPC | Oligodendrocyte | 1.33 | 1.50E-01 | 8.86E-01 | 1.50E-01 | 0 |
| 1104 | 82 | recurrent | NPC | T cell | 1.75 | 2.00E-03 | 9.99E-01 | 2.00E-03 | 1 |
| 1105 | 82 | recurrent | Neuron | AC | 1.00 | 1.00E+00 | 5.69E-01 | 5.69E-01 | 0 |
| 1106 | 82 | recurrent | Neuron | Astrocyte | 1.99 | 9.99E-04 | 1.00E+00 | 9.99E-04 | 1 |
| 1107 | 82 | recurrent | Neuron | Endothelial | 1.25 | 9.99E-04 | 1.00E+00 | 9.99E-04 | 1 |
| 1108 | 82 | recurrent | Neuron | MES | 1.07 | 2.00E-03 | 9.99E-01 | 2.00E-03 | 1 |
| 1109 | 82 | recurrent | Neuron | Macrophage | 1.04 | 4.66E-01 | 5.37E-01 | 4.66E-01 | 0 |
| 1110 | 82 | recurrent | Neuron | Microglia | 1.07 | 2.23E-01 | 7.80E-01 | 2.23E-01 | 0 |
| 1111 | 82 | recurrent | Neuron | NK cell | 1.23 | 9.99E-04 | 1.00E+00 | 9.99E-04 | 1 |
| 1112 | 82 | recurrent | Neuron | NPC | 1.02 | 2.36E-01 | 7.76E-01 | 2.36E-01 | 0 |
| 1113 | 82 | recurrent | Neuron | Neuron | 3.17 | 9.99E-04 | 1.00E+00 | 9.99E-04 | 1 |
| 1114 | 82 | recurrent | Neuron | OPC | 1.12 | 9.99E-04 | 1.00E+00 | 9.99E-04 | 1 |
| 1115 | 82 | recurrent | Neuron | Oligodendrocyte | 1.37 | 9.99E-04 | 1.00E+00 | 9.99E-04 | 1 |
| 1116 | 82 | recurrent | Neuron | T cell | 1.52 | 9.99E-04 | 1.00E+00 | 9.99E-04 | 1 |
| 1117 | 82 | recurrent | OPC | AC | 0.00 | 1.00E+00 | 5.09E-02 | 5.09E-02 | 0 |
| 1118 | 82 | recurrent | OPC | Astrocyte | 2.31 | 9.99E-04 | 1.00E+00 | 9.99E-04 | 1 |
| 1119 | 82 | recurrent | OPC | Endothelial | 1.15 | 3.97E-01 | 6.23E-01 | 3.97E-01 | 0 |
| 1120 | 82 | recurrent | OPC | MES | 0.00 | 1.00E+00 | 1.13E-01 | 1.13E-01 | 0 |
| 1121 | 82 | recurrent | OPC | Macrophage | 1.00 | 1.00E+00 | 6.12E-01 | 6.12E-01 | 0 |
| 1122 | 82 | recurrent | OPC | Microglia | 1.11 | 2.02E-01 | 8.16E-01 | 2.02E-01 | 0 |
| 1123 | 82 | recurrent | OPC | NK cell | 1.33 | 4.00E-03 | 9.98E-01 | 4.00E-03 | 1 |
| 1124 | 82 | recurrent | OPC | NPC | 1.00 | 9.79E-01 | 9.72E-01 | 9.72E-01 | 0 |
| 1125 | 82 | recurrent | OPC | Neuron | 3.11 | 9.99E-04 | 1.00E+00 | 9.99E-04 | 1 |
| 1126 | 82 | recurrent | OPC | OPC | 1.12 | 8.49E-02 | 9.16E-01 | 8.49E-02 | 0 |
| 1127 | 82 | recurrent | OPC | Oligodendrocyte | 1.37 | 1.40E-02 | 9.87E-01 | 1.40E-02 | 0 |
| 1128 | 82 | recurrent | OPC | T cell | 1.21 | 9.52E-01 | 4.90E-02 | 4.90E-02 | 0 |
| 1129 | 82 | recurrent | Oligodendrocyte | AC | 1.00 | 1.00E+00 | 8.92E-01 | 8.92E-01 | 0 |
| 1130 | 82 | recurrent | Oligodendrocyte | Astrocyte | 1.86 | 2.00E-03 | 9.99E-01 | 2.00E-03 | 1 |
| 1131 | 82 | recurrent | Oligodendrocyte | Endothelial | 1.38 | 9.99E-04 | 1.00E+00 | 9.99E-04 | 1 |
| 1132 | 82 | recurrent | Oligodendrocyte | MES | 1.09 | 2.80E-02 | 9.79E-01 | 2.80E-02 | 0 |
| 1133 | 82 | recurrent | Oligodendrocyte | Macrophage | 1.01 | 8.81E-01 | 1.29E-01 | 1.29E-01 | 0 |
| 1134 | 82 | recurrent | Oligodendrocyte | Microglia | 1.18 | 9.99E-04 | 1.00E+00 | 9.99E-04 | 1 |
| 1135 | 82 | recurrent | Oligodendrocyte | NK cell | 1.29 | 9.99E-04 | 1.00E+00 | 9.99E-04 | 1 |
| 1136 | 82 | recurrent | Oligodendrocyte | NPC | 1.04 | 1.44E-01 | 8.72E-01 | 1.44E-01 | 0 |
| 1137 | 82 | recurrent | Oligodendrocyte | Neuron | 2.79 | 1.01E-01 | 9.00E-01 | 1.01E-01 | 0 |
| 1138 | 82 | recurrent | Oligodendrocyte | OPC | 1.10 | 5.00E-03 | 9.96E-01 | 5.00E-03 | 1 |
| 1139 | 82 | recurrent | Oligodendrocyte | Oligodendrocyte | 1.68 | 9.99E-04 | 1.00E+00 | 9.99E-04 | 1 |
| 1140 | 82 | recurrent | Oligodendrocyte | T cell | 1.48 | 9.99E-04 | 1.00E+00 | 9.99E-04 | 1 |
| 1141 | 82 | recurrent | T cell | AC | 1.00 | 1.00E+00 | 8.61E-01 | 8.61E-01 | 0 |
| 1142 | 82 | recurrent | T cell | Astrocyte | 1.77 | 4.01E-01 | 6.00E-01 | 4.01E-01 | 0 |
| 1143 | 82 | recurrent | T cell | Endothelial | 1.31 | 9.99E-04 | 1.00E+00 | 9.99E-04 | 1 |
| 1144 | 82 | recurrent | T cell | MES | 1.00 | 1.00E+00 | 9.24E-01 | 9.24E-01 | 0 |
| 1145 | 82 | recurrent | T cell | Macrophage | 1.07 | 7.19E-02 | 9.32E-01 | 7.19E-02 | 0 |
| 1146 | 82 | recurrent | T cell | Microglia | 1.19 | 9.99E-04 | 1.00E+00 | 9.99E-04 | 1 |
| 1147 | 82 | recurrent | T cell | NK cell | 1.30 | 9.99E-04 | 1.00E+00 | 9.99E-04 | 1 |
| 1148 | 82 | recurrent | T cell | NPC | 1.00 | 1.00E+00 | 7.44E-01 | 7.44E-01 | 0 |
| 1149 | 82 | recurrent | T cell | Neuron | 2.58 | 1.00E+00 | 9.99E-04 | 9.99E-04 | -1 |
| 1150 | 82 | recurrent | T cell | OPC | 1.08 | 1.10E-02 | 9.91E-01 | 1.10E-02 | 0 |
| 1151 | 82 | recurrent | T cell | Oligodendrocyte | 1.32 | 9.99E-04 | 1.00E+00 | 9.99E-04 | 1 |
| 1152 | 82 | recurrent | T cell | T cell | 1.95 | 9.99E-04 | 1.00E+00 | 9.99E-04 | 1 |
| 1153 | 84 | primary | AC | AC | 0.00 | 1.00E+00 | 9.98E-01 | 9.98E-01 | 0 |
| 1154 | 84 | primary | AC | Astrocyte | 3.67 | 8.19E-02 | 9.61E-01 | 8.19E-02 | 0 |
| 1155 | 84 | primary | AC | Endothelial | 1.00 | 7.88E-01 | 7.68E-01 | 7.68E-01 | 0 |
| 1156 | 84 | primary | AC | MES | 1.50 | 9.23E-01 | 1.20E-01 | 1.20E-01 | 0 |
| 1157 | 84 | primary | AC | Macrophage | 3.00 | 2.00E-03 | 1.00E+00 | 2.00E-03 | 1 |
| 1158 | 84 | primary | AC | Microglia | 0.00 | 1.00E+00 | 6.12E-01 | 6.12E-01 | 0 |
| 1159 | 84 | primary | AC | NK cell | 1.00 | 5.07E-01 | 9.45E-01 | 5.07E-01 | 0 |
| 1160 | 84 | primary | AC | NPC | 0.00 | 1.00E+00 | 9.77E-01 | 9.77E-01 | 0 |
| 1161 | 84 | primary | AC | Neuron | 0.00 | 1.00E+00 | 8.60E-01 | 8.60E-01 | 0 |
| 1162 | 84 | primary | AC | OPC | 0.00 | 1.00E+00 | 9.69E-01 | 9.69E-01 | 0 |
| 1163 | 84 | primary | AC | Oligodendrocyte | | | | | |
| 1164 | 84 | primary | AC | T cell | 0.00 | 1.00E+00 | 6.97E-01 | 6.97E-01 | 0 |
| 1165 | 84 | primary | Astrocyte | AC | 1.00 | 1.00E+00 | 9.97E-01 | 9.97E-01 | 0 |
| 1166 | 84 | primary | Astrocyte | Astrocyte | 3.81 | 9.99E-04 | 1.00E+00 | 9.99E-04 | 1 |
| 1167 | 84 | primary | Astrocyte | Endothelial | 1.42 | 9.99E-04 | 1.00E+00 | 9.99E-04 | 1 |
| 1168 | 84 | primary | Astrocyte | MES | 2.32 | 9.96E-01 | 5.00E-03 | 5.00E-03 | -1 |
| 1169 | 84 | primary | Astrocyte | Macrophage | 1.19 | 9.99E-04 | 1.00E+00 | 9.99E-04 | 1 |
| 1170 | 84 | primary | Astrocyte | Microglia | 1.11 | 2.00E-03 | 9.99E-01 | 2.00E-03 | 1 |
| 1171 | 84 | primary | Astrocyte | NK cell | 1.35 | 9.99E-04 | 1.00E+00 | 9.99E-04 | 1 |
| 1172 | 84 | primary | Astrocyte | NPC | 1.00 | 1.00E+00 | 9.19E-01 | 9.19E-01 | 0 |
| 1173 | 84 | primary | Astrocyte | Neuron | 1.10 | 9.99E-04 | 1.00E+00 | 9.99E-04 | 1 |
| 1174 | 84 | primary | Astrocyte | OPC | 1.07 | 3.00E-03 | 9.98E-01 | 3.00E-03 | 1 |
| 1175 | 84 | primary | Astrocyte | Oligodendrocyte | | | | | |
| 1176 | 84 | primary | Astrocyte | T cell | 1.11 | 9.99E-04 | 1.00E+00 | 9.99E-04 | 1 |
| 1177 | 84 | primary | Endothelial | AC | 1.00 | 7.88E-01 | 1.00E+00 | 7.88E-01 | 0 |
| 1178 | 84 | primary | Endothelial | Astrocyte | 2.49 | 9.39E-01 | 6.19E-02 | 6.19E-02 | 0 |
| 1179 | 84 | primary | Endothelial | Endothelial | 2.24 | 9.99E-04 | 1.00E+00 | 9.99E-04 | 1 |
| 1180 | 84 | primary | Endothelial | MES | 2.59 | 9.99E-04 | 1.00E+00 | 9.99E-04 | 1 |
| 1181 | 84 | primary | Endothelial | Macrophage | 1.13 | 1.53E-01 | 8.48E-01 | 1.53E-01 | 0 |
| 1182 | 84 | primary | Endothelial | Microglia | 1.16 | 2.00E-03 | 9.99E-01 | 2.00E-03 | 1 |
| 1183 | 84 | primary | Endothelial | NK cell | 1.49 | 9.99E-04 | 1.00E+00 | 9.99E-04 | 1 |
| 1184 | 84 | primary | Endothelial | NPC | 1.00 | 9.99E-01 | 9.76E-01 | 9.76E-01 | 0 |
| 1185 | 84 | primary | Endothelial | Neuron | 1.10 | 1.60E-02 | 9.85E-01 | 1.60E-02 | 0 |
| 1186 | 84 | primary | Endothelial | OPC | 1.00 | 1.00E+00 | 9.54E-01 | 9.54E-01 | 0 |
| 1187 | 84 | primary | Endothelial | Oligodendrocyte | | | | | |
| 1188 | 84 | primary | Endothelial | T cell | 1.16 | 2.00E-03 | 9.99E-01 | 2.00E-03 | 1 |
| 1189 | 84 | primary | MES | AC | 1.00 | 9.99E-01 | 9.95E-01 | 9.95E-01 | 0 |
| 1190 | 84 | primary | MES | Astrocyte | 2.18 | 1.00E+00 | 9.99E-04 | 9.99E-04 | -1 |
| 1191 | 84 | primary | MES | Endothelial | 1.35 | 9.99E-04 | 1.00E+00 | 9.99E-04 | 1 |
| 1192 | 84 | primary | MES | MES | 3.97 | 9.99E-04 | 1.00E+00 | 9.99E-04 | 1 |
| 1193 | 84 | primary | MES | Macrophage | 1.15 | 9.99E-04 | 1.00E+00 | 9.99E-04 | 1 |
| 1194 | 84 | primary | MES | Microglia | 1.12 | 9.99E-04 | 1.00E+00 | 9.99E-04 | 1 |
| 1195 | 84 | primary | MES | NK cell | 1.24 | 9.99E-04 | 1.00E+00 | 9.99E-04 | 1 |
| 1196 | 84 | primary | MES | NPC | 1.03 | 6.49E-02 | 9.44E-01 | 6.49E-02 | 0 |
| 1197 | 84 | primary | MES | Neuron | 1.06 | 5.99E-03 | 9.95E-01 | 5.99E-03 | 1 |
| 1198 | 84 | primary | MES | OPC | 1.03 | 5.09E-02 | 9.54E-01 | 5.09E-02 | 0 |
| 1199 | 84 | primary | MES | Oligodendrocyte | | | | | |
| 1200 | 84 | primary | MES | T cell | 1.14 | 9.99E-04 | 1.00E+00 | 9.99E-04 | 1 |
| 1201 | 84 | primary | Macrophage | AC | 1.00 | 5.10E-01 | 9.99E-01 | 5.10E-01 | 0 |
| 1202 | 84 | primary | Macrophage | Astrocyte | 3.44 | 9.99E-04 | 1.00E+00 | 9.99E-04 | 1 |
| 1203 | 84 | primary | Macrophage | Endothelial | 1.54 | 9.99E-04 | 1.00E+00 | 9.99E-04 | 1 |
| 1204 | 84 | primary | Macrophage | MES | 2.19 | 9.98E-01 | 3.00E-03 | 3.00E-03 | -1 |
| 1205 | 84 | primary | Macrophage | Macrophage | 1.43 | 9.99E-04 | 1.00E+00 | 9.99E-04 | 1 |
| 1206 | 84 | primary | Macrophage | Microglia | 1.27 | 9.99E-04 | 1.00E+00 | 9.99E-04 | 1 |
| 1207 | 84 | primary | Macrophage | NK cell | 1.66 | 9.99E-04 | 1.00E+00 | 9.99E-04 | 1 |
| 1208 | 84 | primary | Macrophage | NPC | 1.00 | 9.65E-01 | 9.91E-01 | 9.65E-01 | 0 |

| row | patient | surgery | from cell type | to cell tye | observed count | permutations (greater than) | permutations (less than) | p value | interaction significance/direction |
|---|---|---|---|---|---|---|---|---|---|
| 1209 | 84 | primary | Macrophage | Neuron | 1.17 | 3.00E-03 | 9.98E-01 | 3.00E-03 | 1 |
| 1210 | 84 | primary | Macrophage | OPC | 1.00 | 9.91E-01 | 9.81E-01 | 9.81E-01 | 0 |
| 1211 | 84 | primary | Macrophage | Oligodendrocyte | | | | | |
| 1212 | 84 | primary | Macrophage | T cell | 1.13 | 2.90E-02 | 9.73E-01 | 2.90E-02 | 0 |
| 1213 | 84 | primary | Microglia | AC | 0.00 | 1.00E+00 | 6.12E-01 | 6.12E-01 | 0 |
| 1214 | 84 | primary | Microglia | Astrocyte | 3.30 | 9.99E-04 | 1.00E+00 | 9.99E-04 | 1 |
| 1215 | 84 | primary | Microglia | Endothelial | 1.58 | 9.99E-04 | 1.00E+00 | 9.99E-04 | 1 |
| 1216 | 84 | primary | Microglia | MES | 2.48 | 2.80E-02 | 9.73E-01 | 2.80E-02 | 0 |
| 1217 | 84 | primary | Microglia | Macrophage | 1.34 | 9.99E-04 | 1.00E+00 | 9.99E-04 | 1 |
| 1218 | 84 | primary | Microglia | Microglia | 1.20 | 2.50E-02 | 9.76E-01 | 2.50E-02 | 0 |
| 1219 | 84 | primary | Microglia | NK cell | 1.59 | 9.99E-04 | 1.00E+00 | 9.99E-04 | 1 |
| 1220 | 84 | primary | Microglia | NPC | 1.00 | 8.83E-01 | 9.96E-01 | 8.83E-01 | 0 |
| 1221 | 84 | primary | Microglia | Neuron | 1.07 | 1.60E-01 | 8.66E-01 | 1.60E-01 | 0 |
| 1222 | 84 | primary | Microglia | OPC | 0.00 | 1.00E+00 | 3.40E-02 | 3.40E-02 | 0 |
| 1223 | 84 | primary | Microglia | Oligodendrocyte | | | | | |
| 1224 | 84 | primary | Microglia | T cell | 1.29 | 9.99E-04 | 1.00E+00 | 9.99E-04 | 1 |
| 1225 | 84 | primary | NK cell | AC | 1.00 | 5.07E-01 | 9.99E-01 | 5.07E-01 | 0 |
| 1226 | 84 | primary | NK cell | Astrocyte | 2.38 | 9.97E-01 | 4.00E-03 | 4.00E-03 | -1 |
| 1227 | 84 | primary | NK cell | Endothelial | 1.63 | 9.99E-04 | 1.00E+00 | 9.99E-04 | 1 |
| 1228 | 84 | primary | NK cell | MES | 2.46 | 3.70E-02 | 9.64E-01 | 3.70E-02 | 0 |
| 1229 | 84 | primary | NK cell | Macrophage | 1.56 | 9.99E-04 | 1.00E+00 | 9.99E-04 | 1 |
| 1230 | 84 | primary | NK cell | Microglia | 1.25 | 9.99E-04 | 1.00E+00 | 9.99E-04 | 1 |
| 1231 | 84 | primary | NK cell | NK cell | 1.93 | 9.99E-04 | 1.00E+00 | 9.99E-04 | 1 |
| 1232 | 84 | primary | NK cell | NPC | 1.00 | 9.69E-01 | 9.87E-01 | 9.69E-01 | 0 |
| 1233 | 84 | primary | NK cell | Neuron | 1.00 | 1.00E+00 | 7.00E-01 | 7.00E-01 | 0 |
| 1234 | 84 | primary | NK cell | OPC | 1.00 | 9.85E-01 | 9.76E-01 | 9.76E-01 | 0 |
| 1235 | 84 | primary | NK cell | Oligodendrocyte | | | | | |
| 1236 | 84 | primary | NK cell | T cell | 1.12 | 3.90E-02 | 9.62E-01 | 3.90E-02 | 0 |
| 1237 | 84 | primary | NPC | AC | 0.00 | 1.00E+00 | 9.77E-01 | 9.77E-01 | 0 |
| 1238 | 84 | primary | NPC | Astrocyte | 2.54 | 4.96E-01 | 5.31E-01 | 4.96E-01 | 0 |
| 1239 | 84 | primary | NPC | Endothelial | 1.00 | 9.99E-01 | 3.09E-01 | 3.09E-01 | 0 |
| 1240 | 84 | primary | NPC | MES | 2.83 | 7.59E-02 | 9.32E-01 | 7.59E-02 | 0 |
| 1241 | 84 | primary | NPC | Macrophage | 1.00 | 9.65E-01 | 7.68E-01 | 7.68E-01 | 0 |
| 1242 | 84 | primary | NPC | Microglia | 1.00 | 8.83E-01 | 8.72E-01 | 8.72E-01 | 0 |
| 1243 | 84 | primary | NPC | NK cell | 1.00 | 9.69E-01 | 7.28E-01 | 7.28E-01 | 0 |
| 1244 | 84 | primary | NPC | NPC | 0.00 | 1.00E+00 | 9.54E-01 | 9.54E-01 | 0 |
| 1245 | 84 | primary | NPC | Neuron | 0.00 | 1.00E+00 | 4.77E-01 | 4.77E-01 | 0 |
| 1246 | 84 | primary | NPC | OPC | 0.00 | 1.00E+00 | 8.71E-01 | 8.71E-01 | 0 |
| 1247 | 84 | primary | NPC | Oligodendrocyte | | | | | |
| 1248 | 84 | primary | NPC | T cell | 0.00 | 1.00E+00 | 2.10E-01 | 2.10E-01 | 0 |
| 1249 | 84 | primary | Neuron | AC | 0.00 | 1.00E+00 | 8.60E-01 | 8.60E-01 | 0 |
| 1250 | 84 | primary | Neuron | Astrocyte | 4.20 | 9.99E-04 | 1.00E+00 | 9.99E-04 | 1 |
| 1251 | 84 | primary | Neuron | Endothelial | 1.44 | 2.20E-02 | 9.80E-01 | 2.20E-02 | 0 |
| 1252 | 84 | primary | Neuron | MES | 2.12 | 9.75E-01 | 2.60E-02 | 2.60E-02 | 0 |
| 1253 | 84 | primary | Neuron | Macrophage | 1.04 | 8.17E-01 | 1.84E-01 | 1.84E-01 | 0 |
| 1254 | 84 | primary | Neuron | Microglia | 1.00 | 1.00E+00 | 4.24E-01 | 4.24E-01 | 0 |
| 1255 | 84 | primary | Neuron | NK cell | 1.00 | 1.00E+00 | 1.66E-01 | 1.66E-01 | 0 |
| 1256 | 84 | primary | Neuron | NPC | 0.00 | 1.00E+00 | 4.77E-01 | 4.77E-01 | 0 |
| 1257 | 84 | primary | Neuron | Neuron | 1.19 | 3.80E-02 | 9.63E-01 | 3.80E-02 | 0 |
| 1258 | 84 | primary | Neuron | OPC | 1.00 | 5.80E-01 | 9.98E-01 | 5.80E-01 | 0 |
| 1259 | 84 | primary | Neuron | Oligodendrocyte | | | | | |
| 1260 | 84 | primary | Neuron | T cell | 1.00 | 1.00E+00 | 6.40E-01 | 6.40E-01 | 0 |
| 1261 | 84 | primary | OPC | AC | 0.00 | 1.00E+00 | 9.69E-01 | 9.69E-01 | 0 |
| 1262 | 84 | primary | OPC | Astrocyte | 2.81 | 1.63E-01 | 8.43E-01 | 1.63E-01 | 0 |
| 1263 | 84 | primary | OPC | Endothelial | 1.00 | 1.00E+00 | 1.79E-01 | 1.79E-01 | 0 |
| 1264 | 84 | primary | OPC | MES | 3.08 | 9.99E-03 | 9.91E-01 | 9.99E-03 | 1 |
| 1265 | 84 | primary | OPC | Macrophage | 1.00 | 9.91E-01 | 6.71E-01 | 6.71E-01 | 0 |
| 1266 | 84 | primary | OPC | Microglia | 0.00 | 1.00E+00 | 3.40E-02 | 3.40E-02 | 0 |
| 1267 | 84 | primary | OPC | NK cell | 2.00 | 1.30E-02 | 9.99E-01 | 1.30E-02 | 0 |
| 1268 | 84 | primary | OPC | NPC | 0.00 | 1.00E+00 | 8.71E-01 | 8.71E-01 | 0 |
| 1269 | 84 | primary | OPC | Neuron | 1.33 | 1.60E-02 | 9.86E-01 | 1.60E-02 | 0 |
| 1270 | 84 | primary | OPC | OPC | 1.00 | 1.01E-01 | 9.99E-01 | 1.01E-01 | 0 |
| 1271 | 84 | primary | OPC | Oligodendrocyte | | | | | |
| 1272 | 84 | primary | OPC | T cell | 1.00 | 8.67E-01 | 9.12E-01 | 8.67E-01 | 0 |
| 1273 | 84 | primary | Oligodendrocyte | AC | | | | | |
| 1274 | 84 | primary | Oligodendrocyte | Astrocyte | | | | | |
| 1275 | 84 | primary | Oligodendrocyte | Endothelial | | | | | |
| 1276 | 84 | primary | Oligodendrocyte | MES | | | | | |
| 1277 | 84 | primary | Oligodendrocyte | Macrophage | | | | | |
| 1278 | 84 | primary | Oligodendrocyte | Microglia | | | | | |
| 1279 | 84 | primary | Oligodendrocyte | NK cell | | | | | |
| 1280 | 84 | primary | Oligodendrocyte | NPC | | | | | |
| 1281 | 84 | primary | Oligodendrocyte | Neuron | | | | | |
| 1282 | 84 | primary | Oligodendrocyte | OPC | | | | | |
| 1283 | 84 | primary | Oligodendrocyte | Oligodendrocyte | | | | | |
| 1284 | 84 | primary | Oligodendrocyte | T cell | | | | | |
| 1285 | 84 | primary | T cell | AC | 0.00 | 1.00E+00 | 6.97E-01 | 6.97E-01 | 0 |
| 1286 | 84 | primary | T cell | Astrocyte | 2.48 | 8.18E-01 | 1.83E-01 | 1.83E-01 | 0 |
| 1287 | 84 | primary | T cell | Endothelial | 1.47 | 9.99E-04 | 1.00E+00 | 9.99E-04 | 1 |
| 1288 | 84 | primary | T cell | MES | 3.07 | 9.99E-04 | 1.00E+00 | 9.99E-04 | 1 |
| 1289 | 84 | primary | T cell | Macrophage | 1.44 | 9.99E-04 | 1.00E+00 | 9.99E-04 | 1 |
| 1290 | 84 | primary | T cell | Microglia | 1.20 | 1.20E-02 | 9.89E-01 | 1.20E-02 | 0 |
| 1291 | 84 | primary | T cell | NK cell | 1.68 | 9.99E-04 | 1.00E+00 | 9.99E-04 | 1 |
| 1292 | 84 | primary | T cell | NPC | 0.00 | 1.00E+00 | 2.10E-01 | 2.10E-01 | 0 |
| 1293 | 84 | primary | T cell | Neuron | 1.00 | 1.00E+00 | 8.46E-01 | 8.46E-01 | 0 |
| 1294 | 84 | primary | T cell | OPC | 1.00 | 8.67E-01 | 9.92E-01 | 8.67E-01 | 0 |
| 1295 | 84 | primary | T cell | Oligodendrocyte | | | | | |
| 1296 | 84 | primary | T cell | T cell | 1.23 | 2.30E-02 | 9.81E-01 | 2.30E-02 | 0 |
| 1297 | 84 | recurrent | AC | AC | 0.00 | 1.00E+00 | 1.00E+00 | 1.00E+00 | 0 |
| 1298 | 84 | recurrent | AC | Astrocyte | 0.00 | 1.00E+00 | 3.40E-02 | 3.40E-02 | 0 |
| 1299 | 84 | recurrent | AC | Endothelial | 0.00 | 1.00E+00 | 8.52E-01 | 8.52E-01 | 0 |
| 1300 | 84 | recurrent | AC | MES | 5.00 | 9.99E-03 | 9.99E-01 | 9.99E-03 | 1 |
| 1301 | 84 | recurrent | AC | Macrophage | 0.00 | 1.00E+00 | 8.24E-01 | 8.24E-01 | 0 |
| 1302 | 84 | recurrent | AC | Microglia | 0.00 | 1.00E+00 | 8.91E-01 | 8.91E-01 | 0 |
| 1303 | 84 | recurrent | AC | NK cell | 0.00 | 1.00E+00 | 8.14E-01 | 8.14E-01 | 0 |
| 1304 | 84 | recurrent | AC | NPC | | | | | |
| 1305 | 84 | recurrent | AC | Neuron | 0.00 | 1.00E+00 | 9.48E-01 | 9.48E-01 | 0 |
| 1306 | 84 | recurrent | AC | OPC | | | | | |
| 1307 | 84 | recurrent | AC | Oligodendrocyte | 0.00 | 1.00E+00 | 6.00E-01 | 6.00E-01 | 0 |
| 1308 | 84 | recurrent | AC | T cell | 1.00 | 6.99E-02 | 9.97E-01 | 6.99E-02 | 0 |
| 1309 | 84 | recurrent | Astrocyte | AC | 0.00 | 1.00E+00 | 3.40E-02 | 3.40E-02 | 0 |
| 1310 | 84 | recurrent | Astrocyte | Astrocyte | 4.05 | 9.99E-04 | 1.00E+00 | 9.99E-04 | 1 |
| 1311 | 84 | recurrent | Astrocyte | Endothelial | 1.26 | 9.99E-04 | 1.00E+00 | 9.99E-04 | 1 |
| 1312 | 84 | recurrent | Astrocyte | MES | 1.99 | 9.99E-04 | 1.00E+00 | 9.99E-04 | 1 |
| 1313 | 84 | recurrent | Astrocyte | Macrophage | 1.12 | 2.00E-03 | 9.99E-01 | 2.00E-03 | 1 |
| 1314 | 84 | recurrent | Astrocyte | Microglia | 1.08 | 2.80E-02 | 9.73E-01 | 2.80E-02 | 0 |
| 1315 | 84 | recurrent | Astrocyte | NK cell | 1.13 | 6.99E-03 | 9.94E-01 | 6.99E-03 | 1 |
| 1316 | 84 | recurrent | Astrocyte | NPC | | | | | |
| 1317 | 84 | recurrent | Astrocyte | Neuron | 1.12 | 9.99E-04 | 1.00E+00 | 9.99E-04 | 1 |
| 1318 | 84 | recurrent | Astrocyte | OPC | | | | | |
| 1319 | 84 | recurrent | Astrocyte | Oligodendrocyte | 1.54 | 9.99E-04 | 1.00E+00 | 9.99E-04 | 1 |

| row | patient | surgery | from cell type | to cell tye | observed count | permutations (greater than) | permutations (less than) | p value | interaction significance/direction |
|---|---|---|---|---|---|---|---|---|---|
| 1320 | 84 | recurrent | Astrocyte | T cell | 1.00 | 1.00E+00 | 9.99E-04 | 9.99E-04 | -1 |
| 1321 | 84 | recurrent | Endothelial | AC | 0.00 | 1.00E+00 | 8.52E-01 | 8.52E-01 | 0 |
| 1322 | 84 | recurrent | Endothelial | Astrocyte | 2.53 | 1.00E+00 | 9.99E-04 | 9.99E-04 | -1 |
| 1323 | 84 | recurrent | Endothelial | Endothelial | 2.32 | 9.99E-04 | 1.00E+00 | 9.99E-04 | 1 |
| 1324 | 84 | recurrent | Endothelial | MES | 1.93 | 5.01E-01 | 5.01E-01 | 5.01E-01 | 0 |
| 1325 | 84 | recurrent | Endothelial | Macrophage | 1.35 | 9.99E-04 | 1.00E+00 | 9.99E-04 | 1 |
| 1326 | 84 | recurrent | Endothelial | Microglia | 1.15 | 6.49E-02 | 9.37E-01 | 6.49E-02 | 0 |
| 1327 | 84 | recurrent | Endothelial | NK cell | 1.37 | 9.99E-04 | 1.00E+00 | 9.99E-04 | 1 |
| 1328 | 84 | recurrent | Endothelial | NPC | | | | | |
| 1329 | 84 | recurrent | Endothelial | Neuron | 1.00 | 1.00E+00 | 8.28E-01 | 8.28E-01 | 0 |
| 1330 | 84 | recurrent | Endothelial | OPC | | | | | |
| 1331 | 84 | recurrent | Endothelial | Oligodendrocyte | 1.38 | 3.20E-02 | 9.69E-01 | 3.20E-02 | 0 |
| 1332 | 84 | recurrent | Endothelial | T cell | 1.13 | 6.89E-02 | 9.52E-01 | 6.89E-02 | 0 |
| 1333 | 84 | recurrent | MES | AC | 1.00 | 8.32E-01 | 1.00E+00 | 8.32E-01 | 0 |
| 1334 | 84 | recurrent | MES | Astrocyte | 2.70 | 1.00E+00 | 9.99E-04 | 9.99E-04 | -1 |
| 1335 | 84 | recurrent | MES | Endothelial | 1.30 | 9.99E-04 | 1.00E+00 | 9.99E-04 | 1 |
| 1336 | 84 | recurrent | MES | MES | 2.91 | 9.99E-04 | 1.00E+00 | 9.99E-04 | 1 |
| 1337 | 84 | recurrent | MES | Macrophage | 1.22 | 9.99E-04 | 1.00E+00 | 9.99E-04 | 1 |
| 1338 | 84 | recurrent | MES | Microglia | 1.11 | 2.00E-03 | 9.99E-01 | 2.00E-03 | 1 |
| 1339 | 84 | recurrent | MES | NK cell | 1.26 | 9.99E-04 | 1.00E+00 | 9.99E-04 | 1 |
| 1340 | 84 | recurrent | MES | NPC | | | | | |
| 1341 | 84 | recurrent | MES | Neuron | 1.10 | 2.00E-03 | 9.99E-01 | 2.00E-03 | 1 |
| 1342 | 84 | recurrent | MES | OPC | | | | | |
| 1343 | 84 | recurrent | MES | Oligodendrocyte | 1.70 | 9.99E-04 | 1.00E+00 | 9.99E-04 | 1 |
| 1344 | 84 | recurrent | MES | T cell | 1.18 | 9.99E-04 | 1.00E+00 | 9.99E-04 | 1 |
| 1345 | 84 | recurrent | Macrophage | AC | 0.00 | 1.00E+00 | 8.24E-01 | 8.24E-01 | 0 |
| 1346 | 84 | recurrent | Macrophage | Astrocyte | 3.45 | 9.99E-04 | 1.00E+00 | 9.99E-04 | 1 |
| 1347 | 84 | recurrent | Macrophage | Endothelial | 1.53 | 9.99E-04 | 1.00E+00 | 9.99E-04 | 1 |
| 1348 | 84 | recurrent | Macrophage | MES | 2.35 | 9.99E-04 | 1.00E+00 | 9.99E-04 | 1 |
| 1349 | 84 | recurrent | Macrophage | Macrophage | 1.45 | 9.99E-04 | 1.00E+00 | 9.99E-04 | 1 |
| 1350 | 84 | recurrent | Macrophage | Microglia | 1.15 | 5.39E-02 | 9.53E-01 | 5.39E-02 | 0 |
| 1351 | 84 | recurrent | Macrophage | NK cell | 1.29 | 3.00E-03 | 9.98E-01 | 3.00E-03 | 1 |
| 1352 | 84 | recurrent | Macrophage | NPC | | | | | |
| 1353 | 84 | recurrent | Macrophage | Neuron | 1.00 | 1.00E+00 | 7.94E-01 | 7.94E-01 | 0 |
| 1354 | 84 | recurrent | Macrophage | OPC | | | | | |
| 1355 | 84 | recurrent | Macrophage | Oligodendrocyte | 1.36 | 7.39E-02 | 9.29E-01 | 7.39E-02 | 0 |
| 1356 | 84 | recurrent | Macrophage | T cell | 1.33 | 9.99E-04 | 1.00E+00 | 9.99E-04 | 1 |
| 1357 | 84 | recurrent | Microglia | AC | 0.00 | 1.00E+00 | 8.91E-01 | 8.91E-01 | 0 |
| 1358 | 84 | recurrent | Microglia | Astrocyte | 3.18 | 7.99E-03 | 9.93E-01 | 7.99E-03 | 1 |
| 1359 | 84 | recurrent | Microglia | Endothelial | 1.50 | 9.99E-04 | 1.00E+00 | 9.99E-04 | 1 |
| 1360 | 84 | recurrent | Microglia | MES | 2.38 | 9.99E-04 | 1.00E+00 | 9.99E-04 | 1 |
| 1361 | 84 | recurrent | Microglia | Macrophage | 1.12 | 2.40E-01 | 7.61E-01 | 2.40E-01 | 0 |
| 1362 | 84 | recurrent | Microglia | Microglia | 1.35 | 8.99E-03 | 9.92E-01 | 8.99E-03 | 1 |
| 1363 | 84 | recurrent | Microglia | NK cell | 1.38 | 2.00E-03 | 9.99E-01 | 2.00E-03 | 1 |
| 1364 | 84 | recurrent | Microglia | NPC | | | | | |
| 1365 | 84 | recurrent | Microglia | Neuron | 1.00 | 9.99E-01 | 8.79E-01 | 8.79E-01 | 0 |
| 1366 | 84 | recurrent | Microglia | OPC | | | | | |
| 1367 | 84 | recurrent | Microglia | Oligodendrocyte | 1.25 | 5.75E-01 | 4.58E-01 | 4.58E-01 | 0 |
| 1368 | 84 | recurrent | Microglia | T cell | 1.31 | 6.99E-03 | 9.94E-01 | 6.99E-03 | 1 |
| 1369 | 84 | recurrent | NK cell | AC | 0.00 | 1.00E+00 | 8.14E-01 | 8.14E-01 | 0 |
| 1370 | 84 | recurrent | NK cell | Astrocyte | 3.16 | 2.00E-03 | 9.99E-01 | 2.00E-03 | 1 |
| 1371 | 84 | recurrent | NK cell | Endothelial | 1.76 | 9.99E-04 | 1.00E+00 | 9.99E-04 | 1 |
| 1372 | 84 | recurrent | NK cell | MES | 2.39 | 9.99E-04 | 1.00E+00 | 9.99E-04 | 1 |
| 1373 | 84 | recurrent | NK cell | Macrophage | 1.31 | 2.00E-03 | 9.99E-01 | 2.00E-03 | 1 |
| 1374 | 84 | recurrent | NK cell | Microglia | 1.31 | 9.99E-04 | 1.00E+00 | 9.99E-04 | 1 |
| 1375 | 84 | recurrent | NK cell | NK cell | 1.68 | 9.99E-04 | 1.00E+00 | 9.99E-04 | 1 |
| 1376 | 84 | recurrent | NK cell | NPC | | | | | |
| 1377 | 84 | recurrent | NK cell | Neuron | 1.11 | 7.69E-02 | 9.46E-01 | 7.69E-02 | 0 |
| 1378 | 84 | recurrent | NK cell | OPC | | | | | |
| 1379 | 84 | recurrent | NK cell | Oligodendrocyte | 1.59 | 9.99E-04 | 1.00E+00 | 9.99E-04 | 1 |
| 1380 | 84 | recurrent | NK cell | T cell | 1.57 | 9.99E-04 | 1.00E+00 | 9.99E-04 | 1 |
| 1381 | 84 | recurrent | NPC | AC | | | | | |
| 1382 | 84 | recurrent | NPC | Astrocyte | | | | | |
| 1383 | 84 | recurrent | NPC | Endothelial | | | | | |
| 1384 | 84 | recurrent | NPC | MES | | | | | |
| 1385 | 84 | recurrent | NPC | Macrophage | | | | | |
| 1386 | 84 | recurrent | NPC | Microglia | | | | | |
| 1387 | 84 | recurrent | NPC | NK cell | | | | | |
| 1388 | 84 | recurrent | NPC | NPC | | | | | |
| 1389 | 84 | recurrent | NPC | Neuron | | | | | |
| 1390 | 84 | recurrent | NPC | OPC | | | | | |
| 1391 | 84 | recurrent | NPC | Oligodendrocyte | | | | | |
| 1392 | 84 | recurrent | NPC | T cell | | | | | |
| 1393 | 84 | recurrent | Neuron | AC | 0.00 | 1.00E+00 | 9.48E-01 | 9.48E-01 | 0 |
| 1394 | 84 | recurrent | Neuron | Astrocyte | 3.51 | 9.99E-04 | 1.00E+00 | 9.99E-04 | 1 |
| 1395 | 84 | recurrent | Neuron | Endothelial | 1.50 | 5.00E-03 | 9.99E-01 | 5.00E-03 | 1 |
| 1396 | 84 | recurrent | Neuron | MES | 2.94 | 9.99E-04 | 1.00E+00 | 9.99E-04 | 1 |
| 1397 | 84 | recurrent | Neuron | Macrophage | 1.10 | 4.01E-01 | 6.45E-01 | 4.01E-01 | 0 |
| 1398 | 84 | recurrent | Neuron | Microglia | 1.00 | 9.99E-01 | 7.28E-01 | 7.28E-01 | 0 |
| 1399 | 84 | recurrent | Neuron | NK cell | 1.25 | 9.49E-02 | 9.40E-01 | 9.49E-02 | 0 |
| 1400 | 84 | recurrent | Neuron | NPC | | | | | |
| 1401 | 84 | recurrent | Neuron | Neuron | 1.17 | 2.70E-02 | 9.74E-01 | 2.70E-02 | 0 |
| 1402 | 84 | recurrent | Neuron | OPC | | | | | |
| 1403 | 84 | recurrent | Neuron | Oligodendrocyte | 1.56 | 1.20E-02 | 9.90E-01 | 1.20E-02 | 0 |
| 1404 | 84 | recurrent | Neuron | T cell | 1.25 | 7.49E-02 | 9.50E-01 | 7.49E-02 | 0 |
| 1405 | 84 | recurrent | OPC | AC | | | | | |
| 1406 | 84 | recurrent | OPC | Astrocyte | | | | | |
| 1407 | 84 | recurrent | OPC | Endothelial | | | | | |
| 1408 | 84 | recurrent | OPC | MES | | | | | |
| 1409 | 84 | recurrent | OPC | Macrophage | | | | | |
| 1410 | 84 | recurrent | OPC | Microglia | | | | | |
| 1411 | 84 | recurrent | OPC | NK cell | | | | | |
| 1412 | 84 | recurrent | OPC | NPC | | | | | |
| 1413 | 84 | recurrent | OPC | Neuron | | | | | |
| 1414 | 84 | recurrent | OPC | OPC | | | | | |
| 1415 | 84 | recurrent | OPC | Oligodendrocyte | | | | | |
| 1416 | 84 | recurrent | OPC | T cell | | | | | |
| 1417 | 84 | recurrent | Oligodendrocyte | AC | 0.00 | 1.00E+00 | 6.00E-01 | 6.00E-01 | 0 |
| 1418 | 84 | recurrent | Oligodendrocyte | Astrocyte | 2.44 | 1.00E+00 | 9.99E-04 | 9.99E-04 | -1 |
| 1419 | 84 | recurrent | Oligodendrocyte | Endothelial | 1.30 | 9.99E-04 | 1.00E+00 | 9.99E-04 | 1 |
| 1420 | 84 | recurrent | Oligodendrocyte | MES | 2.14 | 9.99E-04 | 1.00E+00 | 9.99E-04 | 1 |
| 1421 | 84 | recurrent | Oligodendrocyte | Macrophage | 1.06 | 7.58E-01 | 2.56E-01 | 2.56E-01 | 0 |
| 1422 | 84 | recurrent | Oligodendrocyte | Microglia | 1.14 | 9.99E-03 | 9.92E-01 | 9.99E-03 | 1 |
| 1423 | 84 | recurrent | Oligodendrocyte | NK cell | 1.20 | 9.99E-04 | 1.00E+00 | 9.99E-04 | 1 |
| 1424 | 84 | recurrent | Oligodendrocyte | NPC | | | | | |
| 1425 | 84 | recurrent | Oligodendrocyte | Neuron | 1.00 | 1.00E+00 | 5.58E-01 | 5.58E-01 | 0 |
| 1426 | 84 | recurrent | Oligodendrocyte | OPC | | | | | |
| 1427 | 84 | recurrent | Oligodendrocyte | Oligodendrocyte | 2.78 | 9.99E-04 | 1.00E+00 | 9.99E-04 | 1 |
| 1428 | 84 | recurrent | Oligodendrocyte | T cell | 1.42 | 9.99E-04 | 1.00E+00 | 9.99E-04 | 1 |
| 1429 | 84 | recurrent | T cell | AC | 1.00 | 6.99E-02 | 1.00E+00 | 6.99E-02 | 0 |
| 1430 | 84 | recurrent | T cell | Astrocyte | 2.93 | 3.85E-01 | 6.16E-01 | 3.85E-01 | 0 |

| row | patient | surgery | from cell type | to cell tye | observed count | permutations (greater than) | permutations (less than) | p value | interaction significance/direction |
|---|---|---|---|---|---|---|---|---|---|
| 1431 | 84 | recurrent | T cell | Endothelial | 1.80 | 9.99E-04 | 1.00E+00 | 9.99E-04 | 1 |
| 1432 | 84 | recurrent | T cell | MES | 2.50 | 9.99E-04 | 1.00E+00 | 9.99E-04 | 1 |
| 1433 | 84 | recurrent | T cell | Macrophage | 1.12 | 2.44E-01 | 7.59E-01 | 2.44E-01 | 0 |
| 1434 | 84 | recurrent | T cell | Microglia | 1.35 | 6.99E-03 | 9.94E-01 | 6.99E-03 | 1 |
| 1435 | 84 | recurrent | T cell | NK cell | 1.42 | 3.00E-03 | 9.98E-01 | 3.00E-03 | 1 |
| 1436 | 84 | recurrent | T cell | NPC | | | | | |
| 1437 | 84 | recurrent | T cell | Neuron | 1.00 | 9.88E-01 | 9.14E-01 | 9.14E-01 | 0 |
| 1438 | 84 | recurrent | T cell | OPC | | | | | |
| 1439 | 84 | recurrent | T cell | Oligodendrocyte | 1.56 | 3.00E-03 | 9.98E-01 | 3.00E-03 | 1 |
| 1440 | 84 | recurrent | T cell | T cell | 1.59 | 3.00E-03 | 9.98E-01 | 3.00E-03 | 1 |

**Supplementary Table 10. Summarised pair-wise cell-cell interactions compared to a null model of spatial randomness.** Statistical significance and direction of the interactions were determined using a permutation test, with p-values indicating interactions more or less likely than random. The interaction significance/direction denotes the strength and direction of all the pair-wise interactions: >=1 (significant attraction interactions); <= -1 (significant avoidance interactions); 0 (neutral and/or non-statistically significant interactions). When an interaction type is significant across both primary and recurrent surgeries, both surgery-specific p values are listed.

| row | from cell | to cell | significant interaction type | present across (surgery) | interaction significance | primary p value | recurrent p value |
|---|---|---|---|---|---|---|---|
| 1 | AC | AC | Interacting | Primary | 3 | 9.99E-04 | |
| 2 | AC | Astrocyte | | | 0 | | |
| 3 | AC | Endothelial | | | 2 | | |
| 4 | AC | MES | | | -1 | | |
| 5 | AC | Macrophage | | | 1 | | |
| 6 | AC | Microglia | | | 0 | | |
| 7 | AC | NK cell | | | 0 | | |
| 8 | AC | NPC | | | 1 | | |
| 9 | AC | Neuron | | | 0 | | |
| 10 | AC | OPC | | | 2 | | |
| 11 | AC | Oligodendrocyte | | | 1 | | |
| 12 | AC | T cell | | | 1 | | |
| 13 | Astrocyte | AC | | | 2 | | |
| 14 | Astrocyte | Astrocyte | Interacting | Both | 3 | 9.99E-04 | 9.99E-04 |
| 15 | Astrocyte | Endothelial | Interacting | Recurrent | 2 | | 9.99E-04 |
| 16 | Astrocyte | MES | Interacting | Recurrent | 2 | | 9.99E-04 |
| 17 | Astrocyte | Macrophage | Interacting | Recurrent | 1 | | 2.00E-03 |
| 18 | Astrocyte | Microglia | | | 2 | | |
| 19 | Astrocyte | NK cell | Interacting | Recurrent | 0 | | 9.99E-04 |
| 20 | Astrocyte | NPC | Interacting | Recurrent | 1 | | 9.99E-04 |
| 21 | Astrocyte | Neuron | | | 2 | | |
| 22 | Astrocyte | OPC | Interacting | Recurrent | 2 | | 9.99E-04 |
| 23 | Astrocyte | Oligodendrocyte | Interacting | Recurrent | 0 | | 9.99E-04 |
| 24 | Astrocyte | T cell | | | 1 | | |
| 25 | Endothelial | AC | | | 2 | | |
| 26 | Endothelial | Astrocyte | Interacting | Primary | 3 | 9.99E-04 | |
| 27 | Endothelial | Endothelial | Interacting | Both | 5 | 9.99E-04 | 9.99E-04 |
| 28 | Endothelial | MES | Interacting | Primary | 4 | 9.99E-04 | |
| 29 | Endothelial | Macrophage | Interacting | Recurrent | 0 | | 9.99E-04 |
| 30 | Endothelial | Microglia | Interacting | Primary | 3 | 9.99E-04 | |
| 31 | Endothelial | NK cell | Interacting | Both | 4 | 9.99E-04 | 9.99E-04 |
| 32 | Endothelial | NPC | | | 0 | | |
| 33 | Endothelial | Neuron | | | 1 | | |
| 34 | Endothelial | OPC | | | 2 | | |
| 35 | Endothelial | Oligodendrocyte | | | 1 | | |
| 36 | Endothelial | T cell | | | 2 | | |
| 37 | MES | AC | | | -1 | | |
| 38 | MES | Astrocyte | | | 2 | | |
| 39 | MES | Endothelial | Interacting | Both | 3 | 9.99E-04 | 9.99E-04 |
| 40 | MES | MES | Interacting | Both | 5 | 9.99E-04 | 9.99E-04 |
| 41 | MES | Macrophage | Interacting | Recurrent | 1 | | 9.99E-04 |
| 42 | MES | Microglia | | | 2 | | |
| 43 | MES | NK cell | Interacting | Both | 3 | 9.99E-04 | 9.99E-04 |
| 44 | MES | NPC | | | -2 | | |
| 45 | MES | Neuron | | | 0 | | |
| 46 | MES | OPC | | | -2 | | |
| 47 | MES | Oligodendrocyte | Interacting | Recurrent | 1 | | 9.99E-04 |
| 48 | MES | T cell | Interacting | Primary | 4 | 9.99E-04 | |
| 49 | Macrophage | AC | Interacting | Primary | 3 | 9.99E-04 | |
| 50 | Macrophage | Astrocyte | Interacting | Primary | 4 | 9.99E-04 | |
| 51 | Macrophage | Endothelial | Interacting | Both | 5 | 9.99E-04 | 9.99E-04 |
| 52 | Macrophage | MES | Interacting | Both | 3 | 9.99E-04 | 9.99E-04 |
| 53 | Macrophage | Macrophage | Interacting | Both | 3 | 9.99E-04 | 9.99E-04 |
| 54 | Macrophage | Microglia | Interacting | Both | 5 | 9.99E-04 | 9.99E-04 |
| 55 | Macrophage | NK cell | Interacting | Both | 5 | 9.99E-04 | 9.99E-04 |
| 56 | Macrophage | NPC | | | 2 | | |
| 57 | Macrophage | Neuron | Interacting | Primary | 3 | 9.99E-04 | |
| 58 | Macrophage | OPC | | | 1 | | |
| 59 | Macrophage | Oligodendrocyte | Interacting | Recurrent | 2 | | 9.99E-04 |
| 60 | Macrophage | T cell | Interacting | Recurrent | 2 | | 9.99E-04 |
| 61 | Microglia | AC | | | 2 | | |
| 62 | Microglia | Astrocyte | Interacting | Primary | 4 | 9.99E-04 | |
| 63 | Microglia | Endothelial | Interacting | Both | 5 | 9.99E-04 | 9.99E-04 |
| 64 | Microglia | MES | Interacting | Both | 4 | 9.99E-04 | 9.99E-04 |
| 65 | Microglia | Macrophage | Interacting | Both | 3 | 9.99E-04 | 9.99E-04 |
| 66 | Microglia | Microglia | Interacting | Both | 4 | 9.99E-04 | 9.99E-04 |
| 67 | Microglia | NK cell | Interacting | Recurrent | 2 | | 9.99E-04 |
| 68 | Microglia | NPC | Interacting | Recurrent | 1 | | 9.99E-04 |
| 69 | Microglia | Neuron | | | 0 | | |
| 70 | Microglia | OPC | | | 2 | | |
| 71 | Microglia | Oligodendrocyte | Interacting | Recurrent | 2 | | 9.99E-04 |
| 72 | Microglia | T cell | Interacting | Both | 4 | 9.99E-04 | 9.99E-04 |

| row | from cell | to cell | significant interaction type | present across (surgery) | interaction significance | primary p value | recurrent p value |
|---|---|---|---|---|---|---|---|
| 73 | NK cell | AC | | | 2 | | |
| 74 | NK cell | Astrocyte | | | 0 | | |
| 75 | NK cell | Endothelial | Interacting | Both | 5 | 9.99E-04 | 9.99E-04 |
| 76 | NK cell | MES | | | 1 | | |
| 77 | NK cell | Macrophage | Interacting | Recurrent | 2 | | 9.99E-04 |
| 78 | NK cell | Microglia | Interacting | Both | 4 | 9.99E-04 | 9.99E-04 |
| 79 | NK cell | NK cell | Interacting | Both | 3 | 9.99E-04 | 9.99E-04 |
| 80 | NK cell | NPC | | | 2 | | |
| 81 | NK cell | Neuron | | | 1 | | |
| 82 | NK cell | OPC | | | 2 | | |
| 83 | NK cell | Oligodendrocyte | Interacting | Recurrent | 0 | | 9.99E-04 |
| 84 | NK cell | T cell | Interacting | Both | 3 | 9.99E-04 | 9.99E-04 |
| 85 | NPC | AC | | | 1 | | |
| 86 | NPC | Astrocyte | Interacting | Recurrent | 1 | | 9.99E-04 |
| 87 | NPC | Endothelial | | | 2 | | |
| 88 | NPC | MES | | | 0 | | |
| 89 | NPC | Macrophage | | | 1 | | |
| 90 | NPC | Microglia | | | 1 | | |
| 91 | NPC | NK cell | | | -1 | | |
| 92 | NPC | NPC | Interacting | Recurrent | 1 | | 9.99E-04 |
| 93 | NPC | Neuron | | | 2 | | |
| 94 | NPC | OPC | | | 2 | | |
| 95 | NPC | Oligodendrocyte | | | 1 | | |
| 96 | NPC | T cell | | | 0 | | |
| 97 | Neuron | AC | | | 2 | | |
| 98 | Neuron | Astrocyte | Interacting | Recurrent | 2 | 9.99E-04 | 9.99E-04 |
| 99 | Neuron | Endothelial | Interacting | Recurrent | 1 | 9.99E-04 | 9.99E-04 |
| 100 | Neuron | MES | Interacting | Recurrent | 2 | 9.99E-04 | 9.99E-04 |
| 101 | Neuron | Macrophage | | | 2 | | |
| 102 | Neuron | Microglia | | | 1 | | |
| 103 | Neuron | NK cell | | | -1 | | |
| 104 | Neuron | NPC | | | 2 | | |
| 105 | Neuron | Neuron | | | 2 | | |
| 106 | Neuron | OPC | | | 2 | | |
| 107 | Neuron | Oligodendrocyte | | | 1 | | |
| 108 | Neuron | T cell | | | 1 | | |
| 109 | OPC | AC | | | 1 | | |
| 110 | OPC | Astrocyte | | | 1 | | |
| 111 | OPC | Endothelial | | | 1 | | |
| 112 | OPC | MES | | | 1 | | |
| 113 | OPC | Macrophage | | | 1 | | |
| 114 | OPC | Microglia | | | 1 | | |
| 115 | OPC | NK cell | | | -1 | | |
| 116 | OPC | NPC | | | 1 | | |
| 117 | OPC | Neuron | | | 2 | | |
| 118 | OPC | OPC | | | 2 | | |
| 119 | OPC | Oligodendrocyte | | | 1 | | |
| 120 | OPC | T cell | | | 1 | | |
| 121 | Oligodendrocyte | AC | | | 1 | | |
| 122 | Oligodendrocyte | Astrocyte | | | 1 | | |
| 123 | Oligodendrocyte | Endothelial | Interacting | Recurrent | 2 | | 9.99E-04 |
| 124 | Oligodendrocyte | MES | Interacting | Primary | 3 | 9.99E-04 | |
| 125 | Oligodendrocyte | Macrophage | | | 2 | | |
| 126 | Oligodendrocyte | Microglia | Interacting | Recurrent | 2 | | 9.99E-04 |
| 127 | Oligodendrocyte | NK cell | | | -1 | | |
| 128 | Oligodendrocyte | NPC | | | 1 | | |
| 129 | Oligodendrocyte | Neuron | | | 1 | | |
| 130 | Oligodendrocyte | OPC | | | 1 | | |
| 131 | Oligodendrocyte | Oligodendrocyte | Interacting | Recurrent | 2 | | 9.99E-04 |
| 132 | Oligodendrocyte | T cell | Interacting | Recurrent | 2 | | 9.99E-04 |
| 133 | T cell | AC | | | 2 | | |
| 134 | T cell | Astrocyte | | | 1 | | |
| 135 | T cell | Endothelial | Interacting | Both | 5 | 9.99E-04 | 9.99E-04 |
| 136 | T cell | MES | Interacting | Both | 5 | 9.99E-04 | 9.99E-04 |
| 137 | T cell | Macrophage | | | 0 | | |
| 138 | T cell | Microglia | Interacting | Both | 4 | 9.99E-04 | 9.99E-04 |
| 139 | T cell | NK cell | Interacting | Both | 3 | 9.99E-04 | 9.99E-04 |
| 140 | T cell | NPC | | | 1 | | |
| 141 | T cell | Neuron | | | 1 | | |
| 142 | T cell | OPC | | | 2 | | |
| 143 | T cell | Oligodendrocyte | Interacting | Recurrent | 2 | | 9.99E-04 |
| 144 | T cell | T cell | Interacting | Both | 3 | 9.99E-04 | 9.99E-04 |

**Supplementary Table 11. Summary of defined cellular neighborhoods (CNs).** The table details how each CN aligns with Greenwald et al.'s metaprograms, and their relative proportions in primary and recurrent samples.

| cellular neighbourhood | Greenwald et.al layer | Greenwald et.al label | n cells (primary) | proportion (primary) | n cells (recurrent) | proportion (recurrent) |
|---|---|---|---|---|---|---|
| CN1 | 3 | Mac | 6943 | 12.4% | 26 | 0.1% |
| CN2 | 2 | MES-Ast | 2938 | 5.2% | 6143 | 16.1% |
| CN3 | 5 | Oligo | 3454 | 6.2% | 3009 | 7.9% |
| CN4 | 3 | T-cell | 6721 | 12.0% | 4418 | 11.6% |
| CN5 | 3 | Vasc | 5171 | 9.2% | 1560 | 4.1% |
| CN6 | 1 | MES-Hyp | 9186 | 16.4% | 2451 | 6.4% |
| CN7 | 5 | Neuron | 1469 | 2.6% | 5737 | 15.1% |
| CN8 | 4 | AC | 7961 | 14.2% | 887 | 2.3% |
| CN9 | 5 | Reactive-Ast/Neuron | 765 | 1.4% | 4931 | 12.9% |
| CN10 | 3 | T-cell | 3454 | 6.2% | 543 | 1.4% |
| CN11 | 5 | Reactive-Ast | 2725 | 4.9% | 7641 | 20.1% |
| CN12 | 2 | Inflammatory-Mac | 5179 | 9.3% | 746 | 2.0% |

**Supplementary Table 12. Comparison of cellular state (hypoxia and Epithelial to mesenchymal transition - EMT) protein marker abundance between primary and recurrent surgery samples and across each defined cellular neighbourhood (CN).** Statistical significance was assessed using an unpaired Wilcoxon test, with adjusted p-values calculated using the false discovery rate (FDR) method. The p value significance levels are denoted using the following symbols: **** ($p < 0.0001$); *** ($p < 0.001$); ** ($p < 0.01$); * ($p < 0.05$); n.s (not significant).

| Greenwald_et_al_label (CN) | cellular state | comparison groups | n cells (primary) | n cells (recurrent) | p value | adjusted p value | p significance |
|---|---|---|---|---|---|---|---|
| MES-Hyp (CN6) | hypoxia | Prim vs Rec | 6527 | 1407 | 3.70E-20 | 6.78E-20 | **** |
| MES-Ast (CN2) | hypoxia | Prim vs Rec | 748 | 811 | 1.52E-74 | 1.67E-73 | **** |
| Inflammatory-Mac (CN12) | hypoxia | Prim vs Rec | 777 | 71 | 8.52E-01 | 9.13E-01 | n.s |
| Mac (CN1) | hypoxia | Prim vs Rec | 262 | 2 | 6.77E-02 | 8.27E-02 | n.s |
| Vasc (CN5) | hypoxia | Prim vs Rec | 554 | 36 | 8.02E-03 | 1.10E-02 | ** |
| T-cell (CN4 & CN10) | hypoxia | Prim vs Rec | 2406 | 621 | 1.40E-30 | 3.08E-30 | **** |
| AC (CN8) | hypoxia | Prim vs Rec | 5258 | 633 | 6.80E-66 | 3.74E-65 | **** |
| Neuron (CN7) | hypoxia | Prim vs Rec | 311 | 310 | 9.13E-01 | 9.13E-01 | n.s |
| Reactive-Ast/Neuron (CN9) | hypoxia | Prim vs Rec | 106 | 225 | 1.07E-09 | 1.68E-09 | **** |
| Reactive-Ast (CN11) | hypoxia | Prim vs Rec | 248 | 478 | 1.36E-41 | 3.74E-41 | **** |
| Oligo (CN3) | hypoxia | Prim vs Rec | 873 | 440 | 4.17E-42 | 1.53E-41 | **** |
| MES-Hyp (CN6) | EMT | Prim vs Rec | 6527 | 1407 | 6.11E-01 | 6.12E-01 | n.s |
| MES-Ast (CN2) | EMT | Prim vs Rec | 748 | 811 | 2.69E-27 | 9.86E-27 | **** |
| Inflammatory-Mac (CN12) | EMT | Prim vs Rec | 777 | 71 | 1.43E-04 | 2.25E-04 | *** |
| Mac (CN1) | EMT | Prim vs Rec | 262 | 2 | 6.12E-01 | 6.12E-01 | n.s |
| Vasc (CN5) | EMT | Prim vs Rec | 554 | 36 | 4.34E-07 | 7.96E-07 | **** |
| T-cell (CN4 & CN10) | EMT | Prim vs Rec | 2406 | 621 | 2.82E-59 | 1.55E-58 | **** |
| AC (CN8) | EMT | Prim vs Rec | 5258 | 633 | 1.04E-195 | 1.14E-194 | **** |
| Neuron (CN7) | EMT | Prim vs Rec | 311 | 310 | 1.14E-01 | 1.39E-01 | n.s |
| Reactive-Ast/Neuron (CN9) | EMT | Prim vs Rec | 106 | 225 | 6.90E-04 | 9.49E-04 | *** |
| Reactive-Ast (CN11) | EMT | Prim vs Rec | 248 | 478 | 1.80E-11 | 3.96E-11 | **** |
| Oligo (CN3) | EMT | Prim vs Rec | 873 | 440 | 3.60E-22 | 9.90E-22 | **** |
